# Computation-through-Dynamics Toolkit: Simulated datasets and quality metrics for dynamical models of neural activity

**DOI:** 10.1101/2025.02.07.637062

**Authors:** Christopher Versteeg, Jonathan D. McCart, Mitchell Ostrow, David M. Zoltowski, Clayton B. Washington, Laura Driscoll, Olivier Codol, Jonathan A. Michaels, Scott W. Linderman, David Sussillo, Chethan Pandarinath

## Abstract

A primary goal of systems neuroscience is to discover how ensembles of neurons transform inputs into goal-directed behavior, a process known as neural computation. A powerful framework for understanding neural computation uses neural dynamics – the rules that govern how neural activity evolves over time – to explain how goal-directed input-output transformations occur. As dynamical rules are not directly observable, we need computational models that can infer neural dynamics from recorded neural activity. We typically validate such models using synthetic datasets with known ground-truth dynamics, but unfortunately existing synthetic datasets don’t reflect fundamental features of neural computation and may therefore be poor proxies for neural systems. Further, the field lacks validated metrics for quantifying the accuracy of the dynamics inferred by models. The Computation-through-Dynamics Toolkit (CtDToolkit) addresses these critical gaps by providing: 1) synthetic datasets that reflect computational properties of biological neural circuits, 2) interpretable metrics for quantifying model performance, and 3) a standardized pipeline for training and evaluating models with or without known external inputs. In this manuscript, we demonstrate how CtDToolkit can help guide the development, tuning, and troubleshooting of neural dynamics models. In summary, CtDToolkit provides a necessary framework for model developers to better understand and characterize neural computation through the lens of dynamics.

**Author Summary:** Understanding how the brain works requires interpretable accounts of how populations of neurons process information to produce behavior. One powerful approach is to study “neural dynamics”, the patterns of how neural activity evolves over time. Scientists develop computational models to infer these dynamics from neural recordings, but it has been challenging to know when the inferred dynamics are trustworthy. Existing datasets often lack key features of biological neural circuits, and current performance metrics can provide an incomplete picture of model quality.

We developed the Computation-through-Dynamics Toolkit (CtDToolkit) to solve these problems. Our toolkit provides three key resources: biologically motivated synthetic datasets, improved metrics that provide more holistic accounts of model performance, and a standardized workflow for training and evaluating models. We hope that CtDToolkit enables researchers to rigorously test, improve, and troubleshoot their models before applying them to real brain data. This work establishes a crucial foundation for developing better methods to understand neural computation, ultimately advancing our ability to decode how the brain transforms sensory information into thought and action.

## 1 Introduction

### 1.1 What is the computation-through-dynamics toolkit?

Understanding how the brain performs computation is the central aim of systems neuroscience. Although modern neural interfaces can now monitor hundreds or thousands of neurons simultaneously [11], we struggle to translate these massive new datasets into interpretable accounts of neural computation. We need a conceptual language that can describe how neural populations transform inputs into goal-directed behavior. Neural dynamics - the rules that govern how neural activity evolves over time - have gained renewed attention through advances in artificial neural network research [1, 3, 5, 15]. Neural dynamics offer a framework for connecting neural observations with neural computation [35].

A dynamics-based understanding of neural computation requires new methods that can estimate the (not directly observable) dynamics of neural circuits. Recent years have seen a surge in data-driven models that attempt to infer these dynamics by learning to reconstruct observed neural activity as the product of a model dynamical system [16, 21, 19, 25, 41, 36, 42, 66]. Unfortunately, we lack consensus on the types of synthetic systems and performance criteria that are appropriate for evaluating these models. A common set of datasets and metrics for model evaluation would facilitate model comparisons and help promising innovations be disseminated more quickly through the field.

Notably, the broader field of dynamical systems reconstruction (DSR) has long grappled with the analogous problem of validating an inferred dynamical system against a known ground-truth system, and has developed a mature set of evaluation measures for doing so on canonical (non-neural) systems such as the Lorenz and Rössler attractors [54, 45, 48]. However, the additional challenges presented by the dynamics implementing neural computation, chiefly the presence of substantial unknown external inputs and goal-directed task structure, require a framework that links these well-established methods to biological computational systems.

In this manuscript, we introduce the Computation-through-Dynamics Toolkit (CtDToolkit), a model development platform designed to help researchers assess the strengths and weaknesses of different data-driven (DD) dynamics models across a range of simulated neural datasets. As part of CtDToolkit, we present (1) a library of synthetic datasets that reflect goal-directed dynamical computations, (2) performance criteria and metrics designed to reveal specific model failures, and (3) a public codebase that allows researchers to submit new models, generate customized synthetic datasets, and iterate rapidly during model development. CtDToolkit is designed for modularity and extensibility, serving both as a shared framework for comparing DD models and as a flexible platform for tailoring datasets to specific experimental contexts.

This manuscript is divided into three sections: Section 1.2 lays out a theoretical foundation for the synthetic systems and performance metrics included in CtDToolkit, which Section 2 then describes in detail. Section 3 provides illustrative examples of how CtDToolkit datasets and metrics can help guide DD model development.

### 1.2 Computation-through-Dynamics: Definitions, Approach, and Challenges

#### 1.2.1 Problem Definition

Any satisfying account of neural computation needs to span three conceptual levels [9]: computational, algorithmic, and implementation (Fig. 1A). In this section, we provide a high-level description of each level and illustrative examples from a simple computational system, the 1-bit flip-flop.

**Figure 1.**
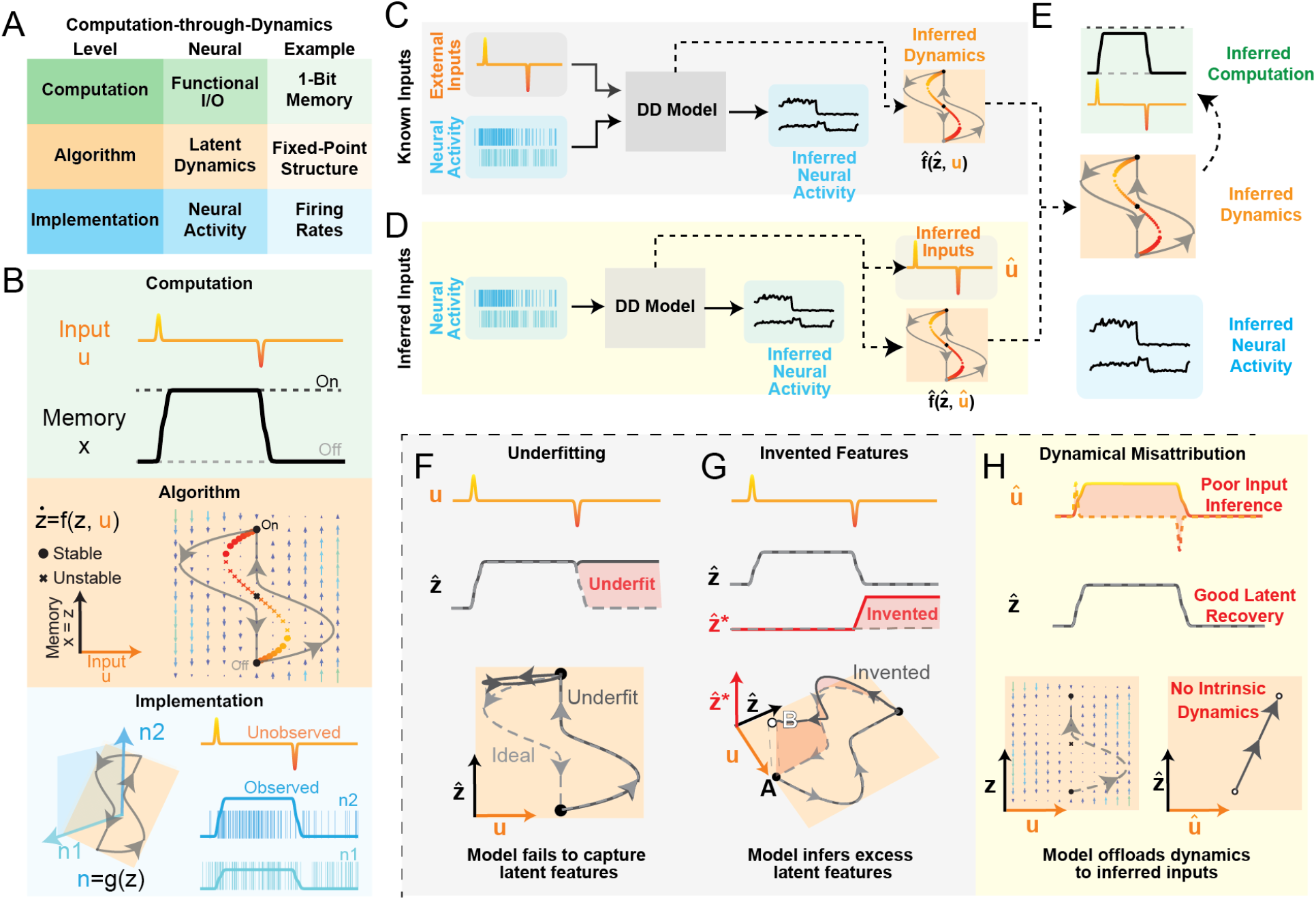
Computation-through-Dynamics framework and data-driven model failure modes. **A)** Overview of Computation-through-Dynamics framework with levels, corresponding concepts in neuroscience, and specific examples from 1-Bit Memory (1BFF) task. **B)** Schematic of conceptual hierarchy of 1-Bit Memory task. Top row (green): Computation – Example of input pulses **u** and desired output p for a single trial of 1BFF task. Middle row (orange): Algorithm – State-space diagram of dynamical system that performs 1BFF. x-axis: external input **u.**y-axis: latent state **z**. Flow field (written as **f**) controls how **z** evolves as a function of both **z** and **u**. circles: stable fixed points, : unstable fixed points. Output **x** is generated by projection **h** acting on the latent **z**. In this example **h** is the identity function. Bottom row (blue): Implementation of latent dynamics in neural activity. Left: Schematic of simple linear observation model of 1D latent dynamics into 2D neural activity. Inputs **u** are orthogonal to the linear observation model, implying that inputs are not directly visible in neural activity. Right: Simulated firing rates **n** and spiking activity **y** for two neurons in blue. **C)** Schematic of data-training pipeline (with known external inputs). Solid lines represent inputs and outputs from the DD model, dashed black lines represent what we hope our DD models infer indirectly. Neural observations (blue) and external inputs (orange) are provided to the dynamics model, which is optimized to reconstruct neural activity. **D)** Data-training pipeline (with inferred external inputs): Same as C, except the DD model does not receive external inputs and is instead tasked with inferring them. **E)** Our goal is to infer the computation performed by a neural circuit, but we only have access to observed neural activity. We hope that the dynamics 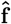 inferred by models in panels C,D match the true dynamics **f**, and will provide the basis for us to infer the computation performed by the circuit (green). **F-G)** Schematics of failure modes for models fit to 1BFF with supplied external inputs. **F)** Underfitting – Top row: True inputs **u** (orange/red) and true/inferred latent activity 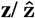 (solid/dashed, respectively). Red shaded area shows the activity that the underfit model fails to capture. Bottom row: State-space diagram of an ideal DD model (gray dashed line) and underfit DD model (black solid line). **G)** Invented Features – Top row: Same as F, except DD model has extra latent dimension 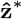, highlighted by a red shaded area. Bottom row: 3D state-space diagram of ideal DD model (gray dashed line) and DD model with invented features (solid black line). Ideal model and invented features model share fixed points (filled black circles, “A”), but the invented features model has an additional “invented” fixed point (white circle, “B”). This panel is a simplified, conceptual illustration of what an invented feature can look like for a short input sequence (here, a single pair of input pulses), and is not intended to depict the full dynamics under prolonged or repeated inputs. **H)** Schematic of the dynamical misattribution failure mode, which affects models that infer external inputs. Top row: Model-inferred inputs do not match the true inputs (highlighted by red shaded area), but accurately capture the latent activity without inventing features. Bottom row: State-space diagram with flow field for ideal DD model (left) and DD model with poor input accuracy (right). In this schematic, we only show the rising bit-flip for visual clarity. With inferred inputs, DD models are not obligated to learn any intrinsic dynamics.

##### What do we mean by “neural computation”?

The computation level is concerned with the goal a system is trying to accomplish. Because brains evolved to generate adaptive behavior, it is impossible to fully understand a neural circuit without reference to a computational goal. We loosely formalize a neural computation as a mapping from inputs **u** to outputs **x** tuned to accomplish a specific behaviorally-relevant goal (e.g., memory, sensory integration, control). Importantly, these inputs and outputs do not necessarily relate exclusively to the external world: they can include communication with other brain areas (e.g., internal models, planning activity).

To make this concrete, consider the 1-bit flip-flop (1BFF) computation (Fig. 1B, Computation): the output should reflect the sign of the most recent input pulse. Crucially, a computation specifies *what* must be done, but not *how*.

##### How are algorithms built by neural dynamics?

We define an algorithm as a set of rules that, when followed, enact a particular computation. In the CtD framework, these algorithmic rules are built from the neural dynamics of a circuit. Formally, neuronal circuits learn a *D*-dimensional latent dynamical system 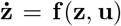 and corresponding output projection **x** = **h**(**z**) whose time-evolution approximates the input/output mapping **u** 1→ **x**.

For the 1BFF computation, we show that a relatively simple dynamical system **f** with output projection **h** (in this case, the identity function) can accomplish the desired mapping via an input-dependent flow field (Fig. 1B, Algorithm). Sufficiently large input pulses carry the state over a saddle point, which mediates the transition to a new attractor state.

##### How are these dynamics implemented in neural activity?

In the brain, neural dynamics arise from the underlying biophysics of neural circuits, which in turn arises from the physical properties of individual neurons, such as their ion channels, synapses, neuromodulators, and cell types. Yet the dimensionality of neural population activity is typically far lower than the total number of neurons, suggesting that neural dynamics can instead be modeled by a low-dimensional dynamical system embedded in a high-dimensional neural activity space (*N* ≫ *D*) [14, 18]. We denote this mapping from latent to neural activity **y** = **g**(**z**) : ℝ^*D*^ → ℝ^*N*^ and refer to **g** as the (true) observation model [38, 18, 68].

We simulated the observation model *g* of the 1-bit memory system using a simple linear-exponential transformation of the one-dimensional latent state into two-dimensional neural rates **n**, followed by sampling from an inhomogeneous Poisson process to generate neural spiking activity **y** (Fig. 1B, Implementation). Importantly, external inputs **u** are typically unobservable; they must be inferred from their effects on observable neural activity **y**.

#### 1.2.2 Climbing levels: From neurons to dynamics to computation

##### Data-driven modeling

While an interpretable account of a neural circuit spans all three levels, we typically have direct access only to the implementation level – the recorded neural activity **y**. We need methods that can accurately infer algorithmic features – dynamics **f**, observation model **g**, latent activity **z** from these neural observations (Fig. 1C, D, with supplied/inferred external inputs respectively). We use 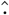 to denote model-inferred features (e.g., inferred dynamics 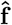 versus ground-truth **f**). Models that can be trusted to infer dynamics accurately (i.e., 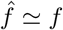) could allow us to climb from implementation (neural activity) to algorithm (neural dynamics), and then, hopefully, from algorithm to computation (Fig. 1E).

In recent years, a new class of “data-driven” (DD) models has emerged that are trained to reconstruct recorded neural activity as the product of inferred low-dimensional dynamics and observation models (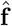 and **ĝ**, respectively) [16, 21, 19, 25, 41, 40, 53, 51, 55, 36, 42, 52, 29, 37, 68]. DD models are typically evaluated by how well they reconstruct simulated neural activity from synthetic systems with known ground-truth dynamics **f** . However, reconstruction accuracy can be a poor proxy for how faithfully a model captures the underlying dynamics, and commonly used synthetic systems lack many features fundamental to real neural circuits [56, 66, 40]. Moreover, the absence of a shared language of synthetic systems and performance criteria makes it difficult to compare published models or test new architectures systematically.

The primary goal of CtDToolkit is to provide a common, theoretically grounded foundation for model development and validation. It allows researchers both to benchmark models on standardized synthetic systems and to design customized datasets that mirror the structure, inputs, or feedback properties of their own experimental recordings, enabling targeted model development for specific neural systems. Together, these capabilities will accelerate progress toward DD models that can accurately and reliably infer neural dynamics from biological data.

##### Synthetic systems as proxies for neural circuits

Following the lead of the broader dynamics modeling community, most neural dynamics models have been validated using synthetic systems drawn from a class of well-characterized, low-dimensional chaotic attractors such as Lorenz or Arneodo [2, 16, 25, 21, 40, 41, 42, 50, 66]. At first glance, these chaotic attractors seem well-suited for dynamical model validation. For benchmarking generic dynamical systems, they offer several appealing properties. First, they are well-understood and identifiable, having no external inputs. Second, they exhibit chaotic dynamics – i.e., small changes in the system state can have a large impact on how the system develops over time – and therefore present a formidable modeling challenge. Finally, they are low-dimensional, which makes model training less computationally expensive and the results more easily interpretable.

Unfortunately, the features that make these systems appealing test-cases for generic dynamics models make them poor proxies for neural circuits that perform computation. First, chaotic attractors don’t “do” anything, lacking both the intended computation and external inputs that are fundamental features of goal-oriented neural circuits. Second, though some evidence exists that chaos may be beneficial in certain contexts – e.g., during learning, or for amplifying small signals in sensory integration tasks [13, 10, 65] – the degree of chaos in dynamical systems trained to perform tasks is typically lower than in these chaotic attractors [13, 8]. When chaos is present, its structure is likely constrained by the need for stable behavioral output: a system can be unpredictable along some latent dimensions while remaining predictable in its task-relevant readout. Finally, while the dimensionality of neural dynamics is still an open question, our rich behavioral repertoire suggests that neural dynamics are likely much higher-dimensional than these low-dimensional attractors. Therefore, we need our validation datasets to be *computational* (reflecting a goal-directed input-output transformation), *regular* (not overly chaotic), and *dimensionally-rich*.

Importantly, the same observed neural activity could in principle be generated by many different underlying assumptions—for example, about the form of inputs, the structure of feedback, or the dimensionality of latent dynamics. As a result, the choices made when simulating a dataset can dramatically affect the DD models that are best at capturing the ground-truth dynamics. DD model developers must take care to select or construct synthetic datasets whose properties best match those of the biological data to which they ultimately wish to apply their models. To enable this, our framework is designed to incorporate new datasets tailored to particular experimental needs, ensuring that validation exercises remain relevant and informative for the systems under study.

In Section 2.3, we describe how we obtain proxy systems with these properties by training dynamics models to perform specific tasks; we call these models “task-trained” (TT) models to distinguish them from the “data-driven” (DD) models that are trained to reconstruct neural activity.

##### Performance criteria

DD models are typically trained to reconstruct recorded neural activity, but recent work has shown that even near-perfect reconstruction does not imply that inferred dynamics are an accurate estimate of the underlying system (i.e., 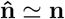 does not imply 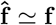) [34, 50, 63]. To address this problem, we have identified three key performance criteria that can collectively provide a more holistic assessment of DD model performance: *reconstruction, simplicity*, and *input accuracy*.

In this section, we provide intuition for the role of these criteria in model selection, while in Section 2.4, we provide formal definitions of specific metrics for quantifying model performance on each criterion.

##### Reconstruction

The first performance criterion is reconstruction, or the extent to which a model can re-generate neural activity from trials held-out from the training set. We note that this definition may differ from its usage in the broader field of dynamical systems identification, where it broadly refers to parity between the inferred and ground-truth systems. Poor reconstruction can be a symptom of an important mode of model failure called underfitting. Underfitting in this context refers to a model failing to capture the latent features **z** that underlie the observed neural activity patterns **y**, often because the model lacks computational capacity, is over-regularized, or has had insufficient time to train. In the last few years, reconstruction has improved dramatically as the field has moved from simple linear dynamics to more expressive model classes (e.g., stacked state-space models, transformers) [19, 59, 44, 49].

In Figure 1F, we show an example of underfitting: a hypothetical DD model whose inferred latents 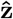 accurately capture the true latent activity **z** for the positive bit-flip (0 → 1), but fail to capture it for the negative bit-flip (1 → 0).

##### Simplicity

The second performance criterion is simplicity, with an associated failure mode of feature invention. Feature invention is the error-of-commission counterpart to underfitting’s error of omission: while underfit models miss real features, hallucinating models invent features that the true system lacks. Existing methods to quantify simplicity treat the latent activity **z** as the “feature” of interest, measuring how well the ground-truth system can predict the model-inferred latents 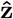 [50]. In principle, the same criterion could instead compare the inferred dynamics as a whole, such as entire flow fields rather than latent activity alone [34]. Models with both optimal simplicity and optimal reconstruction have a 1:1 mapping between the model-inferred latent features and the features of the underlying system.

One particularly insidious form of feature invention occurs when the invented features actually lead to improvements in reconstruction on both training and validation datasets. When ranking DD models by their ability to reconstruct neural activity, it has been shown that models with invented features often *outperform* those without [66]. This suggests that not only is reconstruction insufficient for DD model selection, but relying on reconstruction alone may be actively misleading [34]. While some metrics to quantify model simplicity have been released [40, 50], these methods require comparisons against the ground-truth system and therefore are not applicable to models trained on biological datasets. Of note, some attempts have been made to control invented features without reference to ground truth, e.g., by identifying the minimum-complexity regime in which independently trained models converge on consistent inferred features [34], or by constraining the model’s observation model **ĝ** to be injective [66].

We show an example of feature invention in Figure 1G. Though our hypothetical model perfectly captures the true latent activity **z**, it also infers an additional latent dimension 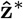. This “invented” activity evolves such that when the bit turns off, the state returns to a location that is different from where it began the trial. Alone, reconstruction and simplicity can each be insufficient to judge DD model quality, but together they can provide interpretable accounts of DD model performance when external inputs are known.

##### Input Accuracy

Now we consider the more realistic scenario in which the external inputs **u** are unknown and therefore must be inferred alongside the latent dynamics. The third performance criterion is input accuracy, or how well the model-inferred inputs **û** match the true external inputs **u** to the system.

In Figure 1H, we show a hypothetical model that accurately infers the latent activity 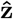 without inventing features, yet fails to capture the true dynamics **f**. Its inferred inputs **û** can reconstruct the data without relying on any intrinsic dynamics, while producing exactly the same predictions as the true model. Since both models are equally supported by the evidence, they cannot be disambiguated from observations alone—an instance of non-identifiability, the general problem in which distinct underlying systems produce indistinguishable observations. We call this particular failure mode dynamical misattribution. Because input accuracy cannot be evaluated on most biological datasets, this failure mode is particularly troubling: even when 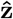 is accurate, 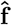 can still be a poor estimate of the true **f**.

## 2 Methods: Overview of the Computation-through-Dynamics Toolkit

### 2.1 Key Motivation

Our goal is to facilitate the development of data-driven (DD) models that can recover the unobserved latent activity **z**, dynamics **f**, observation model **g**, and external inputs **u** that underlie observable neural activity **y**. CtDToolkit supports this goal by addressing the two shortcomings described above—the limitations of existing synthetic systems and the insufficiency of existing performance metrics—within a single standardized, extensible codebase. Rather than crowning a single best model, it offers a structured process through which developers can iteratively test, refine, and tailor models to datasets with different properties, and share the resulting innovations with the community. Below, we walk through this process as three steps (Fig. 2): selecting a task, training DD models, and comparing them against task-trained ground truth, with an optional path for generating new synthetic datasets.

**Figure 2.**
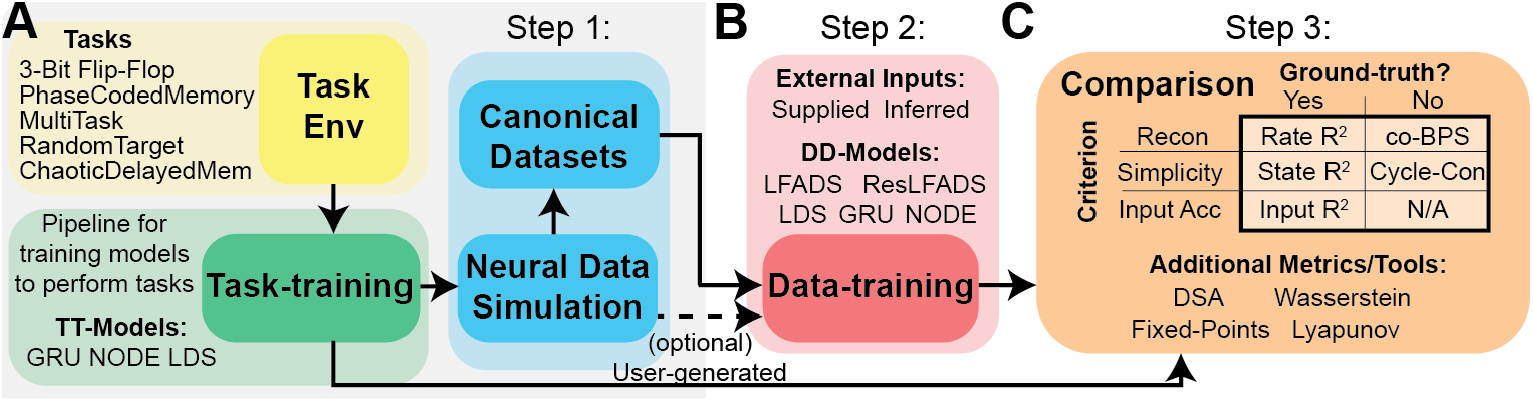
Features of the CtD Toolkit. **A**) Dataset generation pipeline-Task Environments (yellow) are taken as input to the task-training pipeline (green), which trains a dynamics model (available models: GRU, NODE, LDS) to perform the task. After training, the TT model is passed to a neural data simulator (cyan), which simulates spiking neural activity with configurable parameters. Users can either generate their own simulated datasets (dashed line) or use the pre-generated canonical datasets (solid line) **B)** Step 2: Data-training pipeline: Train a chosen DD model (available baseline models: LFADS, GRU, LDS, NODE) to reconstruct simulated neural activity, using either supplied or inferred inputs. Trained models are saved for analysis and comparison. **C**) Step 3: Model Comparisons-Top: Table of primary performance metrics, organized by criterion (rows), and need for ground-truth (columns). Bottom: Additional performance metrics/visualization tools.

### 2.2 Getting Started with CtDToolkit

#### Step 1: Select a task

First, users must select a task to test their data-driven models. At release, CtDToolkit provides pre-generated datasets for five tasks:

– *Three-Bit Flip-Flop*: Storing a 3-bit memory state that can be modified by external inputs.
– *MultiTask*: Performing a set of cognitive sub-tasks including category matching, sensory integration, etc.
– *RandomTarget*: Controlling the endpoint of a simplified biomechanical effector to instructed locations.
– *PhaseCodedMemory*: Remembering an input stimulus with reference to an input oscillation.
– *ChaoticDelayedMatching*: Comparing two sequentially-presented cues using near-chaotic dynamics.

Each canonical dataset includes everything needed for DD model training and evaluation, including simulated spiking activity and external inputs. We also provide the task-trained models from which the synthetic datasets were generated, allowing users to directly inspect the dynamics of the ground-truth system. Each dataset was chosen either to reflect a canonical property of biological neural circuits or to provide a simple, easily interpretable test case (Fig. 2A); we expand on these choices in Section 2.3.

#### Step 2: Train models on canonical datasets

Within the CtDToolkit data-training pipeline, users can fit models on canonical datasets and easily perform large hyperparameter sweeps (Fig. 2B). We support two training modes: supplied ground-truth external inputs or model-inferred external inputs. In addition to training user-defined models on synthetic data, users have access to a library of baseline models, including standard sequential auto-encoders and LFADS [25]. We define a common interface so that new models can be contributed to CtDToolkit, expanding the set of baseline models for future developers (Appendix D).

#### Step 3: Compare task-trained and data-driven models

CtDToolkit provides tools that facilitate the interpretation of task-trained and data-driven models, including metrics for quantifying performance on the criteria defined in Section 1.2.2. Some of the included metrics do not require ground truth and can be used to assess models trained on biological datasets (Fig. 2C). CtDToolkit also provides methods for data handling, model inference, fixed-point finding, and visualizations that allow users to quickly and easily assess model performance and iterate during model development (Section 2.4).

#### Optional: Generate new synthetic neural datasets

We do not intend for any single synthetic dataset to serve as a definitive benchmark for all models. Instead, we envision researchers iteratively generating and refining synthetic datasets whose properties best match the biological data they aim to study. To support this process, CtDToolkit provides templates for custom tasks and TT model architectures that integrate with the existing task-training and data simulation pipelines. The simulation pipeline itself is configurable, enabling users to explore how design choices such as neuron count, noise model, or the true observation model **g** affect DD model performance (Appendix C) and evaluation by Toolkit metrics (Section 2.4).

### 2.3 Task Environments

In contrast with previous validation datasets, CtDToolkit datasets were designed as biologically motivated synthetic benchmarks that capture selected challenges present in neural recordings while preserving full ground-truth access (see Section 1.2.2). These datasets were simulated from dynamical systems trained to perform specific tasks, defined in CtDToolkit as “task environments”, which provide simulated inputs and objective functions for quantifying TT model performance. Over the course of training, TT models learn dynamics that approximate the input/output mapping of the desired computation. For easy extensibility, task environment objects inherit from the Gymnasium Env class [60].

We include five task environments in CtDToolkit, chosen to provide a range in difficulty in 1) complexity of dynamics, 2) complexity of external inputs, and 3) present state of task interpretability. By *input complexity* we mean a qualitative descriptor of how amenable a task’s external inputs are to standard dynamical-systems analysis, rather than a formally defined scalar metric. We consider inputs to be *simpler* (i.e., lower complexity) when they are (i) piecewise constant over time and (ii) independent of the model’s outputs (open-loop). Piecewise constant inputs allow the dynamics to be analyzed under effectively stationary conditions within each epoch, making tools such as fixed point analysis directly applicable, whereas continuously varying or output-coupled inputs (e.g., in PhaseCodedMemory or RandomTarget) introduce time-dependent forcing that complicates interpretation of the intrinsic dynamics. We show the task schematics for 3BFF, MultiTask, and RandomTarget tasks in Fig. 4, for the PhaseCodedMemory task in Supp. Fig. S4, and for the ChaoticDelayedMatching task in Supp. Fig. S5.

#### Three-Bit Flip-Flop (3BFF)

3BFF is a 3-bit memory task with the goal of remembering the sign of the last input pulse on each of three noisy input channels (Fig. 3A). We chose to include 3BFF in CtDToolkit because it has 1) simple dynamics, 2) simple inputs (Fig. 3B), and 3) is well characterized by previous research [15]. TT models performing 3BFF typically exhibit low-dimensional activity organized into a cube, with stable fixed points at the vertices and unstable fixed points along the edges (Fig. 3C).

**Figure 3.**
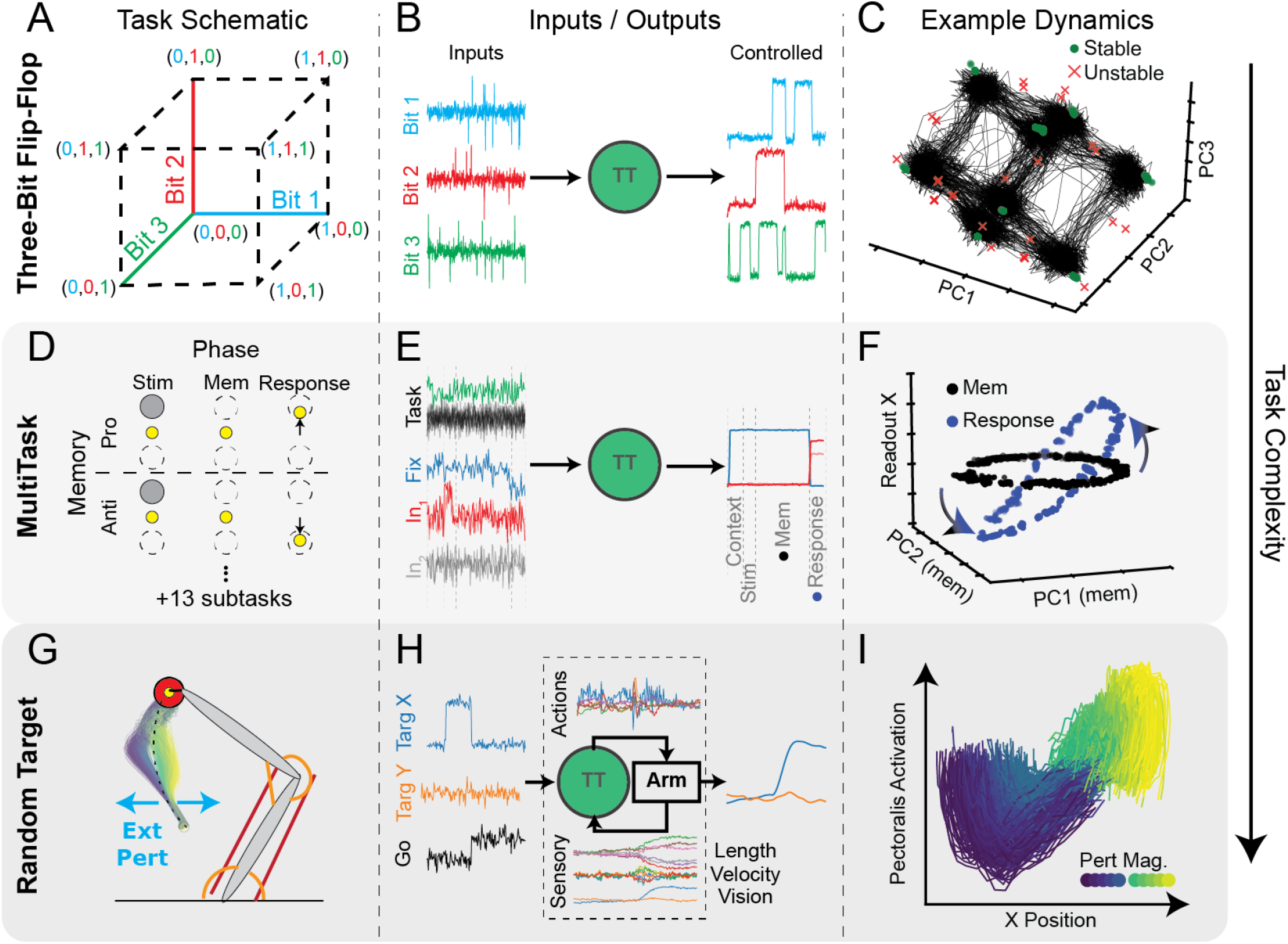
CtDToolkit Datasets have complex and interpretable dynamical features. **A**) Schematic of three-bit flip-flop (3BFF) task environment [15]. Three bits encoded according to inputs corresponding to 8 potential memory states. **B**) Inputs (left) and outputs (right) of the task-trained model for an example trial. **C**) Visualization of latent activity (black traces) and fixed points (FP) from canonical 3BFF TT network. Stable FPs (green circles) are found on vertices and unstable FPs (red × s) along edges of the cube. [23] **D**) Schematic of 2 of 15 tasks in MultiTask environment (MemoryPro, MemoryAnti). Each task has different rules for how to generate outputs. For full details, see [46] **E**) Example of single trial inputs/outputs from canonical MultiTask TT-GRU performing “MemoryPro” task. Task phases indicated by dashed vertical lines. **F**) Example FP architectures during two phases of MemoryPro task (Mem1 (black) and Resp (blue)). Ring of FPs rotates during the response phase into dimensions of the model that affect the output, executing the correct response based on location of activity in the ring attractor. **G**) Schematic of RandomTarget environment [61]. The TT model was trained to control the effector endpoint (yellow circle) to acquire target (red circle) after a go-cue was provided. External forces (cyan) were applied to perturb the hand, random in timing, direction, and magnitude. We show the resulting kinematics of the hand when applying left/right bumps of variable magnitude to the hand during a reach away from the body. **H**) Example of single trial inputs/outputs for canonical TT-GRU performing RandomTarget. As RandomTarget is coupled, actions (muscle commands) and sensory signals (muscle kinematics, visual inputs) are transmitted between the TT-GRU and the arm. **I**) Latent activity in a behaviorally-relevant plane of canonical TT-GRU when responding to perturbation in G. X-axis: Dimension of TT latent activity with strongest linear relationship to x-position of hand. Y-axis: magnitude of projection of TT latent activity onto Pectoralis motor-potent dimension. Bump magnitude (left:blue/right:green-yellow) shown in inset color-map.

Intuitively, RNNs trained to perform 3BFF use stable fixed points to remember the current memory state, and unstable fixed points to guide input-driven transitions between memory states. With low-dimensional activity and well-characterized dynamics, the 3BFF dataset is intended to provide data-driven model developers with clear and relatively unambiguous feedback about model performance. As such, we consider 3BFF to be the introductory task, and recommend it as the first task on which data-driven model performance is evaluated.

#### MultiTask

MultiTask is a previously published task [33, 46, 47] consisting of 15 distinct cognitive sub-tasks that has been used to investigate shared representations and learning in task-trained networks (Fig. 3D-F). Networks trained on MultiTask have 1) moderately complex dynamics, 2) simple, but relatively high-dimensional inputs, and 3) shared dynamical structures that provide a foothold for interpreting how the computations are performed.

Inputs to MultiTask include a one-of-15 (one-hot) input set that indicates which task is engaged on each trial, 4 noisy sensory input channels, and one fixation input (Fig. 3E). For each task, sensory inputs must be transformed into 3-dimensional outputs that follow a task-specific rule. Each task has piecewise inputs that change abruptly across phases– e.g., input presentation, delay period, etc. We show simplified versions of 2 example tasks and their phases in Fig. 3D (more details in Appendix B.1.2 and [33, 46]).

Models trained to perform MultiTask learn “dynamical motifs”: shared structures that are reused across multiple tasks [46]. Because MultiTask networks receive piecewise constant external inputs, fixed point (FP) analysis can be used to reveal how the dynamics change across task phases. Previous research studying MultiTask has provided a clear demonstration of how input-dependent changes in FP structure can shed light on how computation is organized in a dynamical system.

To demonstrate how the dynamics of the MultiTask network can illuminate the underlying computation, we analyzed the FP structure of a Multitask TT model performing the MemoryPro task (Fig. 3F). As shown previously [33], trained models exhibit a ring attractor of stable FPs that are used to memorize a continuous circular input. When the model was prompted to produce an output corresponding to the input it had seen previously [46], the FP ring rotated into a set of output dimensions that generated the correct response. With its complex and shared dynamical features and its piecewise constant inputs, MultiTask represents the current limits of our ability to interpret task-trained dynamics models, and therefore a challenging but tractable system for data-driven model validation.

#### RandomTarget

The RandomTarget task is based on planar arm reaching, a common experimental paradigm in motor neuroscience [4, 26]. Here, a model must learn to control a 2 degree-of-freedom arm—actuated by 6 Hill-type muscles and simulated with the MotorNet musculoskeletal modeling package [61]—moving the arm’s endpoint to target locations sampled randomly from within its range of motion, while also correcting for intermittent bump perturbations applied to the hand (Fig. 3G).

Models trained to perform this rich motor task have 1) intrinsic dynamics whose effects are difficult to disambiguate from those of their 2) high-dimensional, time-varying inputs, and 3) underlying computations that are not yet well-understood. The model receives both sensory (muscle lengths and velocities, hand endpoint position) and contextual (target position, go cue) inputs, and generates efferent commands that influence the force generated by each muscle. In contrast with previous tasks, this synthetic system is coupled to the environment, meaning that the inputs that it receives are affected by the motor commands it generates (and vice versa, see Fig. 3H). As the inputs are constantly changing, this can make direct interpretation of the dynamics very difficult.

To find a signature of dynamical computations underlying movement corrections, we trained a model on RandomTarget and tested how it learned to correct for left/right perturbations in the middle of a reach away from the body (Fig. 3G, colored trajectories). To visualize how sensory information is transformed into corrective muscle commands, we projected the TT model’s activity onto the plane defined by the pectoralis muscle activation (responsible for shoulder flexion) and the endpoint *x*-position. We found a stereotyped rotational pattern in this plane that varied smoothly with the perturbation magnitude and direction (Fig. 3I, color gradient). TT model dynamics have apparently learned to perform goal-directed sensory-motor transformations that correct for perturbations to the desired hand trajectory; this structure resembles a feedback control system [39].

Unlike the other canonical datasets, the main challenge in RandomTarget is not modeling its dynamics, which are relatively simple, but rather distinguishing the model’s internal dynamics from the coupled dynamics of the TT model and the arm. As biological circuits also receive complex, time-varying inputs that likely reflect coupled dynamics, RandomTarget is also arguably most similar to datasets that might be collected experimentally.

#### PhaseCodedMemory

Cyclic, oscillatory dynamics are a common feature of neural recordings, and we wanted to include a task that lets DD models be tested on a dataset with this dynamical structure. Based on recent work investigating the relationship between oscillatory activity and memory [62], the PhaseCodedMemory (PCM) task simulates a computational mechanism for short-term memory called “phase-coding” that may be present in the hippocampus. Phase-coding models remember the content of input signals by encoding them in the relative phase of an internal oscillation to a reference input oscillation (in this case, an oscillatory signal similar to LFP), as opposed to by differences in mean firing rate across stimulus conditions (i.e., a “rate-coding” strategy). In addition to producing strongly cyclic activity, PCM represents computation in a brain area, the hippocampus (HC), from which recordings differ qualitatively from typical intracortical MEA recordings in both their yield and typical firing rates. The PCM task exhibits simple, interpretable dynamics, yet its time-dependent inputs introduce additional challenges absent from 3BFF. We discuss the differences between the PhaseCodedMemory dataset and the other canonical datasets in further detail in Appendix B.1.4.

#### ChaoticDelayedMatching

While the canonical datasets above were designed to emphasize the *regular* dynamics typical of task-trained networks (Section 1.2.2), chaotic dynamics constitute a large and well-studied class of systems that are potentially relevant to neural computation, particularly in the contexts of learning and sensory processing [13, 8]. To extend the relevance of CtDToolkit to this regime and to better connect with the broader dynamical-systems reconstruction literature, we include ChaoticDelayedMatching, a delayed non-match-to-sample task based on work by Miconi [20]. Here, a recurrent network of tanh units initialized in a near-chaotic regime (via gain-scaled random recurrent weights) is trained to compare two sequentially-presented binary cues (A or B), separated by delay periods, and to report during a response window whether the cues match (A-A, B-B) or differ (A-B, B-A). This defines a simple and well-specified input-output mapping while requiring the network to maintain and transform information over time. The resulting dynamics exhibit key signatures of chaos in the weakly-driven regime, including sensitivity to initial conditions assessed via the perturbation-driven divergence of nearby trajectories (Supp. Fig. S5D,E), yet still produce the correct task-relevant output during the response period—reflecting stable computation embedded within high-dimensional, near-chaotic dynamics. We provide additional details on the task and task-trained model in Appendix B.1.5.

### 2.4 Metrics and Visualizations

Recent research has shown that reconstruction can be an unreliable indicator of model quality [34, 50]. CtDToolkit includes new metrics that provide a more holistic account of model performance on key criteria of reconstruction, simplicity, and input accuracy (Section 1.2.2). Some of these metrics (Rate *R*^2^, State *R*^2^, Input *R*^2^) require access to ground-truth, while others (co-smoothing bits-per-spike and cycle-consistency) can be applied even when the ground-truth is unknown. We note that all synthetic datasets in this work were generated with a linear–exponential observation model **g**, in which latent activity is linearly projected to log-firing-rates before exponentiation; unless otherwise noted, the metrics described below assume this linear mapping between latent states and log-rates.

Also included in CtDToolkit are several previously released methods: Dynamical Similarity Analysis (DSA [56]), which quantifies aspects of both reconstruction and simplicity; and two common metrics from the broader field of dynamical systems identification—a distribution-matching metric based on the Wasserstein distance, and an estimate of the full Lyapunov spectrum that characterizes system stability and sensitivity to initial conditions. We show in Section 3 and Section E.7 how these metrics can be used to identify common failure modes of DD models and guide model and hyperparameter selection. We used validation datasets for all metrics and visualizations to prevent overfitting from affecting our interpretations of DD model quality.

#### 2.4.1 Metrics comparing DD models to ground-truth

##### Rate *R*^**2**^

Rate *R*^2^ measures a DD model’s ability to capture features of the true system (i.e., firing rates). Rate *R*^2^ is obtained by finding the coefficient of determination between the true and predicted firing rates of each neuron in the dataset, yielding *N R*^2^ values, which we summarize with the variance-weighted mean across all neurons. By weighting by each neuron’s firing rate variance, Rate *R*^2^ is not artificially deflated by poorly predicted neurons that are only weakly modulated by the task. A Rate *R*^2^ value of 1 indicates perfect reconstruction of the underlying rates for all neurons. As Rate *R*^2^ requires access to ground-truth neural firing rates, it is not typically applicable for DD models trained on biological datasets.

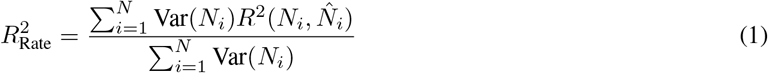

##### State *R*^**2**^

The possibility of invented features means that a DD model’s reconstruction is an unreliable indicator of accurately inferred dynamics (see Fig. 1G). The first CtDToolkit metric for quantifying model simplicity is called State *R*^2^ [50]. State *R*^2^ assesses the degree to which the inferred latent activity contains features that cannot be linearly explained by the ground-truth latent activity. State *R*^2^ is computed by fitting an affine transformation from the true hidden unit activity of the TT model (*z* ∈ ℝ^*D*^) to the activity inferred by the DD model 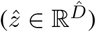, and then computing the variance-weighted coefficient of determination. As State *R*^2^ requires access to ground-truth latent activity, it is not applicable for DD models trained on biological datasets.

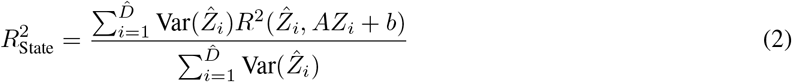

##### Input *R*^**2**^

As shown in Figure 1H, accurately inferred inputs are critical to ensure that a DD model’s inferred dynamics are trustworthy. We provide a metric called Input *R*^2^ that assesses the accuracy of these inferred inputs. Similar to State *R*^2^, Input *R*^2^ is calculated by fitting an affine transformation from inferred inputs 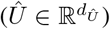 to true inputs 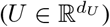, then computing the variance-weighted coefficient of determination across input dimensions. As Input *R*^2^ requires access to ground-truth external inputs, it is not applicable for DD models trained on biological datasets.

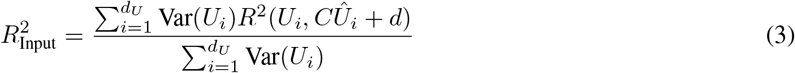

#### 2.4.2 Metrics assessing DD models without ground-truth

Because dynamics and external inputs are often unobservable for biological systems, it is important that we develop metrics that can be applied even when we are unable to compare against ground-truth. In this section, we describe two metrics, co-BPS and cycle-consistency, that quantify DD model performance on the criteria of reconstruction and simplicity.

##### Co-Smoothing Bits-per-Spike

Co-smoothing bits-per-spike (co-BPS) is a previously released method to assess DD model reconstruction [49]. co-BPS quantifies how well spiking activity in a set of held-out neurons can be predicted by models that only have access to a set of held-in neurons, under an assumed Poisson observation model; it is positive when predicted held-out activity is more informative than each neuron’s mean firing rate, and does not require ground-truth firing rates. Co-smoothing is specific to autoencoder-based DD models that condition on observed spikes at inference time, where the held-in/held-out split prevents trivial identity mappings. For DD models that do not require spikes as input, a likelihood computed across all neurons on held-out trials uses more of the data and avoids sensitivity to the neuron split, and should be preferred. co-BPS is calculated as:

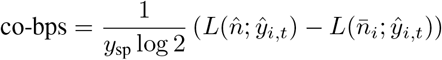

where *y*_sp_ is the total number of spikes from the held-out neurons, *L* is the negative log-likelihood function, 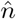 represents the model-predicted firing rates, 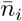 is the mean firing rate of neuron *i*, and *ŷ*_*i,t*_ are the observed spike counts for neuron *i* at time *t*.

##### Cycle-Consistency

Cycle-consistency estimates DD model simplicity *without* requiring ground-truth latent activity, by interrogating the relationship between a model’s inferred latent activity and its inferred firing rates.

The key idea is that a model can only invent features without paying a reconstruction penalty if its inferred observation model **ĝ** is *functionally non-injective* on the data—that is, if some changes in latent activity produce no (or negligible) change in predicted rates. The canonical example is a linear observation model with a null space, in which latent activity can vary within the null space without affecting predicted neural activity (Fig. 1G). The relevant property is functional injectivity of the *learned* mapping at the end of training, not whether the architecture imposes it as a hard constraint: constraining the observation model to be injective [66] is one way to encourage it, but unconstrained networks may also learn functionally injective mappings.

Cycle-consistency quantifies the functional injectivity of **ĝ** on held-out data and uses it as an indirect measure of feature invention: models with high cycle-consistency have few inferred latent features that are not reflected in their predicted neural activity. We emphasize that functional injectivity is necessary but not sufficient to rule out invented features—a model may introduce latent structure that changes predicted rates only negligibly, escaping detection by spike-level metrics such as co-BPS—so cycle-consistency is best treated as one diagnostic among several, complementary to ground-truth measures (State *R*^2^, Rate *R*^2^) where available.

Mathematically, cycle-consistency tests whether a linear model can re-generate the inferred latents 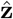 from the inferred neural activity 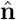. We first apply principal components analysis (PCA) to the inferred latents and inferred log-rates to sort dimensions by explained variance, then fit a linear observation model from the inferred latents to the inferred log-rates, apply singular-value thresholding to remove effectively-null dimensions (configurable; by default, those associated with <1% of the total variance) [7], and use its pseudo-inverse to re-generate the latents 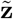. We then report the variance-weighted *R*^2^ between inferred latents 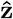 and re-generated latents 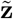:

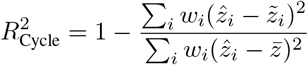

Here *w*_*i*_ are weights proportional to the variance of each latent dimension; because the metric uses only inferred variables, it can be computed without access to the ground-truth system.

This linear version assumes the canonical linear–exponential link from latent activity to rates; under other link functions or nonlinear manifolds it can mislead, so CtDToolkit also includes a previously published nonlinear version (`nl_cycle_con`) for arbitrary observation models [66] (Supp. Fig. S9, Section E.4).

##### Distribution Matching (Wasserstein Distance)

To connect CtDToolkit with the broader dynamical-systems identification literature, we include a distributional metric based on the empirical Wasserstein distance. Rather than comparing latent trajectories directly, it compares the distribution of *observed* validation activity to the distribution of model-predicted activity, after projecting both into a common, observation-derived low-dimensional space (optionally with a delay embedding to capture local temporal structure). Following Patel and Ott [57], we summarize the discrepancy as an average of per-dimension one-dimensional Wasserstein-1 distances, each normalized by the range of the reference data [54]; lower values indicate closer agreement. Full implementation details—temporal binning and smoothing, PCA projection, delay embedding, and normalization— are given in Appendix E.5.

Because this metric operates in the rate/observation space rather than the latent space, it is likely insensitive to invented features that fall in the null space of the inferred observation model **ĝ**: such features leave predicted activity unchanged, and so do not register in a distributional comparison of observed and predicted activity.

#### 2.4.3 Additional Metrics and Visualization

##### Dynamical Similarity Analysis (DSA)

DSA [56] is a nonlinear similarity metric that compares the spatiotemporal structure of two dynamical systems. Applied to CtDToolkit, DSA measures whether DD models reconstruct the core dynamic features of their TT model. More specifically, DSA captures whether these features (e.g. fixed points) are aligned in a one-to-one fashion and the DD model has neither invented superfluous features nor deleted relevant ones. DSA has three components (further details are in Appendix E.3): first, the method lifts the data into a higher dimensional space using kernels or delay embeddings [17, 28]. Second, a linear system is fit to each embedding, resulting in dynamics matrices *A*_*x*_, *A*_*y*_. Finally, these systems are compared using an extension of Procrustes Analysis [43]. In this paper, we used the angular form of the distance, which scales from 0 (most similar) to *π/*2 (most dissimilar). In practice, however the full range is not always used, and so relative distances are typically emphasized over absolute. As applied here, DSA compares the inferred dynamics of a DD model against the dynamics of the ground-truth TT model, and therefore requires access to the true latent activity; it is consequently applicable only to synthetic datasets where ground truth is available, and not to biological recordings.

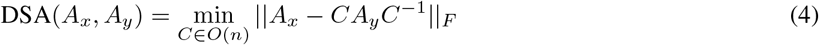

##### Fixed-Point Visualization

In addition to the quantitative metrics, CtDToolkit provides tools for qualitatively assessing the true and inferred dynamics through their fixed point structures. We distinguish *true fixed points*, where the flow is exactly zero (*f* (*z, u*) = *z*), from *functional fixed points*: regions of state space where the dynamics are sufficiently slow (or contracting) to appear stable over behaviorally relevant timescales and to permit linear approximation. In practice, the fixed-point finder below locates functional fixed points by minimizing (rather than exactly zeroing) the speed of the dynamics. The arrangement and stability of these fixed points provide key insights into the local behavior of the system. To find fixed points, we find locations in the model’s state space 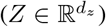 that minimize the magnitude of the kinetic energy of the system (Δ*h*^2^ ~ *q*). CtDToolkit includes an extension of a previously-released fixed point finding toolkit [23] for use with both TT and DD models. Since input changes can alter fixed point structure [46, 31], our modified fixed point finder accepts user-defined static inputs, allowing researchers to visualize how fixed point structures change as inputs change.

##### Lyapunov Spectrum

The Lyapunov spectrum characterizes the long-term behavior of a dynamical system by quantifying how quickly nearby trajectories diverge or converge along different directions; it is a standard tool for comparing the stability and sensitivity-to-initial-conditions of an inferred model against a ground-truth system. Following [67], we estimate the full spectrum {*λ*_1_ ≥ · · · ≥ *λ*_*D*_*}* of the discrete-time system **z**_*t*+1_ = *f* (**z**_*t*_, **u**_*t*_) using the QR-decomposition method applied to Jacobians sampled along trajectories. The maximal exponent *λ*_max_ is the component most readily estimated from data and most directly compared across TT and DD models (*λ*_max_ *>* 0 indicates chaos; *λ*_max_ *<* 0, asymptotic stability), but the full spectrum is needed to distinguish, for example, a chaotic *attracting* set from a divergent one—via summaries such as the sum of exponents ∑_*i*_ *λ*_*i*_ (the mean rate of phase-space volume contraction) and the Kaplan–Yorke dimension. CtDToolkit provides `compute_lyapunov_spectrum` (and `Analysis_TT.compute_lyapunov_spectrum`) routines for this purpose. We validated the estimator on the canonical Lorenz and Rössler systems, recovering spectra that closely match their literature references (Supp. Fig. S10). Full algorithmic details and validation values are provided in Appendix E.6.

## 3 Results

### 3.1 Canonical datasets provide a library of biologically-motivated dynamical systems

#### Task-training and dataset simulation pipeline

We first used a CtDToolkit task environment to generate training data—external inputs, target outputs, and the additional information needed for training (Appendix B.1)—and trained a task-trained (TT) dynamics model to produce outputs that minimize the task’s objective function (Fig. 4A, left). By the end of training, each TT model had learned latent dynamics that performed the computation defined by its task environment.

We then simulated neural datasets from the hidden-unit activity of the TT models (Fig. 4A, middle; Appendix B). We fed the task-training trials to each TT model, recorded its hidden activity, and—for most datasets—sampled *N* hidden dimensions without replacement, scaled them, and used them as the rate parameter of a Poisson process. (The PhaseCodedMemory dataset is an exception: to emulate a larger, low-firing-rate hippocampal population, its neuron count exceeds the TT latent dimensionality, so some latent dimensions are sampled more than once with independent observation noise; Appendix C.1.) For every dataset, this yielded spiking activity from *N* simulated neurons, which we then split into held-in and held-out neuron sets. These held-out neurons should not be confused with held-out validation trials; all performance metrics were computed on validation trials to avoid spurious results from overfitting.

We designed the 3BFF, MultiTask, RandomTarget, and ChaoticDelayedMatching datasets to have properties that resembled those of common intracortical multi-electrode array recordings (e.g., from Utah arrays), with *N <* 100 and moderately high firing rates. In contrast, the PhaseCodedMemory dataset was designed to emulate the properties of datasets recorded from CA1 neurons in the hippocampus, and was therefore composed of 600 neurons (500 held-in, 100 held-out) with relatively sparse activity (~ 2 Hz) (Supp. Fig. S4A-E). We trained DD models to reconstruct held-in and held-out neural activity using only held-in neural activity (Fig. 4A, right).

#### Key observations from canonical datasets

To provide some intuition for how common DD models perform on the canonical datasets, we visualized the ground-truth and inferred latent activity for the tasks included in CtDToolkit and compiled the full performance-metric sweeps across model sizes for each: 3-Bit Flip-Flop (3BFF; Supp. Fig. S11), MultiTask (Supp. Fig. S12), RandomTarget (Supp. Fig. S13), PhaseCodedMemory (Supp. Fig. S14), and ChaoticDelayedMatching (Supp. Fig. S15). We provide descriptions of the three baseline DD models in Section D.3.

##### 3-Bit Flip-Flop (3BFF)

For 3BFF (Figure 4B), we found that LFADS and GRU models inferred latent activity that closely resembled the top three principal components (PCs) of the ground-truth latent dynamics. The LDS model, however, lacked the computational capacity to capture even the relatively simple dynamics of 3BFF. In other words, LDS suffers from the failure mode of *underfitting* described in Section 1.2.2. We note that the LDS failure is more precisely described as *structural underfitting*: the class of representable (linear) dynamics simply does not contain the target system, regardless of parameter values. This contrasts with the capacity-limited underfitting of more expressive models (e.g., a NODE with too few latent dimensions), which can in principle represent the true dynamics but fail to do so with insufficient capacity.

##### MultiTask

In the MultiTask dataset (Figure 4C), we visualized the behavior of our DD models by projecting the inferred latent activity from MemoryPro trials onto three output dimensions (Fixation, X output, Y output) for each model. We found that both GRU and LFADS models qualitatively captured the structure of the ground-truth latent activity in this 3D subspace. However, the LDS model had significant errors in its projection onto output dimensions during the memory period. This suggests that LDS was unable to gate its response appropriately, resulting in premature output generation. Similar to 3BFF, these results indicate that LDS lacks computational capacity, and as a result also underfits the dynamics of the MultiTask dataset.

##### RandomTarget

Given its complex time-varying inputs, performance on RandomTarget (Figure 4D) is the most difficult to interpret. Examining the top three PCs of latent activity (color-coded by reach direction), all three models—including the limited-capacity LDS—appeared to capture the high-level structure of the ground-truth latents. This apparent success is deceptive: because every model was supplied the rich, time-varying ground-truth inputs (muscle lengths and velocities, vision, and goal/timing information) during both training and inference, much of that structure can be inherited directly from the inputs rather than generated by a model’s own intrinsic dynamics. That even LDS—which underfit the far simpler 3BFF and MultiTask—looks good here is precisely the warning: on RandomTarget, matching the ground-truth latents says little about whether a model has captured the underlying dynamics. The real challenge is to disentangle intrinsic dynamics from input-driven activity. This tight coupling between the TT network and another dynamical system makes RandomTarget a more realistic and conceptually challenging addition to CtDToolkit than alternative datasets [39].

##### PhaseCodedMemory

We show the performance of one model class (LFADS) on the PhaseCodedMemory dataset in Supp. Fig. S16. We found that the LFADS model inferred latent activity that pointed towards a “rate-coding” strategy of memory, in contrast with the “phase-coding” strategy of the dynamics of the ground-truth TT model. This contrast is also visible geometrically: in the TT model the latent-space and rate-space principal components share the same structure, whereas in the DD models they diverge—the inferred latents separate the two stimulus conditions even though the reconstructed rate-space structure does not. Notably, this divergence emerges even as reconstruction *improves* with model capacity: across the latent-size sweep, Rate *R*^2^ increases (from 0.68 to 0.77) while State *R*^2^ degrades (from 0.97 to 0.92), so a practitioner selecting models on reconstruction alone would be most confident in precisely the regime where the inferred latent geometry is most misleading (Appendix E.8). This result illustrates how interpreting a DD model’s inferred latent dynamics at face value—without cross-checking against a ground truth or the simplicity metrics advocated here—could lead researchers to incorrect conclusions about how computation is performed in the brain, even when the observed activity is reconstructed well.

##### ChaoticDelayedMatching

ChaoticDelayedMatching extends the canonical set to the near-chaotic regime, in which task-relevant computation is embedded in high-dimensional, sensitive-to-initial-conditions dynamics rather than the regular, attractor-driven dynamics of the other canonical datasets. ChaoticDelayedMatching thus provides a setting in which the reconstruction and dynamical-stability metrics in CtDToolkit (Section E.7) can be exercised on a system whose autonomous dynamics are qualitatively closer to those of the cortical networks the toolkit ultimately aims to model.

### 3.2 Reconstruction and simplicity metrics guide model selection by identifying common failure modes

Next, we sought to demonstrate that CtDToolkit metrics can provide actionable feedback for improving DD models. We trained DD Neural ODE (NODE) models [22] to reconstruct synthetic spiking activity from our canonical 3BFF TT model (Appendix D.3). Because a NODE’s latent dimensionality controls the expressivity of its dynamics, we trained 5 randomly initialized models at each hidden size 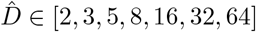 (35 models total), all supplied with ground-truth external inputs; this lets us study how model capacity drives distinct failure modes.

We emphasize that latent dimensionality is only one of many hyperparameters that affect model performance: regularization strength, input penalties, training duration, architecture, and optimization settings can all influence whether an expressive model underfits, faithfully recovers the dynamics, or invents spurious features. For example, the number of training epochs can itself act as a control on model complexity: prior work has shown it can dramatically affect the fidelity of the inferred dynamical landscape even while generalization performance remains high [34].

#### Reconstruction

We measured each model’s reconstruction accuracy with Rate *R*^2^ and co-BPS (as described in Section 2.4; Fig. 5A). Inspecting the inferred firing rate of a single held-out neuron, we found that while DD NODE with 2 dimensions (NODE-2) was able to capture some aspects of the true firing rate, its reconstruction was substantially worse than a DD NODE with 8 hidden units (NODE-8, Fig. 5B, dark blue, green, respectively). Models with a very small number of hidden units apparently had insufficient capacity, and therefore suffered from *underfitting*.

**Figure 4.**
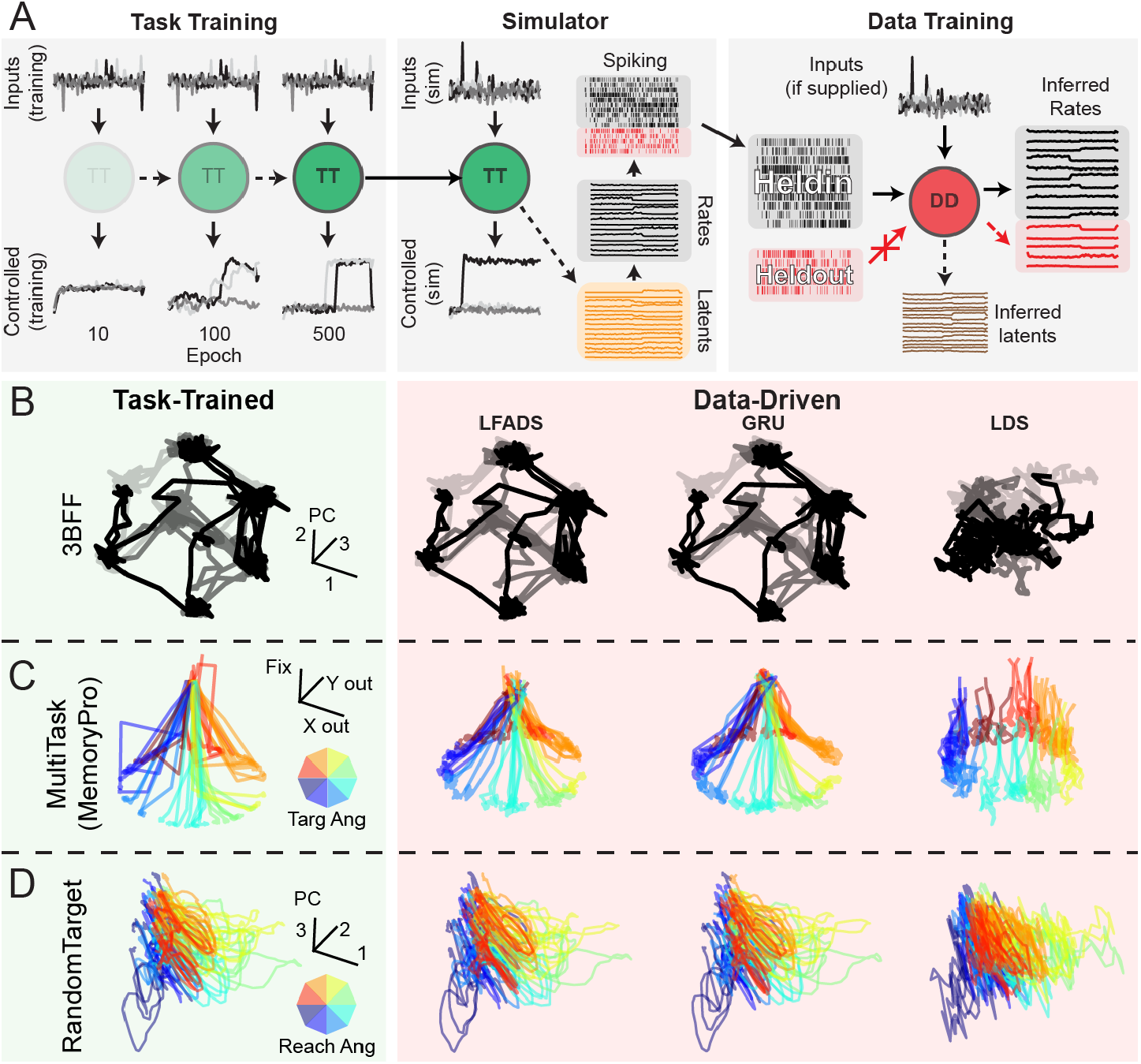
Task-trained and data-driven modeling pipeline, and example inferred latent activity for a subset of canonical datasets: **A)** Schematic of task-training, simulation, and data-training pipelines: (left) Task-training: model inputs converted to control signal by TT model dynamics. Over training epochs, TT models learn to generate control output that accomplishes the task. (middle) Fully-trained model provided as input to the simulator. Simulator was used to sample latent activity from the TT model, convert to rates, and simulate spiking using a Poisson noise process. (right) Simulated spiking split into held-in and held-out sets of neurons, and into training and validation trials. Held-in spiking is fed as input to the DD model, which is trained to infer rates for both held-in and held-out neurons. Hidden activity of the DD model is referred to as inferred latent activity. All metrics are computed on the validation trial set to prevent our results from being due to trivial overfitting. **B-D)** Example ground-truth latent activity (left column, green, without noise in input channels for improved visualization) and latent activity inferred by DD models (from left to right, LFADS, GRU, LDS) for three canonical datasets. DD model inferred latent activity was aligned with TT latent activity using an optimal affine transformation. **B)** Latent activity (top 3 PCs) for three representative trials (indicated by opacity) for 3BFF canonical dataset. **C)** Latent activity for representative trials of MemoryPro subtask for MultiTask canonical dataset. Color of each trace indicates the angle of the correct output response (inset color wheel). Latent activity was projected onto a 3D subspace defined by response 1 (x-axis), response 2 (y-axis), and fixation (z-axis) output dimensions. For DD models, we transformed the inferred latents to the space of the ground-truth latents via an optimal affine transformation, then projected through the ground-truth output mapping. **D)** Example inferred latent activity (top 3 PCs) for representative trials for RandomTarget canonical dataset. Color indicates the reach angle for each trial.

**Figure 5.**
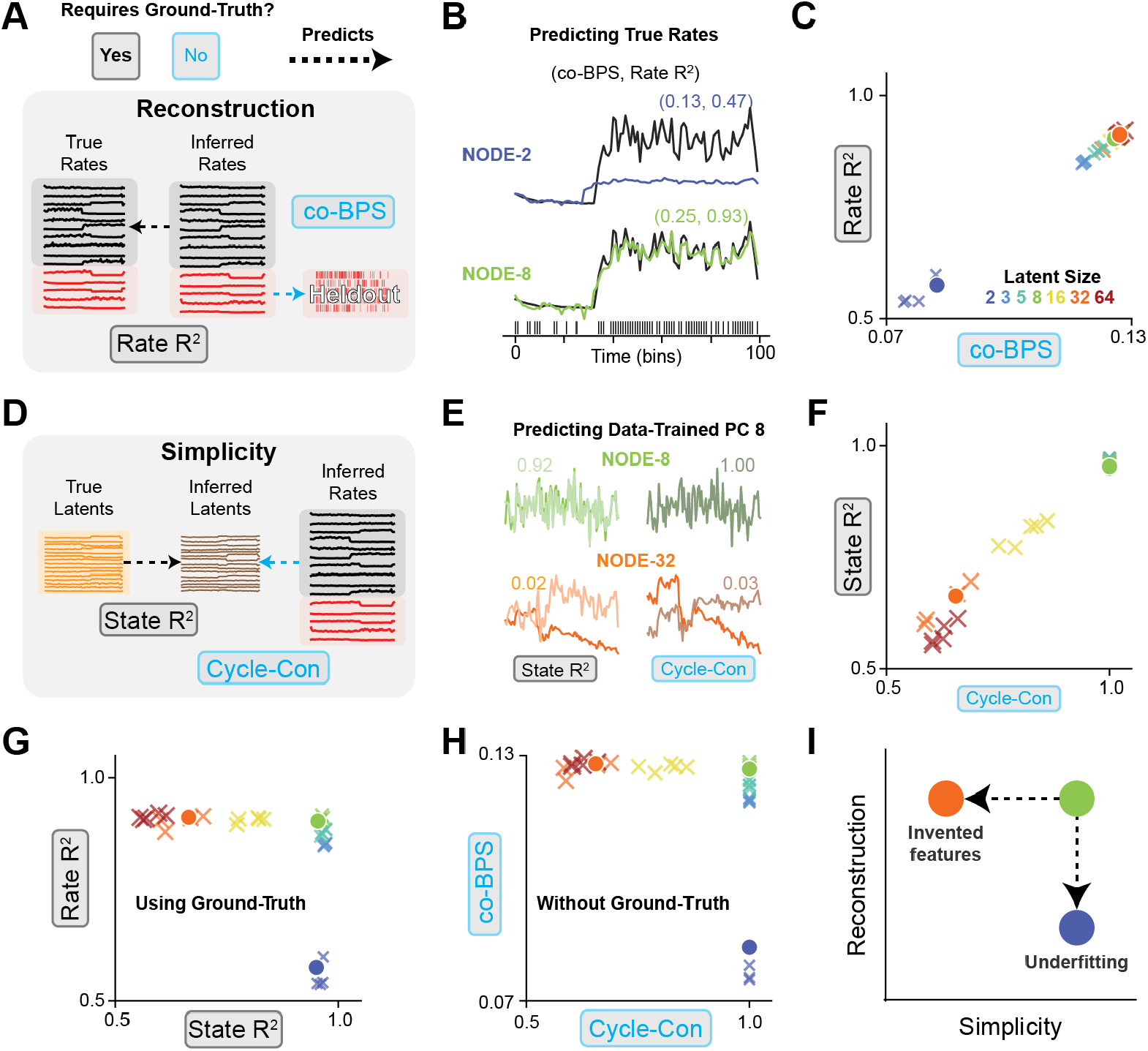
Reconstruction and simplicity metrics for data-driven modeling of 3BFF dataset. **A)** Schematic of reconstruction metrics: Rate *R*^2^ measures prediction accuracy of inferred rates vs. true rates. co-BPS measures accuracy of held-out spiking predictions. **B**) Single-trial predicted firing rate of one held-out neuron from models NODE-2 (2 hidden unit NODE, dark blue) and NODE-8 (8 hidden unit NODE, green). Performance indicated by (Rate *R*^2^, co-BPS) inset. **C)** Scatter plot of Rate *R*^2^ vs. co-BPS for NODE models ranging from 2 to 64 hidden units. Inset legend color indicates hidden size. Circles indicate example models in panel B, Xs indicate other randomly initialized models with the indicated latent dimensionality. **D)** Schematic of simplicity metrics: State *R*^2^ measures prediction accuracy of inferred latents from true latents, cycle-con measures accuracy of inferred latents from inferred rates. **E)** Inferred latent activity PC8 (green = NODE-8, orange = NODE-32), and linear prediction from true latents (left, State *R*^2^) or inferred rates (right, cycle-con). **F)** Scatter plot of State *R*^2^ vs. cycle-con for same models in panel C, indicated by colored markers. **G)** Scatter plot of Rate *R*^2^ vs. State *R*^2^, measuring reconstruction and simplicity with access to ground-truth. **H)** Scatter plot of co-BPS vs. cycle-con, showing metrics of reconstruction and simplicity that can be computed without access to ground truth. **I)** Schematic of underfitting (dark blue, NODE-2) and invented features (orange, NODE-32) failure modes, for comparison with panels G, H.

To confirm that our metric for quantifying reconstruction without access to ground-truth firing rates (co-BPS) gave similar results, we plotted the performance of our 35 models on these two metrics and found a strong linear relationship between Rate *R*^2^ and co-BPS (Fig. 5C). The co-smoothing metric used to quantify reconstruction in previous benchmarks [49] seems to accurately capture reconstruction, even in the absence of ground-truth.

#### Simplicity

We assessed the simplicity of each model to determine the extent to which it had invented features (Fig. 5D). To do this, we visualized the inferred latent activity in a low-variance dimension (8th principal component) for two DD models (Fig. 5E, green: NODE-8, orange: NODE-32), along with linear predictions of this dimension from the true latents (State *R*^2^, left column) and inferred rates (Cycle-Con, right column). We found that the 8th PC of NODE-32 was predicted poorly compared to the 8th PC of NODE-8. The presence of features in the inferred latent activity that cannot be predicted from the ground-truth suggests that the more expressive models tended to invent features that didn’t exist in the underlying system.

To confirm that our metric for quantifying simplicity without access to ground-truth latents (cycle-consistency) gave similar results to our ground-truth simplicity metric (State *R*^2^), we plotted the performance of our 35 models on these metrics and, like our reconstruction metrics, found a strong linear relationship (Fig. 5F). Additionally, we found that as NODE hidden size increased, there was a consistent decline in the simplicity for both metrics, evidence that more expressive models can suffer from the failure mode of *feature invention*.

#### Model selection across multiple criteria

We plotted the two criteria together, showing metrics that rely on access to ground-truth (Fig. 5G) and metrics that do not (Fig. 5H). Both sets of metrics qualitatively recapitulate the failure modes described by our performance criteria presented in Figure 1. Models with insufficient expressivity suffered from underfitting, while models that were too expressive invented features (Fig. 5I). We found similar results when we evaluated the performance of DD models trained on the NBFF, MultiTask, RandomTarget, PhaseCodedMemory, and ChaoticDelayedMatching canonical datasets (see Supp. Figs. S11, S12, S13, S14, S15).

#### Complementary metrics

We also computed two metrics from the broader dynamical-systems identification literature—the Wasserstein distance and the Lyapunov spectrum—for all DD models (Supp. Fig. S11C). The Wasserstein distance gave a ground-truth-free distributional comparison between model-predicted and observed activity across latent sizes; in practice it generally tracked reconstruction performance and was insensitive to invented features, consistent with its operating in the rate/observation space. The Lyapunov spectrum, in contrast, characterized the stability of each model’s inferred dynamics directly from its flow field. Estimating the spectrum for both TT and DD models, we found that the DD models whose maximal Lyapunov exponent best matched the ground-truth TT estimate were those with 8–16 latent dimensions: precisely the regime in which reconstruction had saturated and simplicity was beginning to degrade. The Lyapunov spectrum thus offers a complementary axis for model selection: rather than comparing inferred activity patterns (as the reconstruction and simplicity metrics do), it compares the inferred *dynamics* themselves, and here it independently pointed toward the same well-balanced models. Together, these metrics give users additional tools for diagnosing DD model failures.

Nonlinearities in the mapping between two topologically identical systems could lead some metrics to incorrectly diagnose a model with invented features. We caution that geometric differences cannot always be safely ignored: recent work shows that the geometry of inferred activity can itself reflect genuinely distinct task solutions rather than trivial reparameterizations [69]. For this reason, CtDToolkit includes both linear and nonlinear metrics as complementary measures of model similarity, leaving the choice of whether to treat geometric differences as meaningful to the user.

### 3.3 Input *R*^2^ helps diagnose models that fail by dynamical misattribution

The experiments above assumed known ground-truth inputs, which biological neural datasets rarely provide. A further CtDToolkit metric, Input *R*^2^, quantifies how accurately a model infers these unseen external inputs. Throughout this work we assume the dimensionality of the external inputs is known; this lets us evaluate inferred dynamics without the additional ambiguity of input-dimensionality estimation, which we regard as an important related problem left for future work.

A common architecture for input inference (originally introduced in LFADS [25]) is shown in Figure 6A. Neural activity is provided to two separate modules, the controller and the generator, which produce the inferred inputs and underlying dynamics respectively. Input inference is in general an ill-posed problem because observations alone are insufficient to distinguish a purely input-driven system from a purely autonomous one. Thus, the form of inferred inputs is heavily influenced by model hyperparameters. In the case of LFADS, two critical choices are the prior distribution of the inferred inputs and the magnitude of the penalty for divergence from that prior.

**Figure 6.**
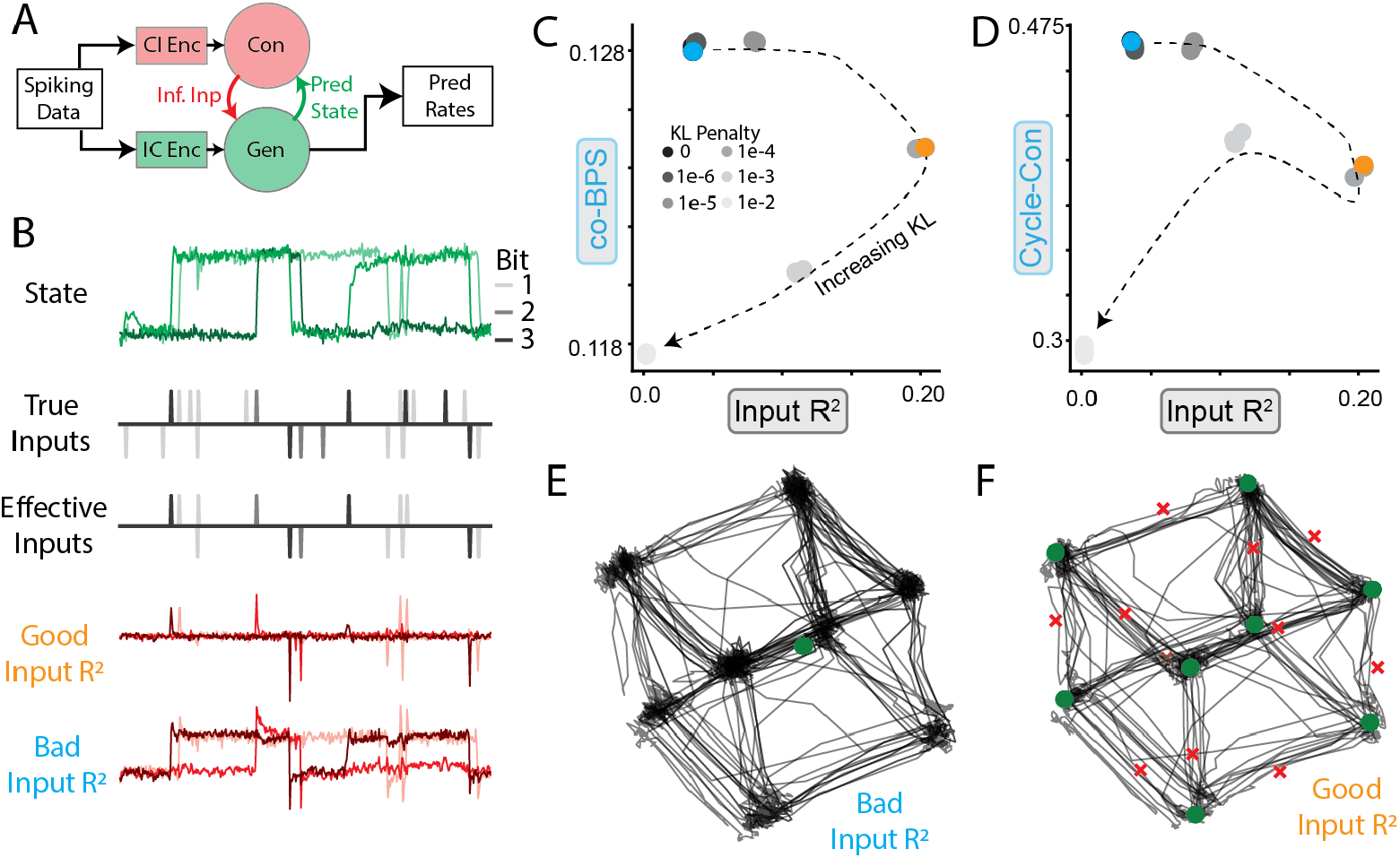
Inferred inputs affect inferred dynamics. **A**) Schematic of common input inference procedure for DD models. IC Enc, CI Enc denote Initial Condition encoder, Controller Input encoder, respectively. Ideally, the generator only models the intrinsic dynamics of the circuit and the controller only predicts inputs when inputs are actually present. **B.** Example activity from a single trial of 3BFF task. Shading indicates bit number. Top row: Output of TT-GRU. Rows 2-3: True and effective inputs (inputs that are expected to affect the state). Rows 4-5: Inferred inputs for two example models with “good” (orange) and “bad” (cyan) input inference. **C**) Scatter plot of co-BPS vs. Input *R*^2^ for models with increasing KL penalty (indicated by shading/ dashed line). Models depicted in B shown in orange/cyan. **D**) Same as C, except cycle-consistency vs. Input *R*^2^. **E**) Inferred FP architecture of model colored in cyan in (C). This model infers no FPs at the vertices of the cube, or any unstable FPs. Note that these are input-determined fixed points, computed under the relevant static inputs to the model (the dynamics, and hence the fixed point structure, are a function of the inputs), rather than fixed points of the autonomous (zero-input) system. **F**) Same as E, except for orange model in (C). Stable (unstable) FPs are depicted as green circles (red ×s). Compare to ground-truth FP structure in Fig. 3C.

To validate that using Input *R*^2^ to select hyperparameters leads to accurate inferred inputs and dynamics, we fit multiple LFADS models with different weights on the prior distribution of inferred inputs (Student’s *t*-distribution with *df* = 5, which promotes sparse inputs) to data from the 3BFF canonical dataset. We compared the inferred inputs to both the true inputs and the “effective” inputs (i.e. inputs that would change the sign of the bit, as opposed to redundantly indicating the current sign, Fig. 6B, top three rows). Some models inferred inputs that qualitatively matched the effective inputs, while other models’ inferred inputs resembled the state of the network.

Plotting the models’ co-BPS and cycle-consistency against the Input *R*^2^ (Fig. 6C,D) demonstrated that many models with high co-BPS and cycle-consistency had extremely poor input inference. As the KL penalty increased, Input *R*^2^ increased until the KL penalty became high enough that the model began to suffer from underfitting (Fig. 6C). Interestingly, we found that models with both high and low Input *R*^2^ could find inferred activity that was almost identical to the true system (Fig. 6E,F, black traces). However, fixed-point analysis revealed that only models with high Input *R*^2^ reproduced the expected FP structure (Fig. 6E,F, ∘, × markers), while those with low Input *R*^2^ failed to capture any of the relevant FPs. This suggests that the model with low KL penalty suffered from the failure mode of *dynamical misattribution*, in which the controller takes over the relevant dynamics from the generator (Fig. 1H).

Together, these results indicate that reconstruction and simplicity alone may be insufficient when input inference is considered. Reconstruction, simplicity, and input accuracy each provide a piece of the puzzle for selecting data-driven models that can accurately infer dynamics from recorded neural activity. However, we caution that the hyperparameters found to produce good Input *R*^2^ on this dataset are unlikely to work well on datasets with different properties; since Input *R*^2^ relies upon knowledge of the ground-truth, this presents a major unresolved challenge for the field of data-driven dynamical modeling. Multi-area recordings and perturbations will likely be essential for resolving the ambiguities associated with external input inference.

## 4 Discussion, Limitations, and Future Directions

The Computation-through-Dynamics Toolkit (CtDToolkit) represents a significant step towards evaluating goal-directed neural computations through neural dynamics modeling. By offering standardized datasets and interpretable metrics, CtDToolkit provides researchers with tools to validate data-driven (DD) models, even when both dynamics and inputs must be inferred. Its modular and extensible design invites contributions from the community, promoting continued evolution to address emerging challenges.

### 4.1 Related approaches

The Neural Latents Benchmark (NLB) [49] demonstrated the value of standardized datasets and a shared evaluation metric (co-BPS) for comparing latent-variable models of neural population activity, and it catalyzed substantial methodological progress in the field. A key distinction is that NLB evaluates models against held-out neural activity, whereas CtDToolkit’s synthetic datasets provide access to the ground-truth latent dynamics and inputs, enabling us to directly assess whether inferred dynamics match the true system rather than only whether neural activity is well reconstructed. We therefore view CtDToolkit as complementary to NLB: NLB grounds evaluation in real recordings, while CtDToolkit provides the ground-truth control needed to validate that reconstruction- and simplicity-based metrics actually track dynamical accuracy.

A related lesson from NLB is that presenting a single headline metric to optimize against can, over time, lead researchers to focus on incrementally improving the number rather than on understanding the computations the brain performs. This risk is an instance of Goodhart’s law—once a measure becomes a target, it can cease to be a good measure. We designed CtDToolkit to guard against this tendency by reporting a suite of complementary criteria (reconstruction, simplicity, and input accuracy) rather than a single score, so that no one metric can be optimized in isolation without exposing trade-offs that a single number would obscure.

NLB also taught us that users tended to focus on a small subset of “workhorse” datasets rather than engaging broadly with the full range of available benchmarks. This suggested that including an extensive collection of diverse datasets could potentially dilute efforts across many standards. Here, we prioritize a smaller, curated set of validated datasets to help the field consolidate around common standards of model evaluation, accepting that this approach may limit applicability to neural recordings with substantially different characteristics or computational demands. We hope that researchers will continue to expand the collection of datasets by submitting their tasks and models to CtDToolkit following the procedure described in Appendix F.2.

Selecting DD models that recover the true dynamics can be approached in complementary ways, and CtDToolkit deliberately commits to one: a *metric-based* approach that supplies interpretable metrics flagging whether a model has fallen into a characteristic failure mode (e.g., underfitting), which can then be used to compare and select among candidates. A complementary strategy instead seeks *consistent* solutions—training multiple models and keeping those whose inferred features agree, or identifying the minimum-complexity regime in which solutions become reproducible across independently trained models [34]. Both target the same goal, so we expect models favored by one to fare well under the other; we adopt the metric-based approach here, but encourage readers to consider consistency-based alternatives where appropriate.

More broadly, we see CtDToolkit as an opportunity to build bridges between the neural-dynamics modeling community and the broader field of dynamical systems reconstruction (DSR), much of which has developed largely outside of neuroscience [54, 45, 48, 57]—with insight flowing in both directions. From DSR, neural-dynamics modeling stands to inherit a mature body of experience validating inferred dynamics against ground truth using measures of *invariant* structure. These include geometric measures, such as state-space (Kullback–Leibler) divergence [29, 45] and Wasserstein distance between distributions of visited states [57], as well as temporal measures, such as power-spectrum or autocorrelation agreement [45, 48]. Such measures are typically computed in a shared, observation-derived space without requiring ground-truth latents or trial-by-trial alignment. Our Wasserstein distribution-matching metric brings this perspective into CtDToolkit, and incorporating the complementary temporal measures is a natural next step. In the other direction, the neural setting confronts DSR with challenges that its canonical (autonomous, non-neural) benchmarks largely lack—unknown external inputs, feedback, and explicit goal-directed computation—and CtDToolkit packages these as concrete, ground-truthed benchmark systems on which DSR methods can be stress-tested and extended. By offering a shared platform and common metrics, CtDToolkit can help the two communities exchange both tools and problems, which we see as a promising avenue for progress.

### 4.2 Limitations

Despite its strengths, CtDToolkit faces limitations rooted in our incomplete understanding of neural dynamics. First, current approaches assume that rate coding is sufficient to describe neural dynamics, potentially ignoring important intracellular and neuromodulatory processes; future extensions could incorporate temporal coding and diffuse neuromodulatory influences to test these assumptions and broaden the toolkit’s applicability. Second, our datasets and most of our metrics assume a linear observation model mapping latent dynamics to neural activity. While CtDToolkit includes the non-linear DSA and nl_cycle_con methods [56, 66], further work is needed to assess how deviations from linearity affect model performance and to develop additional metrics that are robust to these deviations [18, 38].

A further limitation is that all of our metrics are computed on inferred latent trajectories and firing rates rather than on the inferred flow field directly. Trajectory- and rate-based metrics can be confounded by observation noise and input-driven variability, whereas an error metric defined between the inferred and ground-truth flow fields would avoid these confounds. Such a comparison is not straightforward for the GRU-based TT architectures used here, but it is feasible for architectures whose vector field can be evaluated directly (e.g., low-rank RNNs). We therefore view direct comparison between inferred and ground-truth flow fields, enabled by appropriate TT model architectures, as a valuable direction for future work.

Input inference remains a major challenge. Although metrics like co-BPS and cycle-consistency estimate reconstruction and simplicity without ground-truth, CtDToolkit lacks an analogous metric for input accuracy. Simulated perturbations and new metrics for quantifying input inference accuracy could help address this issue. We consider this to be a major focus of future work in this area.

Looking forward, CtDToolkit’s extensibility offers pathways for addressing these limitations. Contributions such as richer datasets, novel metrics, and improved nonlinear methods can enhance the utility of the toolkit. With continued community collaboration, CtDToolkit can drive innovations that deepen our understanding of neural computation and help bridge the gap between models and biological systems. In summary, CtDToolkit not only provides a foundation for evaluating neural dynamics models but also a launchpad for broader investigations into neural computation. As the field advances, CtDToolkit has the potential to play a key role in shaping tools and frameworks that uncover fundamental principles of brain function.

## Author Contributions

**Table 1.**
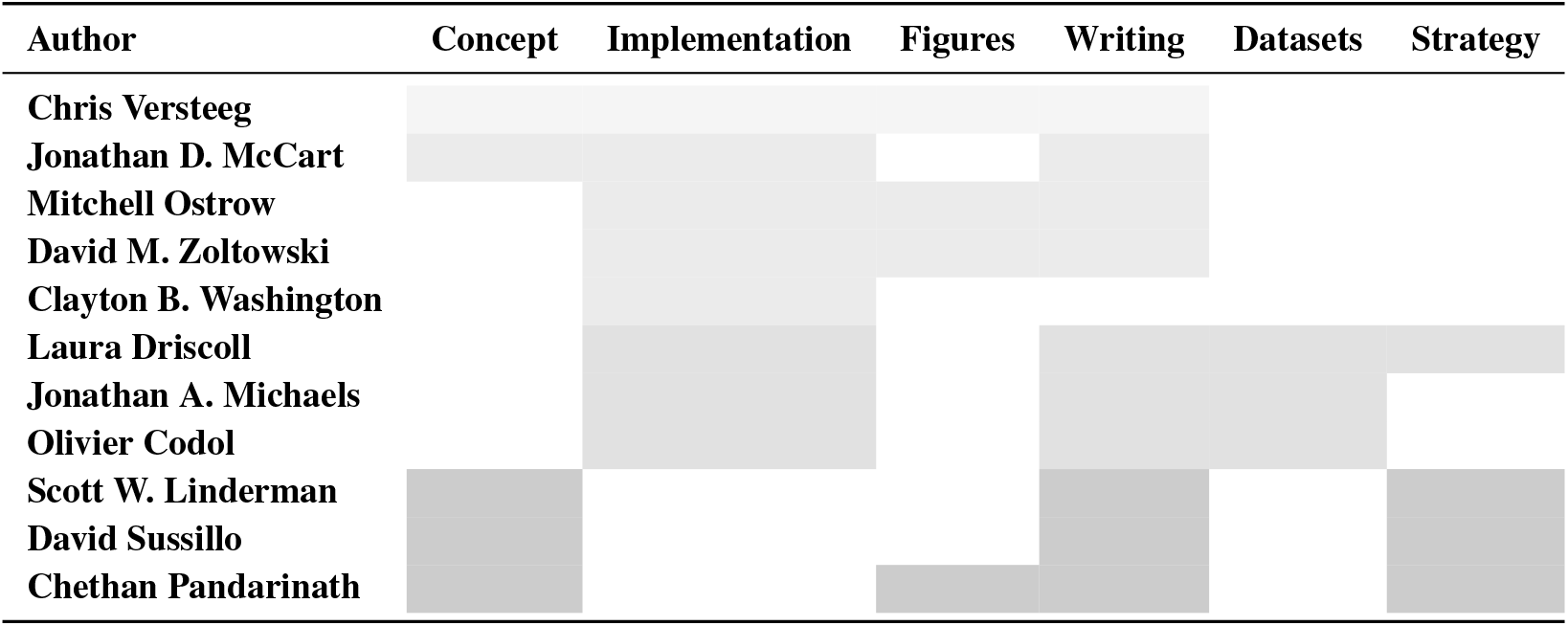
Author contributions. Shaded cells indicate areas of significant involvement. **Concept** includes ideation and experimental design; **Implementation** includes coding, model development, debugging, and/or website creation; **Figures** includes figure creation and data visualizations; **Writing** includes manuscript drafting, revising, and editing; **Datasets** includes guidance on synthetic dataset creation and critical external code packages; **Strategy** indicates broad scientific and conceptual guidance on overall project goals.

## Acknowledgements

We thank Srdjan Ostojic for helpful discussions and feedback on this project and Sophia Sanborn for her helpful PyTorch implementation of the MultiTask environment. We also thank Matthijs Pals for his insight into the PhaseCodedMemory task. This work was supported by NIH BRAIN/NIDA RF1 DA055667 and NIH NINDS/OD DP2 NS127291 (CP), and NIH NINDS 5F32MH132175 (CV); the Simons Foundation through the Simons-Emory International Consortium on Motor Control (CP, CV, JAM) and the Simons Collaboration on the Global Brain awards 543049 (DS), 697092 (SWL), the Transition to Independence Award 1155867 (LD); and the NSF GRFP 2141064 (MO). DMZ is supported by the Wu Tsai Neurosciences Institute. Additional support came from the FRQNT Strategic Clusters Program, Centre UNIQUE – Centre de recherche Neuro-IA du Québec (OC), and the Centre Interdisciplinaire de Recherche sur le Cerveau et l’Apprentissage (OC).

## Supporting Information Legends

**Tables**

**S1 Table. Parameters for the task-training and simulation datasets for 3BFF.**

**S2 Table. NoisyGRU model parameters (3BFF).**

**S3 Table. MultiTask phase length by trial type: [min, max] in bins.**

**S4 Table. Task-training dataset parameters (MultiTask).**

**S5 Table. NoisyGRU model parameters (MultiTask).**

**S6 Table. Task-training dataset parameters (RandomTarget): all timings are in bins (10 ms).**

**S7 Table. NoisyGRU model parameters (RandomTarget).**

**S8 Table. Parameters for the task-training and simulation datasets for PhaseCodedMemory.**

**S9 Table. NoisyGRU model parameters (PhaseCodedMemory).**

**S10 Table. Canonical dataset simulation parameters.**

**S11 Table. Hyperparameters for data-driven models (NODE-D) from Figure 5.**

**S12 Table. Hyperparameters for data-driven LFADS models in Figure 6.**

**S13 Table. Hyperparameters for the DSA fits and fixed-point finding in S8 Figure.**

**S14 Table. Hyperparameters for the NODE-based SAE latent-size sweeps in the compiled-metrics figures (NBFF, MultiTask, RandomTarget, ChaoticDelayedMatching).**

**S15 Table. Hyperparameters for the LFADS latent-size sweep on PhaseCodedMemory in the compiled-metrics figure.**

**Figures**

**S1 Figure. Schematic of task-training pipeline**. Here we show the general procedure for running task training with the structure of the objects that appear in the code, and their interactions, shown graphically. First, the data that define the inputs/outputs of a task must be generated. If the dataset has already been generated, it can be found in `/content/datasets/tt`; otherwise, the datamodule will call the method `generate_dataset()` from the TaskEnvironment. The task data will then be split into training and validation datasets by the `DataModule`. In the case of coupled environments that require feedback at each step (e.g. RandomTarget), the `TaskEnvironment` is also loaded. CtDToolkit uses the `TaskWrapper` to coordinate training and saving, so each of these objects are passed together to the `TaskWrapper` which runs the task-training pipeline described in Section <u>F.</u> When training is complete, the `DataModule`, `TaskEnvironment`, and task-trained model are saved to `/content/trained_models`.

**S2 Figure. Example trial for 3BFF**. Example input pulses, target states, and task-trained model outputs for one 3BFF trial.

**S3 Figure. Example trials for MultiTask**. We show one example trial for each task in MultiTask, organized in columns by trial category. Each trial has four subplots, with the task name indicated to the left of each row. Each subplot shows a 2D signal over time as a scatter plot, color coded by task phase. The left two plots indicate the inputs to stimulus modalities 1 and 2, respectively. The third plot indicates the model output on its two response channels. The rightmost plot indicates the target output.

**S4 Figure. PhaseCodedMemory task-trained model and simulated data. A**) Schematic of the PhaseCodedMemory task. The model is trained to generate an output oscillation shifted by 0.2*π* if an input stimulus is seen on input channel A (purple trace, Stim A) or by 1.2*π* if on input channel B (cyan trace, Stim B). Reference oscillation input is depicted as a dashed black line. **B**) Two example trials from a TT model performing the PhaseCodedMemory task. Trials differ only in the channel that received the input pulse. In both trials, the model receives the same sinusoidal reference oscillation. **C**) Single-unit activity of the TT network in the post-stimulus window is phase shifted relative to the reference oscillation for trials with different input stimulation. **D**) Top three PCs of the latent activity in the post-stimulus window of the PhaseCodedMemory TT model. **E**) Ground-truth firing rates and simulated spiking activity for the synthetic PhaseCodedMemory dataset. **F-G**) Example DD model activity and inferred single-unit responses consistent with a rate-coding rather than phase-coding strategy. **H**) Histogram of normalized absolute differences in activity between stimulus conditions, illustrating that the DD model responses are more dissimilar across stimulus conditions than the TT model responses.

**S5 Figure. ChaoticDelayedMatching task-trained model and chaotic signatures**. Task structure, an example data-driven (DD-NODE) model fit, and the chaotic signatures of the task-trained dynamics for the ChaoticDelayedMatching task. **A)** Delayed non-match-to-sample trial structure with example cue inputs and target output. **B)** Task-trained model output tracks the target during the response window, including for an ensemble of perturbed initial conditions. **C)** Example DD-NODE fit: simulated spikes, ground-truth and predicted firing rates, example rate traces, and inferred latent trajectory. **D)** Initial-condition perturbations diverge approximately exponentially in latent space while the output remains bounded.

**S6 Figure. Neural data simulation pipeline**. We use a separate directory for task data used to generate simulated neural data (`sim`) compared to running task training (`tt`). The task data are loaded into a `DataModule` as a means of curating the trials passed through the trained model to obtain hidden-unit activations. In coupled environments, the `TaskEnvironment` is loaded to enable feedback at each step. Once all trials have been processed by the task-trained model, the hidden-unit activations are used by the Neural Data Simulator to produce simulated neural spiking activity. The simulated data are saved to `dd`, where they can be accessed later when running the data-training pipeline.

**S7 Figure. High-level schematic of data-training pipeline**. The simulated neural data file is loaded from `dd` by the `prepare_data()` and `setup()` methods of the `DataModule`, and packaged into training and validation dataloaders. These dataloaders are used to train the data-driven model, an implementation of a LightningModule. The Trainer also accepts user-defined callbacks and loggers. After training is complete, the data-driven model and the corresponding datamodule are saved to `/content/trained_models`.

**S8 Figure. DSA and fixed-point structure for NODE and GRU models across latent size (3BFF). A)** DSA dissimilarity matrices comparing the ground-truth 3BFF TT model and data-driven NODE and GRU models across latent sizes, on a shared color scale. The boxed bottom row marks TT-vs-model dissimilarities, excluding the TT-vs-TT self-comparison. **B)** DSA dissimilarity to the TT model versus latent size for NODE and GRU models. **C)** Inferred fixed points over latent trajectories for the 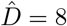 and 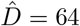 NODE and GRU models. All model latents and fixed points are linearly projected onto the ground-truth 3-bit task state for a common orientation, and fixed points are computed under zero external input.

**S9 Figure. Nonlinear cycle-consistency metric for DD-NODE models trained on the NBFF canonical dataset. A)** `nl_cycle_con` curves (*R*^2^ vs. injected noise standard deviation) for DD-NODE models across latent sizes and random seeds. **B)** Scatter plot comparing nonlinear cycle consistency, evaluated at *σ* = 0.05, against linear cycle consistency. **C)** Training diagnostics for the learned inverse MLP used by the metric: training and validation MSE loss versus epoch. Training and validation losses track closely across all latent sizes, indicating that the inverse mapping is not overfit.

**S10 Figure. Validation of the CtDToolkit Lyapunov-spectrum estimator on canonical chaotic systems**. The full Lyapunov spectrum was estimated for the standard Lorenz and Rössler systems by integrating the vector field with a fourth-order Runge–Kutta scheme, building discrete-time Jacobians by finite differences, and applying `compute_lyapunov_spectrum`. Points and error bars show the mean ± s.d. of the estimated exponents across trajectories; red crosses show literature reference values. Panel titles report the maximal exponent *λ*_max_ and the sum ∑_*i*_ *λ*_*i*_.

**S11 Figure. Compiled metrics for DD-NODE models trained on the NBFF canonical dataset. A)** Reconstruction metrics for DD-NODEs fit to the NBFF canonical dataset: Rate *R*^2^ and co-BPS. **B)** Simplicity metrics for the same models: State *R*^2^ and cycle consistency. **C)** Dynamical-systems-inspired metrics: Wasserstein distance and estimated maximal Lyapunov exponent.

**S12 Figure. Compiled metrics for DD-NODE models trained on the MultiTask canonical dataset. A)** Reconstruction metrics for DD-NODEs fit to the MultiTask canonical dataset: Rate *R*^2^ and co-BPS. **B)** Simplicity metrics for the same models: State *R*^2^ and cycle consistency. **C)** Dynamical-systems-inspired metrics: Wasserstein distance and estimated maximal Lyapunov exponent.

**S13 Figure. Compiled metrics for DD-NODE models trained on the RandomTarget canonical dataset. A)** Reconstruction metrics for DD-NODEs fit to the RandomTarget canonical dataset: Rate *R*^2^ and co-BPS. **B)** Simplicity metrics for the same models: State *R*^2^ and cycle consistency. **C)** Dynamical-systems-inspired metrics: Wasserstein distance and estimated maximal Lyapunov exponent.

**S14 Figure. Compiled metrics for LFADS models trained on the PhaseCodedMemory canonical dataset. A)** Reconstruction metrics for LFADS models fit to the PhaseCodedMemory canonical dataset: Rate *R*^2^ and co-BPS. **B)** Simplicity metrics for the same models: State *R*^2^ and cycle consistency. **C)** Dynamical-systems-inspired metrics: Wasserstein distance and estimated maximal Lyapunov exponent.

**S15 Figure. Compiled metrics for DD-NODE models trained on the ChaoticDelayedMatching canonical dataset. (A)** Reconstruction metrics for DD-NODEs fit to the ChaoticDelayedMatching canonical dataset: Rate *R*^2^ and co-BPS. **(B)** Simplicity metrics for the same models: State *R*^2^ and cycle consistency. **C)** Dynamical-systems-inspired metrics: Wasserstein distance and estimated maximal Lyapunov exponent.

**S16 Figure. Data-driven model validation on PhaseCodedMemory**. Each row corresponds to one model: the ground-truth TT system and LFADS DD models of increasing latent dimensionality. Columns show firing-rate rasters, top PCs of response-period activity in firing-rate space, and top PCs of response-period activity in latent space. Reconstruction improves with latent size while State *R*^2^ degrades, and the DD models do not recover the qualitative phase-coding geometry of the TT model.

**S17 Figure. Objects for user-defined tasks and TT models**. Schematic of the core objects used in the task-training pipeline: TaskEnvironment, TT model, TaskWrapper, TTDatamodule, and Simulator.

## Appendix A Computation-through-Dynamics Toolkit

We envision two classes of CtDToolkit users:

1. **DD model/metric developer:** This group will train their data-driven models on the canonical datasets and use the metrics provided to assess the model performance. We expect this user group to be primarily interested in adding their own data-driven models and metrics to the code-base, and less interested in modifying the existing tasks or simulating their own neural data. This group is referred to as “DD-Developers”.
2. **TT model developer:** This group is primarily interested in adding their own tasks to the CtDToolkit and/or simulating neural data with different characteristics (noise models, observation models, etc.). We have provided a template class that these users can follow to incorporate new tasks into CtD, and describe the resources available to these users in Section F.1.

As described in Section 2 and shown in Figure 2A, CtD has three major components:

1. **Generating simulated neural datasets:** We provide the task environments and logic necessary to produce datasets from each of the tasks in CtD. We also include access to pre-trained models so that users can directly simulate neural data corresponding to these tasks.
2. **Training data-driven models:** We provide a pipeline for training neural dynamics models on simulated neural data produced by a task-trained model.
3. **Comparing model performance:** We provide several metrics for analyzing and comparing neural dynamics model performance, as well as the supporting code for easily logging and visualizing such results.

### A.1 Overview of DD-Developer process flow

We provide five canonical datasets for DD-Developers. Each synthetic dataset was generated from a TT model trained to perform one of the included tasks. DD-Developers can get started with training their models on the canonical datasets by following a few simple steps:

1. **Clone the GitHub repository:** Navigate to [our GitHub page](https://github.com/snel-repo/ComputationThroughDynamicsToolkit) and clone the repository. This repository includes task-training and data-training pipelines, comparison metrics, and five pre-trained models, which are used to generate the canonical datasets.
2. **Set up the environment:** Follow the instructions in the README to set up the environment for the CtD Toolkit.
3. **Generate the canonical datasets:** Users can generate the canonical datasets using pre-trained models for each task by running the script `/examples/gen_datasets.py`. This script generates the canonical datasets by 1) generating inputs and outputs from a saved task-environment, 2) running the pre-trained TT model on this dataset, 3) simulating neural activity from the hidden activity of the pre-trained model (saved in `/content/datasets/dd`). More information on the task-environments and task-training pipeline can be found in Section B.
4. **Train models:** Use `/examples/run_data_training.py` to train models on the canonical datasets. Users can choose the model class (LFADS, SAE), the canonical dataset (NBFF, MultiTask, RandomTarget, PhaseCodedMemory), and the input mode (supplied or inferred). Running this script without changing any parameters will start a run that will replicate the results from Figure 5. More information on data-driven models can be found in Section D.
5. **Compare trained models:** Once the models are done training, users can load the saved models into Analysis_DD objects (`/ctd/comparison/analysis/dd`), which can be compiled in a Comparison object (`/ctd/comparison/comparison.py`). The Comparison object provides useful tools for visualizing and computing metrics across trained models. More information on the available comparisons can be found in Section E, and a useful example notebook can be found at `/examples/compare_tt_dd_models.ipynb`.

## APPENDIX B Canonical Datasets and Task Training

In this section, we give details on the task environments, models and hyperparameters that were used to generate the pre-trained models for each task. We first describe the high-level Task Environment structure, and then describe how it was used for each canonical dataset. Information on using the pre-trained models to generate synthetic datasets is given in Section C. We summarize the training pipeline in Figure S1.

**Figure S1.**
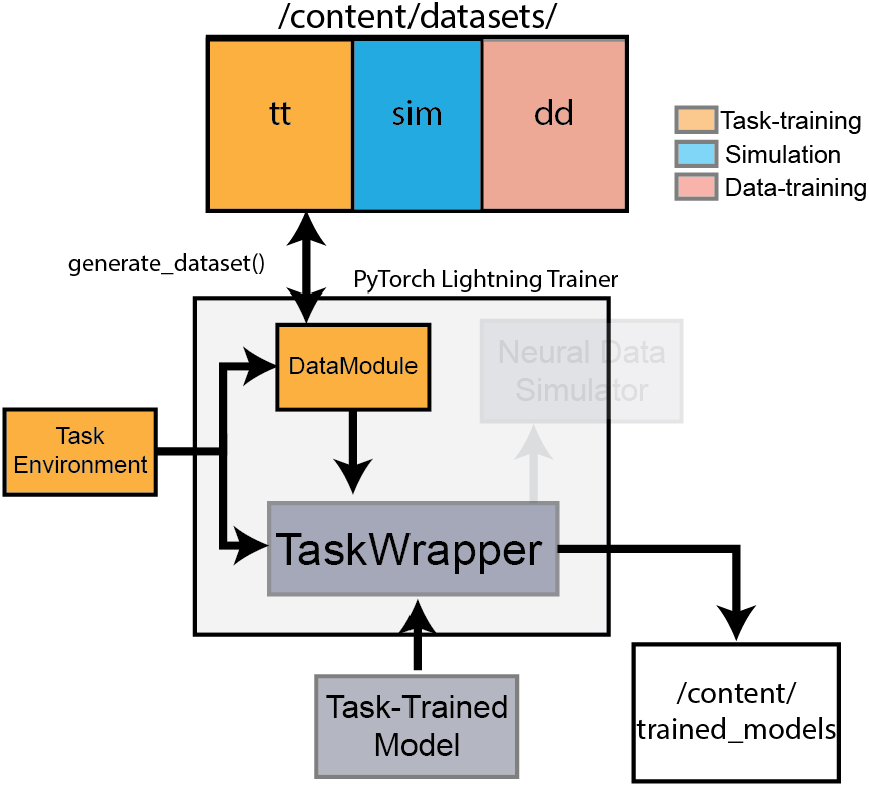
Schematic of task-training pipeline. Here we show the general procedure for running task training with the structure of the objects that appear in the code, and their interactions, shown graphically. First, the data that define the inputs/outputs of a task must be generated. If the dataset has already been generated, it can be found in `/content/datasets/tt`; otherwise, the datamodule will call the method `generate_dataset()` from the TaskEnvironment. The task data will then be split into training and validation datasets by the `DataModule`. In the case of coupled environments that require feedback at each step (e.g. RandomTarget), the `TaskEnvironment` is also loaded. CtDToolkit uses the `TaskWrapper` to coordinate training and saving, so each of these objects are passed together to the `TaskWrapper` which runs the task-training pipeline described in Section F. When training is complete, the `DataModule`, `TaskEnvironment` and task-trained model are saved to `/content/trained_models`.

### B.1 Task Environments

We use the TaskEnvironment structure to generate the task-training dataset for the task-trained model. TaskEnvironments inherit from the Gymnasium.Environment class. Each TaskEnvironment defines the logic of a given task and implements the function `generate_dataset()`, which returns a dictionary that contains the inputs, outputs, and other information necessary to train a model to perform the task. For TT model developers who wish to modify or add TaskEnvironments, we describe more requirements for TaskEnvironments in Section F.1.

The `generate_dataset()` function is the heart of the TaskEnvironment. This function returns a dictionary with the following entries:

- `ics`: (shape: `n_samples` × `D`). The initial conditions of the task environment (e.g., joint angles at trial start, the state of the bits in 3BFF).
- `inputs`: (shape: `n_samples` × `timesteps` × `n_inputs_pre`). The inputs to the model that can be pre-computed (e.g., inputs pulses for 3BFF).
- `inputs_to_env`: (shape: `n_samples` × `timesteps` × `n_ext_inputs`). Any inputs that are applied to the environment rather than passed directly to the TT model. The bump perturbations in the `RandomTarget` task are the only examples of this type of input in the provided datasets.
- `targets`: (shape: `n_samples` × `timesteps` × `n_targets`). The target values for the TT model (e.g., the true state of the 3BFF).
- `true_inputs`: (shape: `n_samples` × `timesteps` × `n_inputs`). This contains the noiseless ground truth inputs for visualization and training.
- `conds`: (shape: `n_samples` × `1`). Condition indices for each trial. Used to re-weight the loss contributions of different trial types and for batching during training (used only in MultiTask).

Throughout, time is discretized into fixed-width *bins*: all per-trial quantities (inputs, outputs, latent states, and spikes) and all timings reported below are expressed in these bins, and the number of bins per trial is set by each task’s `n_timesteps` parameter. Importantly, the task-trained (TT) model, the neural simulator, and the data-driven (DD) models all operate on this same binning—the simulator emits spikes on the bins of the TT latent trajectory, and DD models are fit to those spikes without any rebinning or resampling. The TT and DD models therefore use an identical time step within a given dataset. The physical duration of a bin is task-dependent. For tasks tied to a physical process we set an explicit bin width: 10 ms for RandomTarget (matching the MotorNet effector integration step), 20 ms for MultiTask, 2 ms for PhaseCodedMemory (a 0.8 s trial divided into 400 bins), and 1 ms for ChaoticDelayedMatching (so that the leaky-integration time constant *τ* = 30 ms corresponds to the model’s leak factor *α* = d*t/τ* = 1*/*30). For 3BFF, the underlying dynamical system is scale-free, so time is expressed in dimensionless bins (equivalently, arbitrary time steps) with no physical unit.

Each TaskEnvironment also includes the loss function that is used to train models to perform the task. We describe the loss function for each of the canonical datasets in more detail below.

#### B.1.1 Three-Bit Flip-Flop (3BFF)

##### General Description

The simplest of the canonical datasets is the 3BFF dataset, which is a 3 bit memory task described first in [15]. The task takes inputs **u**_*t*_ ∈ ℝ^3^ representing pulse signals on three channels and produces outputs **x**_*t*_ ∈ ℝ^3^ corresponding to the binary memory states of each channel. Our implementation can be found in `ctd/task_modeling/task_env/task_env.py` in the NBitFlipFlop class. Below, we describe the environment parameters and loss function:

##### Environment Parameters

The NBitFlipFlop class takes the parameters described below. Values used in the canonical 3BFF dataset are given in Table S1.

- `n_timesteps`: The length of each trial in bins.
- `noise`: Standard deviation of Gaussian noise to be added to the inputs. This noise is pre-computed and the same every time a given trial is presented.
- `n`: The number of bits in the NBitFlipFlop (Default: 3).
- `switch_prob`: The probability of an input pulse on a given timestep.
- `transition_blind`: The number of bins after an input pulse that the loss is zeroed out (described in *Loss function* below).

**Table S1.** Parameters for the task-training and simulation datasets for 3BFF.

| Task parameter | 3BFF (train) | 3BFF (sim) |
| --- | --- | --- |
| `n_trials` | 1000 | 1000 |
| `n_timesteps` | 500 | 500 |
| `noise` | 0.15 | 0.15 |
| `n` | 3 | 3 |
| `switch_prob` | 0.01 | 0.01 |
| `transition_blind` | 4 | NA |

##### Trial Structure

Trials begin with all states set to zero. Each timestep has a small probability of a positive or negative input pulse on each channel. Positive inputs push the state from zero to one, negative inputs push the state from one to zero, while positive (negative) inputs have no effect when the state is one (zero). We show an example trial of 3BFF in Figure S2.

**Figure S2.**
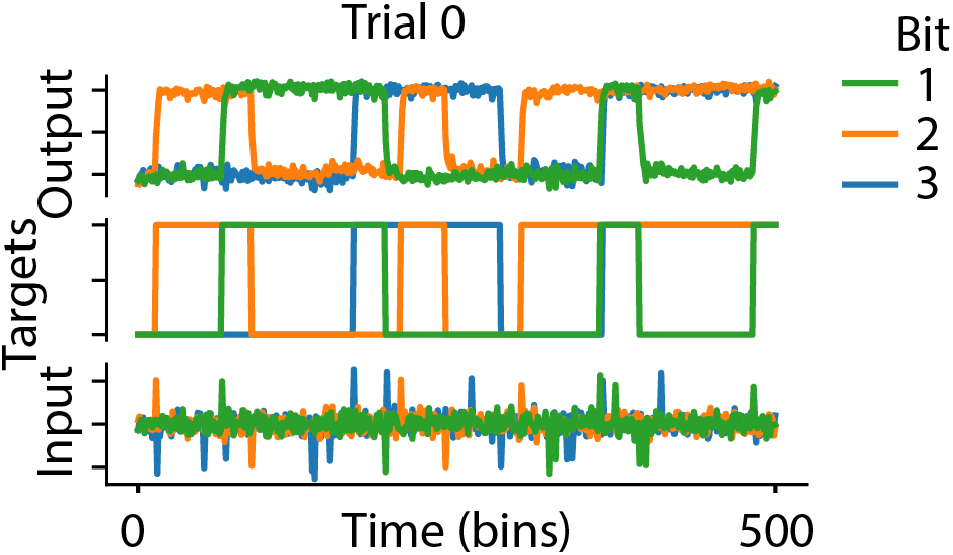
Example trial for 3BFF.

##### Loss Function

During training, we penalize the mean squared error between the model output and the desired output. To avoid interference from initial transient effects, we mask the losses during the first 5 bins of each trial. Additionally, we apply a masking period (default: 4 bins, defined by `transition_blind`) following each input pulse. We found that this masking helped the model learn smoother dynamics that cleanly transitioned from one state to another without large discontinuous jumps in the state. See `/ctd/task_modeling/task_env/loss_func.py` for full implementation details.

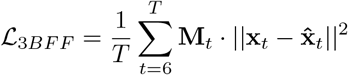

where **x**_*t*_ is the target (desired) output and 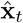 is the model-predicted output at time *t, T* is the trial length in bins, and **M**_*t*_ is the masking function, where *t*_*pulse*_ denotes the time (in bins) of an input pulse:

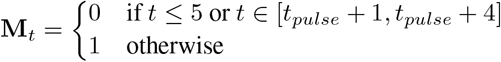

**Table S2.** NoisyGRU model parameters (3BFF)

| Hyperparameters | 3BFF |
| --- | --- |
| learning_rate | 1e-3 |
| max_epochs | 500 |
| input_size | 3 |
| output_size | 3 |
| latent_size | 64 |
| intrinsic noise | 0.01 |
| latent_ic_var | 0.05 |

##### Task-Trained Model (NoisyGRU)

For the canonical 3BFF dataset, we used a modified Gated Recurrent Unit (GRU) network called a NoisyGRU. NoisyGRU adds a configurable intrinsic noise to the state at each time step and to the initial hidden state of the network on each trial. The noise at each time step was drawn from a standard normal distribution with mean zero and isotropic variance of 0.01, while the initial conditions were drawn from a standard normal with mean *µ* and variance of 0.05, where *µ* was a trainable parameter. This ensured that the model learned to stably encode memory regardless of semi-random initial conditions. Full hyperparameters are in Table S2. We provide the NoisyGRU model, fully-trained on 3BFF, in the `content/trained_models/task-trained/tt_3bff/` folder of the CtDToolkit repository.

The full state-update equations of the NoisyGRU are:

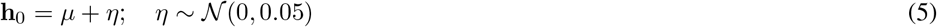

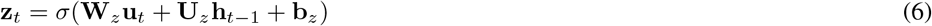

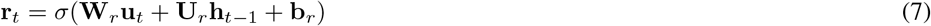

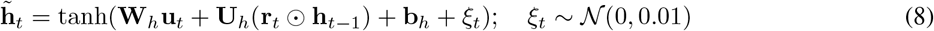

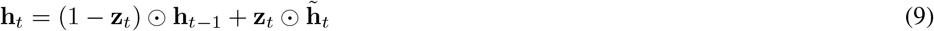

Where **u**_*t*_ is the task input at time step *t*, **h**_0_ is the initial hidden state with noise, **z**_*t*_ is the update gate at time step *t*, **r**_*t*_ is the reset gate at time step *t*, 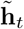 is the candidate hidden state at time step *t*, **h**_*t*_ is the final hidden state at time step *t, η* and *ξ*_*t*_ are the noise terms drawn from normal distributions, *σ*(·) is the sigmoid function, and ⊙ represents the element-wise (Hadamard) product. The trainable parameters are the input weight matrices **W**_*z*_, **W**_*r*_, **W**_*h*_, the recurrent weight matrices **U**_*z*_, **U**_*r*_, **U**_*h*_, the gate biases **b**_*z*_, **b**_*r*_, **b**_*h*_, and the mean initial state *µ*.

#### B.1.2 MultiTask

##### General Description

Our dataset of “intermediate” difficulty was generated using the previously described MultiTask task [33, 46], which is composed of 15 cognitive tasks that have overlapping task requirements (e.g., memory, directional responses, multiple stimulus modalities). To perform MultiTask, a model takes inputs **u**_*t*_ ∈ ℝ^20^ (1 fixation input, 2 inputs for stimulus modality 1, 2 inputs for stimulus modality 2, and 15 one-hot task inputs) and produces outputs 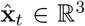 (1 fixation, 2 response channels). The tasks can be broken down into three categories: *response, decision making*, and *matching*.

##### Response

During Response tasks, inputs were received on the 2 input dimensions associated with stimulus modality 1, and the model was asked to respond by producing output either in the direction of the stimulus (“Pro”) or in the opposite direction (“Anti”). Different tasks required that the model respond at different points during the trial; for “React”, the model was trained to respond immediately upon receiving the stimulus. For “Delay”, the model was trained to respond after a short delay period in which the model continues to see the stimulus. For “Memory”, the model was trained to respond after a short period when the stimulus was turned off upon receiving a go-cue. With “Pro/Anti” versions of the “React/Delay/Memory” tasks, there are 6 total tasks in the Response category.

##### Decision Making

Decision Making tasks were designed to emulate systems that make decisions about the content of noisy inputs. In one version of the task (“IntMod”), we presented two stimuli to a single input modality, and the model was trained to move in the direction of the larger magnitude stimulus. In a second version (“ContextInt”), we presented stimuli to both modalities in each stimulation period, and models were trained to ignore one of the modalities based on the rule input. In the final version of the task, the model was trained to incorporate information from both modalities equally before deciding which was larger (“MultiModal”). With “IntMod1”, “IntMod2”, “ContextInt1”, “ContextInt2”, and “ContextIntMultimodal”, there were a total of 5 decision making tasks included in MultiTask.

##### Matching

The final class of tasks are Matching tasks. In two of these tasks, the model must produce output that indicates whether consecutive stimuli provided to the first input modality were the same direction (“Match”) or opposite direction (“Non-Match”) of one another, responding to the second stimulus location if so. The final two Matching tasks (“MatchCat”, “MatchCatAnti”) required the model to respond only if both stimuli took on values in the range [0, *π*) or [*π*, 2*π*).

We show an example trial of each task in Figure S3, grouped by the task type. Lengths of each phase of the tasks are drawn from a min and max length shown in Table S3. Additional implementation details are drawn from [46]. Full task code can be found at `/ctd/task_modeling/task_env/multitask.py`.

**Table S3.** Task phase length by trial type: [min, max] in bins.

| Trial Type | Context | Stim1 | Mem1 | Stim2 | Response |
| --- | --- | --- | --- | --- | --- |
| Delay | [15, 35] | [10, 75] | [0, 0] | [0, 0] | [15, 35] |
| Memory | [15, 35] | [10, 80] | [10, 80] | [0, 0] | [15, 35] |
| React | [25, 125] | [0, 0] | [0, 0] | [0, 0] | [15, 85] |
| Decision | [10, 30] | [10, 80] | [10, 80] | [10, 80] | [15, 35] |
| Match | [10, 30] | [10, 80] | [10, 80] | [0, 0] | [15, 35] |

##### Environment Parameters

The MultiTask class takes the parameters described below. The values for each parameter used when generating the canonical Multitask dataset are given in Table S4.

- `n_timesteps`: The maximum length across trials in bins.
- `noise`: Standard deviation of Gaussian noise to be added to the inputs.
- `num_targets`: The number of targets in the task (distributed uniformly on the circumference of the unit circle.)

**Table S4.** Task-training dataset parameters (MultiTask)

| Task parameter | MultiTask (train) | MultiTask (sim) |
| --- | --- | --- |
| `n_trials` (per task) | 1000 | 500 |
| `n_timesteps` | 320 | 320 |
| `noise` | 0.3 | 0.3 |
| `num_targets` | 128 | 32 |

Of note, all trials were 320 bins long (the longest possible trial given the lengths in Table S4). When feeding trials to our task-trained model, we simulated for the full 320 bins, then truncated the trials to the appropriate length for all subsequent loss computation and analysis.

**Figure S3.**
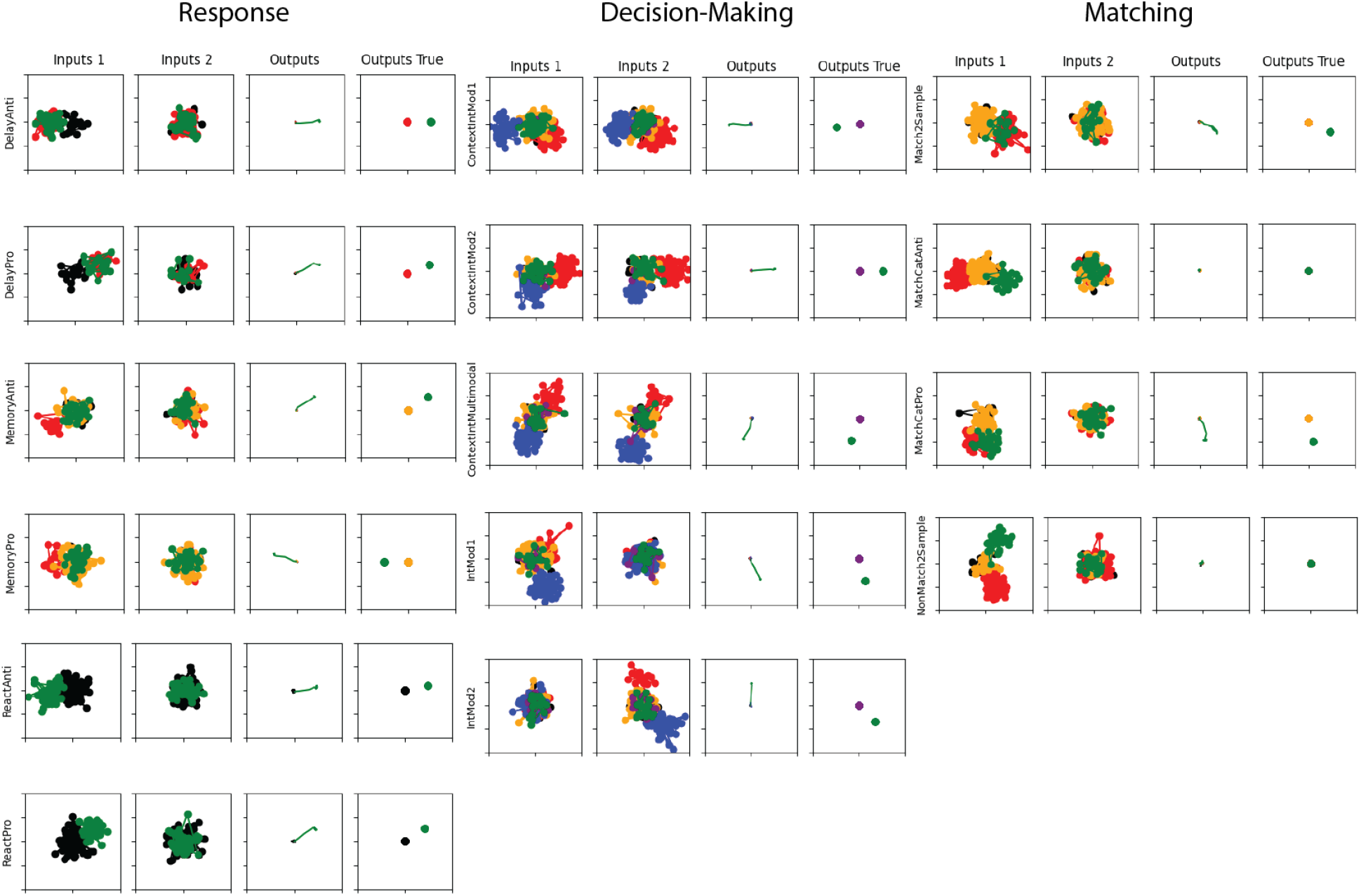
Example trials for MultiTask We show one example trial for each task in MultiTask, organized in columns by the trial category. Each trial has four subplots, with the task name indicated to the left of each row. Each subplot shows a 2D signal over time as a scatter plot, color coded by the task phase. Left two plots indicate the inputs to stimulus modality 1,2, respectively. Third plot indicates the model output on its two response channels. Rightmost plot indicates the targeted output.

##### Trial Structure

Each MultiTask trial proceeds through up to five epochs—a context (fixation) period, a first stimulus (Stim1), a memory delay (Mem1), a second stimulus (Stim2), and a response period—although which epochs are present and their durations depend on the task type. The duration of each epoch is drawn per trial from the [min, max] ranges in Table S3, where a zero range indicates that the epoch is absent for that task type.

##### Loss Function

The MultiTask loss function was to minimize the MSE of the task-trained model outputs against the desired outputs. Because trials were of variable length, we masked the MSE loss to only include time-bins that are within each trial’s total length (i.e., the sum of the randomly drawn intervals from Table S3). We also masked the loss to exclude contributions from the first 5 bins of each trial, and the first 5 bins after the response period begins. Additionally, we re-weighted the loss contribution of the response period so that performance during this period was weighted 5 times more than the rest of the trial. This was done in accordance with [46] to help with training speed and performance.

We computed the performance of our task-trained models using the criteria described in [46], which finds the percentage of trials where the model responds in the appropriate direction (i.e. the angle was correct to within an error of *< π/*10 radians), or on trials without a desired response, that the model does not respond. As with the previous work, we found that our TT models consistently achieved performance that exceeded 80% success rate on the validation dataset for all tasks.

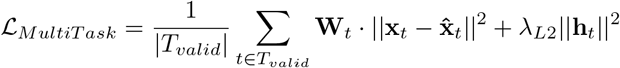

where:

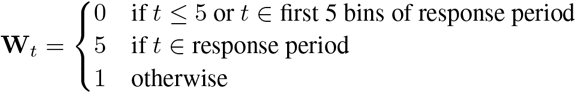

and *T*_*valid*_ includes only time-bins within each trial’s actual length, **h**_*t*_ is the model’s hidden activity, and *λ*_*L*2_ = 10^−6^.

##### Task-Trained Model (NoisyGRU)

For our canonical MultiTask dataset, we used a NoisyGRU (described in Section B.1.1) with the hyperparameters shown in Table S5. Training batches were composed of trials from only one task. We also added a loss term that penalized the squared hidden activity that pressured activity to live near the origin, leading to simpler fixed-point structures (similar to a loss contribution used in [46]). Relevant hyperparameters for the canonical Task-trained MultiTask model are given in Table S5.

**Table S5.** NoisyGRU model parameters (MultiTask)

| Hyperparameters | MultiTask |
| --- | --- |
| learning_rate | 1e-3 |
| max_epochs | 500 |
| input_size | 20 |
| output_size | 3 |
| latent_size | 128 |
| intrinsic noise | 0.01 |
| ic_noise | 0.05 |
| hidden_L2_penalty | 1e-6 |

#### B.1.3 RandomTarget

##### General Description

In the RandomTarget task, a TT model is trained to control a 2 degree-of-freedom arm model actuated by 6 Hill-type muscles (analogous to pectoralis, deltoid, triceps long, triceps lat, biceps, and brachialis). By controlling the muscle activations **a**_*t*_ ∈ ℝ^6^ of the arm model, the TT model must move the effector endpoint to various target locations sampled randomly from within the arm’s range of motion. TT models trained on this task receive inputs **u**_*t*_ ∈ ℝ^17^ consisting of both sensory (muscle lengths and velocities, hand endpoint position; **u**^*sens*^ ∈ ℝ^14^) and contextual (target position, go cue; 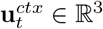) information.

**Figure S4.**
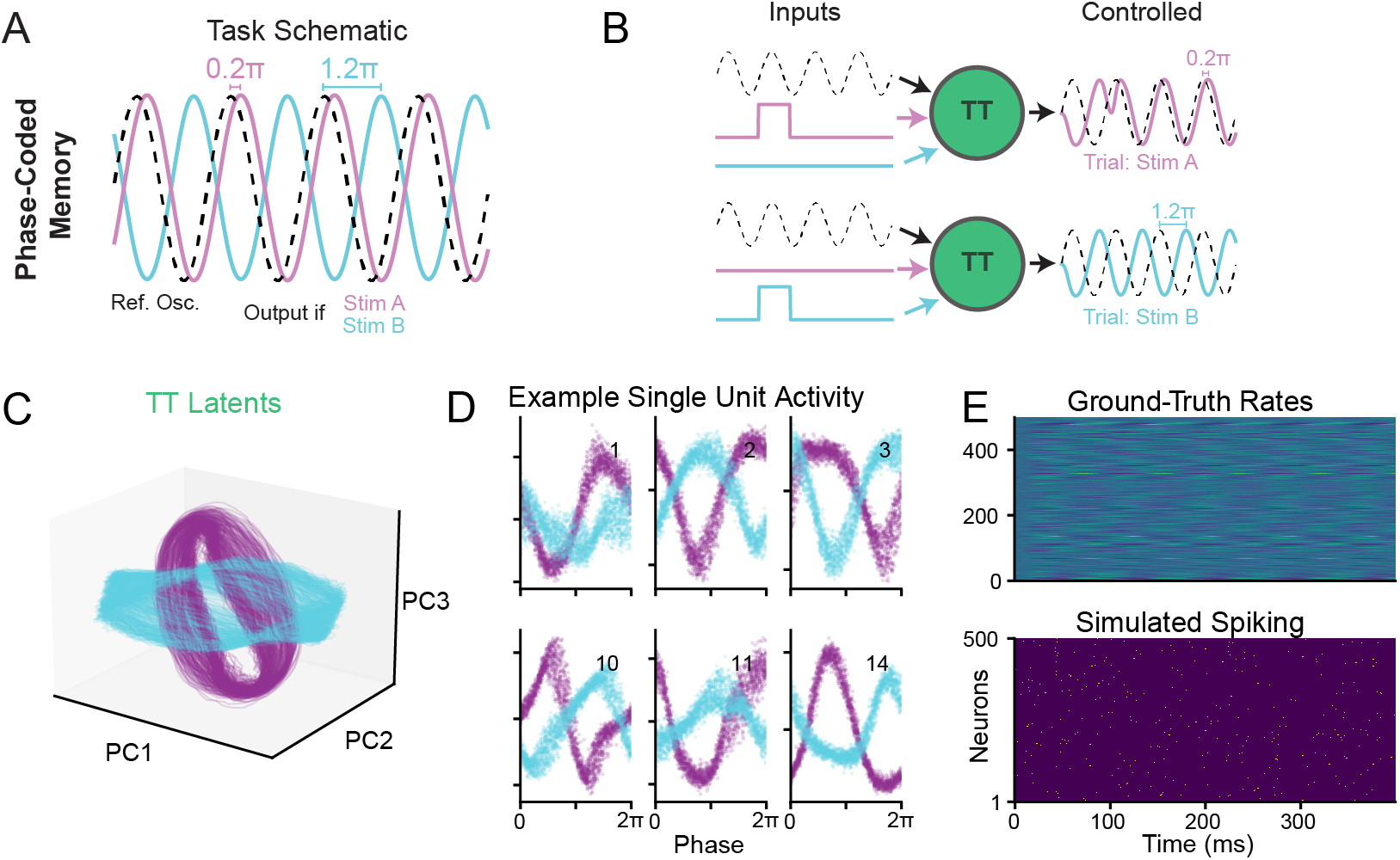
PhaseCodedMemory Task-Trained Model and Simulated Data. **A**) Schematic of PhaseCodedMemory Task. Model is trained to generate an output oscillation shifted by 0.2*π* if an input stimulus is seen on input channel A (purple trace, Stim A) or by 1.2*π* if on input channel B (cyan trace, Stim B). Reference oscillation input is depicted as dashed black line. **B**. Two example trials from a TT model performing the PhaseCodedMemory task. Trials differ only in the channel that received the input pulse (Top row = Stim A, bottom row = Stim B). In both trials, model receives the same sinusoidal reference oscillation (dashed black line). Trained model outputs reflect the desired encoding of the input stimulus identity. **C**) Top three PCs of the latent activity in the post-stimulus window of the PhaseCodedMemory TT model (plotted w/ noiseless inputs for clarity). As in [62], the two rings occupy overlapping regions of state space and are primarily rotated, rather than translated, relative to each other in the high-variance PC dimensions—consistent with a phase-coding strategy. **D**) As in [62], single unit activity of the TT network in the post-stimulus window has activity that is phase-shifted relative to the reference oscillation for trials with different input stimulation. For each panel, x-axis indicates the phase of the reference oscillation, y-axis indicates the activity of the hidden unit identified by the inset label. Stim A trials are purple, Stim B trials are cyan. **E**) Top: ground-truth firing rates of 500 simulated neurons, scaled to have mean FR of 2 Hz, on the scale of typical hippocampus CA1 neurons. Bottom: simulated spiking from Poisson neurons with rates given by the ground-truth above. An example data-driven model fit and validation on this dataset is shown in Supp. Fig. S16.

##### Environment Parameters

We chose parameters similar to those used in [61] for the two-link arm model. We summarize the parameters in Table S6. So that the model couldn’t anticipate the go cue and reach prior to the end of the delay period, 20% of training trials were “catch” trials in which the go-cue was never provided. During training the timing of the target presentation and go cue were random (target presentation occurred sometime in an interval of [100, 300] ms after trial start and the go cue occurred randomly between 50 ms after target presentation and the end of the trial, which lasted a total of 1.55 seconds). When simulating the neural data, the target onset and go-cue were consistent across trials (300 and 700 ms after the start of the trial, respectively), but bump timing was still random. For consistency with typical neural datasets of this type, our simulated data had consistent timings so that the go-cue occurred at a consistent alignment point rather than at a random point in the trial. When we simulated the data, we trimmed the first 5 bins of each trial to ensure that the activity of the network was stable at the starting location before the reaching behavior began.

We briefly describe important parameters of the RandomTarget task here:

- `n_timesteps`: The length of each trial in bins.
- `action_noise`: Standard deviation of Gaussian noise added to the actions output by the model (i.e., motor noise).
- `proprioception_noise`: Standard deviation of Gaussian noise added to the proprioceptive feedback (muscle lengths and velocities)
- `vision_noise`: Standard deviation of Gaussian noise added to the visual feedback (endpoint position)
- `context_input_noise`: Standard deviation of Gaussian noise added to the context inputs (target position and go-cue).
- `proprioception_delay`: Simulated conduction delay of proprioceptive feedback (bins)
- `vision_delay`: Simulated conduction delay of visual feedback (bins)

**Table S6.** Task-training dataset parameters (RandomTarget): All timings are in bins (10 ms)

| Task parameter | RandomTarget (train) | RandomTarget (sim) |
| --- | --- | --- |
| n_trials | 1100 | 1100 |
| n_timesteps | 155 | 155 |
| action_noise | 0.005 | 0.005 |
| proprioception_noise | 0.005 | 0.005 |
| vision_noise | 0.005 | 0.005 |
| context_input_noise | 0.04 | 0.04 |
| proprioception_delay | 2 | 2 |
| vision_delay | 5 | 5 |
| % catch trials | 20 | 0 |
| % bump trials | 50 | 50 |
| target_on | [10,30] | 30 |
| go_cue | [targ_on +5, trial_end] | 70 |
| bump_mag | [5,10] | [5,10] |

##### Trial Structure

Each trial began with the effector endpoint at a random location in the workspace (Fig. 3G), and was composed of 3 phases 1) Context: the model held at the start location. 2) Delay: A target position (x/y) was presented to two of the context inputs of the task-trained model. 3) Reach: The target information was removed and a step-function go-cue was presented, cueing the model to reach to the target location and hold there until the end of the trial. Because information about the target location was not present after the go-cue, the TT model was obligated to remember the target location while making the reach. Additionally, we applied external bump perturbations to the hand at random times in the trial, which were randomly directed and of a variable magnitude. This resulted in reaches with and without corrective movements, thereby increasing the diversity and nature of computation required to complete the task successfully.

##### Loss Function

We trained TT models to minimize the mean squared error between the effector endpoint and the desired endpoint (i.e. the start location prior to the go-cue, and the target location after the go cue). To encourage strategies with minimal energy expenditure, the TT model was also trained to minimize an *L*_2_ norm of the muscle activations. To improve training stability, we incrementally increased the window of loss included in the training. After 200 epochs, the entire trial contributed to the training loss equally.

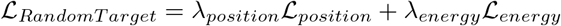

where:

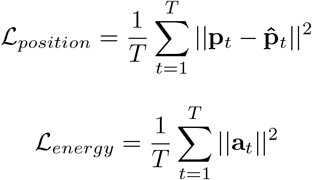

where **p**_*t*_ is the effector endpoint position, 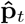 is the desired endpoint position (start location before the go-cue, target location after), and *λ*_*position*_ and *λ*_*energy*_ are the weights on the position and energy terms, both set to 1.0 for the canonical task-trained model.

##### Task-Trained Model (NoisyGRU)

For the canonical RandomTarget dataset, we used a NoisyGRU model (described in Section B.1.1) with the hyperparameters shown in Table S7. As in Section B.1.2, we also penalized the *L*_2_ norm of the hidden activity of this model to force the activity to live closer to the origin, which produces more interesting dynamical structures.

**Table S7.** NoisyGRU model parameters (RandomTarget)

| Hyperparameters | MultiTask |
| --- | --- |
| learning_rate | 5e-3 |
| max_epochs | 2000 |
| input_size | 17 |
| output_size | 6 |
| latent_size | 128 |
| intrinsic noise | 0.01 |
| ic_noise | 0.05 |
| hidden_L2_penalty | 1e-6 |

#### B.1.4 PhaseCodedMemory

##### General Description

A second simple task included in CtDToolkit is the PhaseCodedMemory task, based on recent work investigating short-term memory in the hippocampus [62]. The task receives inputs **u**_*t*_ ∈ ℝ^3^ consisting of a reference oscillation and two stimulus channels, and produces a scalar output *x*_*t*_ ∈ ℝ representing the phase-shifted oscillation encoding stimulus identity. Our implementation can be found in `ctd/task_modeling/task_env/phase_coded_memory.py` in the PhaseCodedMemory class. In Supp. Figure S4 we show the structure of inputs and outputs of the model in more detail:

##### Environment Parameters

The PhaseCodedMemory class takes the parameters described below. Values used in the canonical PhaseCodedMemory dataset are given in Table S8.

- `n_timesteps`: The length of each trial in bins.
- `noise`: Standard deviation of Gaussian noise to be added to the inputs. This noise is pre-computed and the same every time a given trial is presented.
- `post_stim_l2_weight`: Scale of L2 penalty applied to the latent activity (required for model to converge to phase-coding strategy).
- `lat_l2_full`: Boolean. Whether to apply the latent L2 to the full trial, or just the post-stimulus period.
- `pred_loss_full`: Boolean. Whether to apply the prediction loss to the full trial or just to the post-stimulus period.

##### Trial Structure

Trials begin with the reference oscillation with a initial phase drawn from a uniform random distribution from [0, 2*π*]. This oscillation is provided to the TT model on input channel 1. After a delay (drawn from a uniform random distribution from [60, 120] bins from the start of the trial), a stimulus appears on either input channel 2 or 3 (Stim A, B, respectively). This stimulus is maintained for a duration drawn from a uniform random distrubtion ranging from [60, 80] bins. We show example trials of PhaseCodedMemory in Figure S4B.

**Table S8.** Parameters for the task-training and simulation datasets for PhaseCodedMemory.

| Task parameter | PhaseCodedMemory (train) | PhaseCodedMemory (sim) |
| --- | --- | --- |
| n_trials | 5000 | 5000 |
| n_timesteps | 400 | 400 |
| noise | 0.2 | 0.2 |
| post_stim_l2_weight | 0.2 | NA |
| lat_l2_full | False | NA |
| pred_loss_full | False | NA |

##### Loss Function

During training, we penalize the mean squared error between the model output and the desired output in the period after the stimulus offset. Additionally, as previous research found that full-rank RNN models require regularization pressure to converge to a phase-coding solution, we applied an L2 penalty to the latent activity in the post stimulus offset window (see post_stim_l2_weight). See `/ctd/task_modeling/task_env/loss_func.py` for full implementation details.

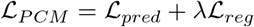

where:

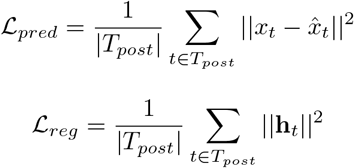

and *T*_*post*_ represents the post-stimulus period, **h**_*t*_ is the model’s latent activity, and *λ* = `post_stim_l2_weight`.

**Table S9.** NoisyGRU model parameters (PhaseCodedMemory)

| Hyperparameters | PhaseCodedMemory |
| --- | --- |
| learning_rate | 1e-3 |
| max_epochs | 50 |
| input_size | 3 |
| output_size | 1 |
| latent_size | 512 |
| latent_ic_var | 0.05 |
| intrinsic noise | 0.01 |

##### Task-Trained Model (NoisyGRU)

For the canonical PhaseCodedMemory dataset, we used the same class of Noisy-GRU as for the 3BFF task. The noise at each time step was drawn from a standard normal distribution with mean zero and isotropic variance of 0.01, while the initial conditions were drawn from a standard normal with mean *µ* and variance of 0.05, where *µ* was a trainable parameter. This ensured that the model learned to stably encode memory regardless of semi-random initial conditions. Full hyperparameters are in Table S9. We provide the NoisyGRU model, fully-trained on PhaseCodedMemory task, in the `content/trained_models/task-trained/tt_PhaseCodedMemory/` folder of the CtDToolkit repository.

#### B.1.5 ChaoticDelayedMatching

##### General Description

ChaoticDelayedMatching extends CtDToolkit to the near-chaotic dynamical regime while retaining a well-specified computation. The task is a delayed non-match-to-sample paradigm based on [20]. The task receives inputs **u**_*t*_ ∈ ℝ^2^ consisting of two cue channels (Cue A and Cue B) and produces a scalar output *x*_*t*_ ∈ ℝ that reports, during a response window, whether the two cues presented on a given trial matched (A-A or B-B, target − 1) or differed (A-B or B-A, target +1). Our implementation can be found in `/ctd/task_modeling/task_env/chaotic_delayed_matching.py` in the ChaoticDelayedMatching class. In Supp. Figure S5 we show the task structure and the chaotic signatures of the task-trained dynamics.

**Figure S5.**
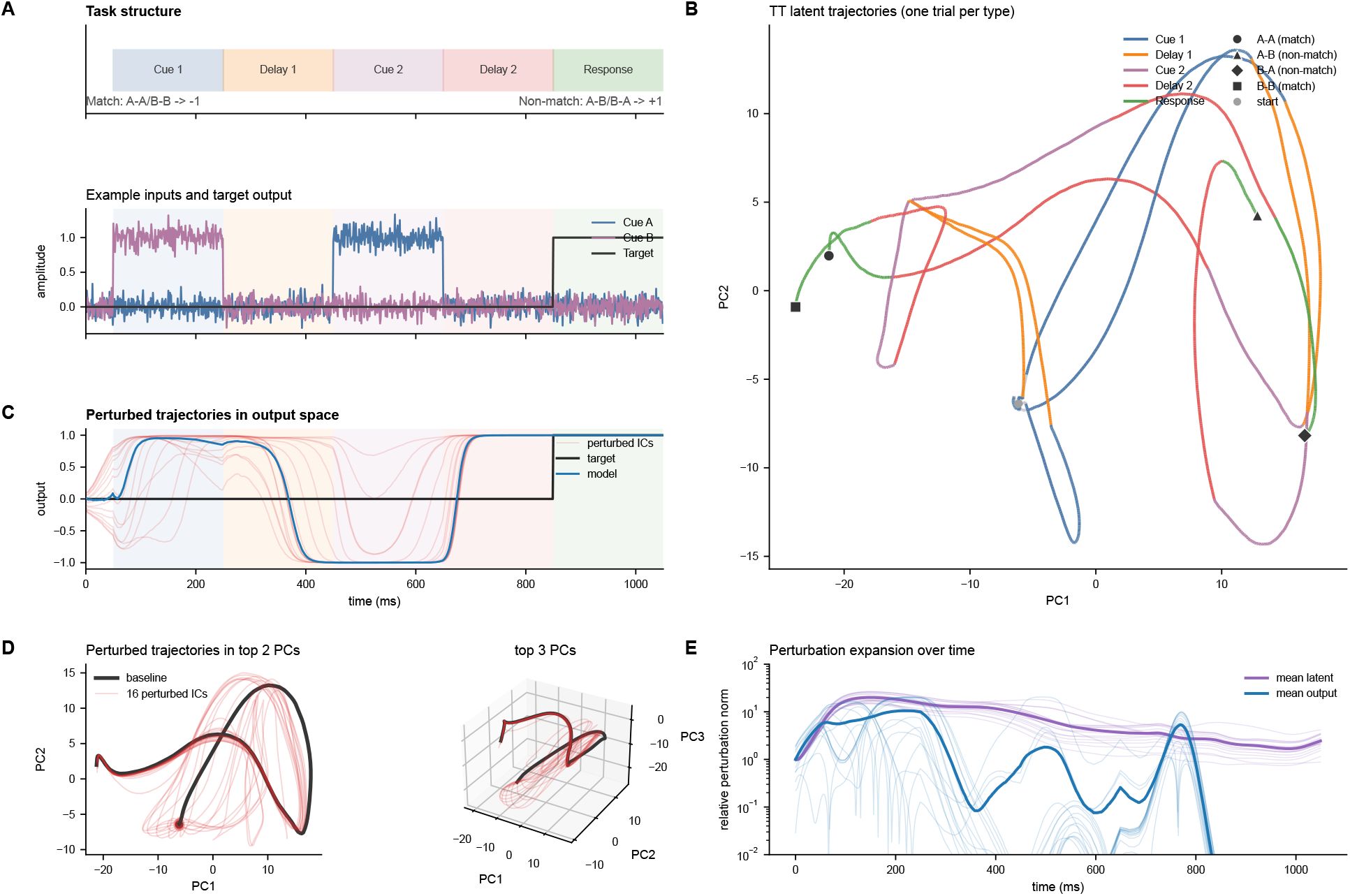
ChaoticDelayedMatching Task-Trained Model and Chaotic Signatures. **A**) Task structure (top row): each trial comprises a baseline period, a first cue (A or B), a delay, a second cue (A or B), a second delay, and a response period; the task-trained model reports match (A-A/B-B → − 1) versus non-match (A-B/B-A → +1). Bottom row: the two cue inputs and the target output for an example trial. **B**) One example trial of each of the four trial types (A-A, B-B, A-B, B-A), shown as the top two PCs of the task-trained latent activity. Each trajectory is colored by trial phase (legend), and its end-of-trial marker encodes the trial type. **C**) Task-trained model output on an example trial and the effect of small initial-condition (IC) perturbations: the baseline output (blue) tracks the target (black) during the response window, and outputs from an ensemble of rollouts initialized with small random IC perturbations (light red) nonetheless converge to the correct response. **D**) Effect of the IC perturbations on the latent activity, shown in the top two PCs (left) and top three PCs (right): the baseline trajectory (black) and the perturbed ensemble (red) diverge over the course of the trial. **E**) Trial-averaged growth of the perturbation norm in the full latent space (purple) and the output space (blue), on a log scale. Each space’s curves are normalized by their initial (t=0) mean norm so both start at 1, placing the differently scaled latent and output norms on a common relative axis so that the log-slope reflects growth rate rather than absolute size. The latent perturbation expands (sensitivity to initial conditions, a hallmark of chaos) while the output perturbation contracts—reflecting stable computation embedded within near-chaotic dynamics.

##### Environment Parameters

The ChaoticDelayedMatching environment takes the following parameters (canonical values in parentheses):

- `n_timesteps`: total trial length in bins (1050).
- `noise`: standard deviation of the Gaussian noise added to the two cue input channels (0.1).
- `cue_scale`: amplitude of the cue pulses (1.0).
- `baseline_range`, `cue1_range`, `delay1_range`, `cue2_range`, `delay2_range`, `response_range`: the [min, max] duration (in bins) of each trial epoch, drawn per trial; for the canonical dataset these are fixed at 50, 200, 200, 200, 200, and 200 bins, respectively.

##### Trial Structure

Each trial lasts 1050 bins (1 ms per bin) and is divided into six contiguous epochs: a baseline period (50 bins), the first cue (200 bins), a first delay (200 bins), the second cue (200 bins), a second delay (200 bins), and a response period (200 bins). On each cue epoch, a pulse is presented on either the Cue A or Cue B channel. Gaussian input noise (standard deviation 0.1) is added to the cue channels. We show example trials of ChaoticDelayedMatching in Supp. Figure S5A.

##### Loss Function

During training, we penalize the mean absolute error between the model output and the target value, computed on the trial-averaged output within the response window. See `/ctd/task_modeling/task_env/loss_func.py` (ChaoticDelayedMatchingLoss) for full implementation details.

##### Task-Trained Model (ChaoticRateRNN)

In contrast with the NoisyGRU used for the other canonical datasets, ChaoticDelayedMatching is solved by a leaky-integration tanh rate RNN (ChaoticRateRNN) whose recurrent weights are initialized with a gain *g* = 1.5, placing the autonomous network in a near-chaotic regime [8]. The model has a latent size of 200, a leak factor *α* = 1*/*30, no bias terms, and fixed (non-trained) random input weights; only the recurrent weights and the linear readout are trained. We simulate neural activity from 96 units (64 held-in, 32 held-out) using an exponential rectifying nonlinearity and pseudo-Poisson observation noise. We provide the task-trained model in the `/pretrained/20260320_ChaoticDelayedMatching_Final/` folder of the CtDToolkit repository.

## C Neural Data Simulation

Here we provide additional details on the process and implementation of the neural data simulation pipeline and provide a schematic in Figure S6.

**Figure S6.**
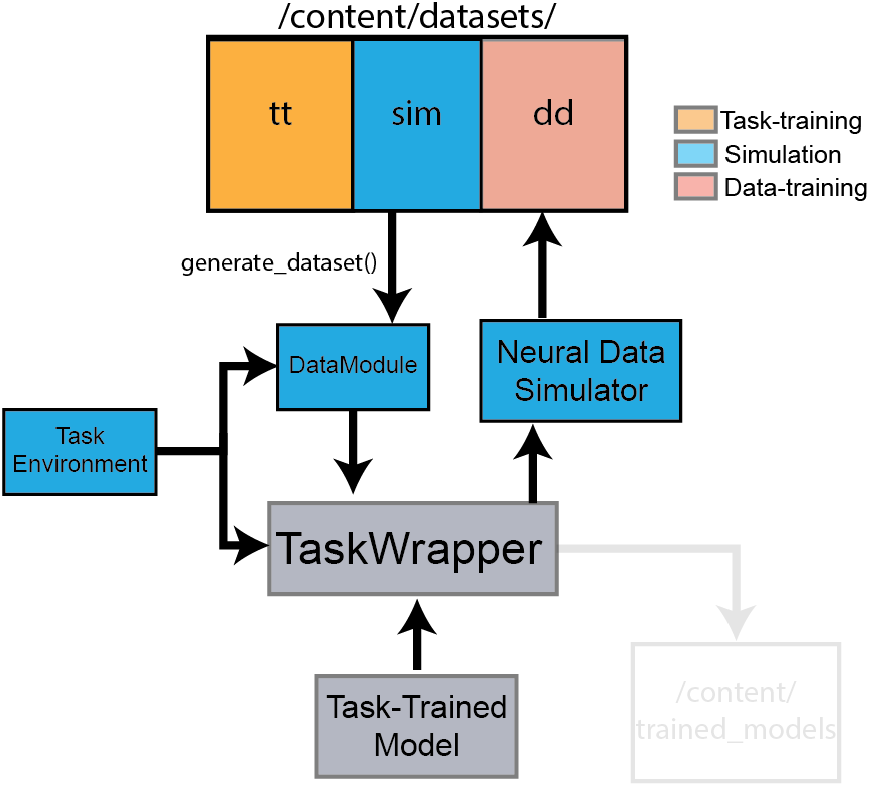
Neural Data Simulation Pipeline. We use a separate directory in the case of task data used for generating the simulated neural data (`sim`) compared to running task training (`tt`). This is because there are often differences in how the task is run to ensure stable and efficient training that are not necessary or desirable to consider as part of the trial structure in the simulated data (see Section B.1.3 and Table S6 for intuition in the case of the RandomTarget task.) Similar to the task-training pipeline, the task data is loaded into a `DataModule`, this time not for training, but instead as a means of curating the trials for passing them through the trained model to obtain the hidden unit activations. Also parallel to the pipeline in Figure S1, the `TaskEnvironment` is loaded in the case of coupled environments to enable feedback at each step. Once the `DataModule` and `TaskEnvironment` are prepared, they are passed to the `TaskWrapper` which coordinates the evaluation of the task-trained model on each of the trials of the task data from `sim`. Once all trials have been processed by the task-trained model, the hidden unit activations are used by the Neural Data Simulator to produce the simulated neural spiking activity for the task (more details in Section C.1). The simulated data is then saved to `dd`, where it can be accessed later when running the data training pipeline (see Section D and Figure S7.)

### C.1 Simulator

#### Sampling simulated neurons

Synthetic neural data used in CtDToolkit is simulated from hidden unit activity of a task trained model. Initially, the requested number of simulated neurons is assigned to hidden units of the task-trained model via a one-hot readout matrix. When the requested neuron count *N* does not exceed the latent dimensionality *D*, units are sampled without replacement (one neuron per latent dimension). When *N > D*, the readout matrix is built by stacking ⌈*N/D*⌉ independent random permutations of the latent indices and retaining the first *N* columns, so that some latent dimensions are observed by more than one simulated neuron. Because each simulated neuron is then passed through the observation nonlinearity and sampled independently (e.g. as an independent Poisson draw), these shared latent dimensions yield distinct noisy “views” of the same underlying rate rather than identical duplicate neurons. This is why, for example, the PhaseCodedMemory dataset can supply 500 + 100 = 600 simulated neurons (Table S10) from a task-trained model with *D* = 512 latent units: with ⌈600*/*512⌉ = 2 stacked permutations, the first 600 of 1024 indices are retained, so 88 latent dimensions are observed by two neurons each.

#### Firing-rate normalization

The activity of these sampled units is then standardized using one of two procedures. By default, we compute a *single* global mean and a *single* global standard deviation across all trials, time points, and sampled units, and transform the activity by subtracting the global mean and dividing by the product of `fr_scaling` and the global standard deviation. Because every unit is normalized by the same two scalars (rather than by unit-specific statistics), this transformation does not alter the relative magnitudes of activity across units inherited from the latent states of the TT model. Alternatively, when `target_mean_rates` is specified, the global normalization is skipped; instead, per-unit additive biases are fit so that, after the rectifying nonlinearity, each unit attains a specified target mean firing rate (this pathway is used here only for the PhaseCodedMemory dataset, for which we targeted low average firing rates).

#### Spike generation

The activity is then rectified and used to define a stochastic process from which spiking events are sampled over the course of the trial. For all of our experiments, the simulated spiking data was sampled from a Poisson process, but for flexibility we include options for sampling from distributions that are under-dispersed or over-dispersed compared to Poisson, such as the Binomial and Negative Binomial distributions, respectively.

#### Held-out neurons

The co-smoothing bits-per-spike metric requires a set of held-out neurons in addition to a held-in set. The number of held-out neurons varies by dataset—for example, 10 for 3BFF, MultiTask, and RandomTarget, and 100 for PhaseCodedMemory (Table S10)—each sampled from the TT network in addition to the held-in set for computing co-BPS.

The neural data simulator is called after the TT model is trained, as part of the Task Training Pipeline (more information in Sections B and F). The Simulator has parameters that control the number of artificial neurons to sample, firing rate scaling, and how hidden activity is embedded into neural rates. Parameters for the neural data simulator across tasks can be seen in Table S10.

The general API for the Simulator is as follows:

#### Variables

- `neuron_dict`: dictionary containing arguments about how many neurons should be in the held-in/held-out datasets.
  – `n_neurons_heldin`: Number of held-in neurons to feed into DD models
  – `n_neurons_heldout`: Number of held-out neurons to be predicted by the DD model (in addition to held-in).

- `embed_dict` : dictionary containing arguments related to how the hidden unit activity of the task-trained model is processed into firing rates. Example keys include:
  – `rect_func` : The type of function to use for rectifying the hidden unit activity (e.g. The rate parameter of the Poisson distribution must be greater than zero.)
  – `fr_scaling` : Scaling factor applied to firing rates. Higher values of `fr_scaling` lead to lower firing rates.
  – `target_mean_rates` : (optional) Alternative to fr_scaling, in which firing rates are optimized such that the mean firing rate of the neurons match a desired target.

- `noise_dict` : dictionary containing arguments related to how spiking events are sampled from the simulated firing rates. Example keys include:
  – `obs_noise` : *String*. For all datasets included in CtDToolkit, we used a generalization of the Poisson distribution that considers the dispersion of the probability density function, controlled by a dispersion factor given by `dispersion`.
  – `dispersion` : Parameter controlling the level of dispersion of the probability density function. For `dispersion` = 1.0, the distribution is Poisson.

**Table S10.** Canonical dataset simulation parameters.

| Simulation Parameters | 3BFF | PhaseCodedMemory | MultiTask | RandomTarget |
| --- | --- | --- | --- | --- |
| n_neurons_heldin | 50 | 500 | 50 | 50 |
| n_neurons_heldout | 10 | 100 | 10 | 10 |
| rect_func | exp | exp | exp | exp |
| fr_scaling | 2.0 | NA | 2.0 | 4.0 |
| target_mean_rates | NA | 2.0 | NA | NA |
| obs_noise | Poisson | Poisson | Poisson | Poisson |
| dispersion | 1.0 | 1.0 | 1.0 | 1.0 |

#### Methods

`generate_simulated_data()`: Uses task-trained model to generate simulated neural responses, as well as collect additional information about the task and trials from the datamodule used to train the TT model.

#### Arguments

- `task_trained_model` : Task Wrapper (described in Section F) object for the TT model after it has been fit to the task data from `datamodule`.
- `datamodule` : PyTorch Lightning Data Module for serving the task input/output data to the TT model.
- `seed` : Makes data generation deterministic for reproducibility (e.g. sampling which TT model hidden units to turn into neurons, sampling of spiking events, etc.)

`simulate_neural_data()`: Calls `generate_simulated_data()`, compiles all fields relevant for data-driven modeling, and saves them as HDF5 files.

#### Arguments

- `task_trained_model` : Task Wrapper (described in Section F) object for the TT model after it has been fit to the task data from `datamodule`.
- `datamodule` : PyTorch Lightning datamodule for serving the task input/output data to the TT model.
- `run_tag` : filename for saving the simulated data to the project, written in `/examples/run_task_training.py`
- `subfolder` : optional subfolder to save data somewhere more specific inside the project, specified by overrides passed to the `train()` function in `ctd/task_modeling/task_train_prep.py`
- `dataset_path` : highest level save path, acquired from `path_dict` in `examples/run_task_training.py`

Note that the majority of the above arguments are set at a higher level in the task training scripts and passed through to the simulator. Any direct interactions with the simulator are via configuration files.

## D Utilizing the Data-Training Pipeline

### D.1 Training Pipeline Overview

The data training pipeline allows users to deploy models of neural dynamics on the simulated neural data produced by task-trained models. Our design allows for modular training and saving of data-driven model checkpoints and performance metrics, and supports end-to-end model training using PyTorch Lightning.

We use hydra [32] to instantiate configuration files for default model parameters, and Ray-tune [24] for hyperparameter tuning. We recommend using the template provided in `ctd/data_modeling/` for adding new data-driven models or modifying existing models. See Section D.2 for details and see Figure S7 for a schematic.

**Figure S7.**
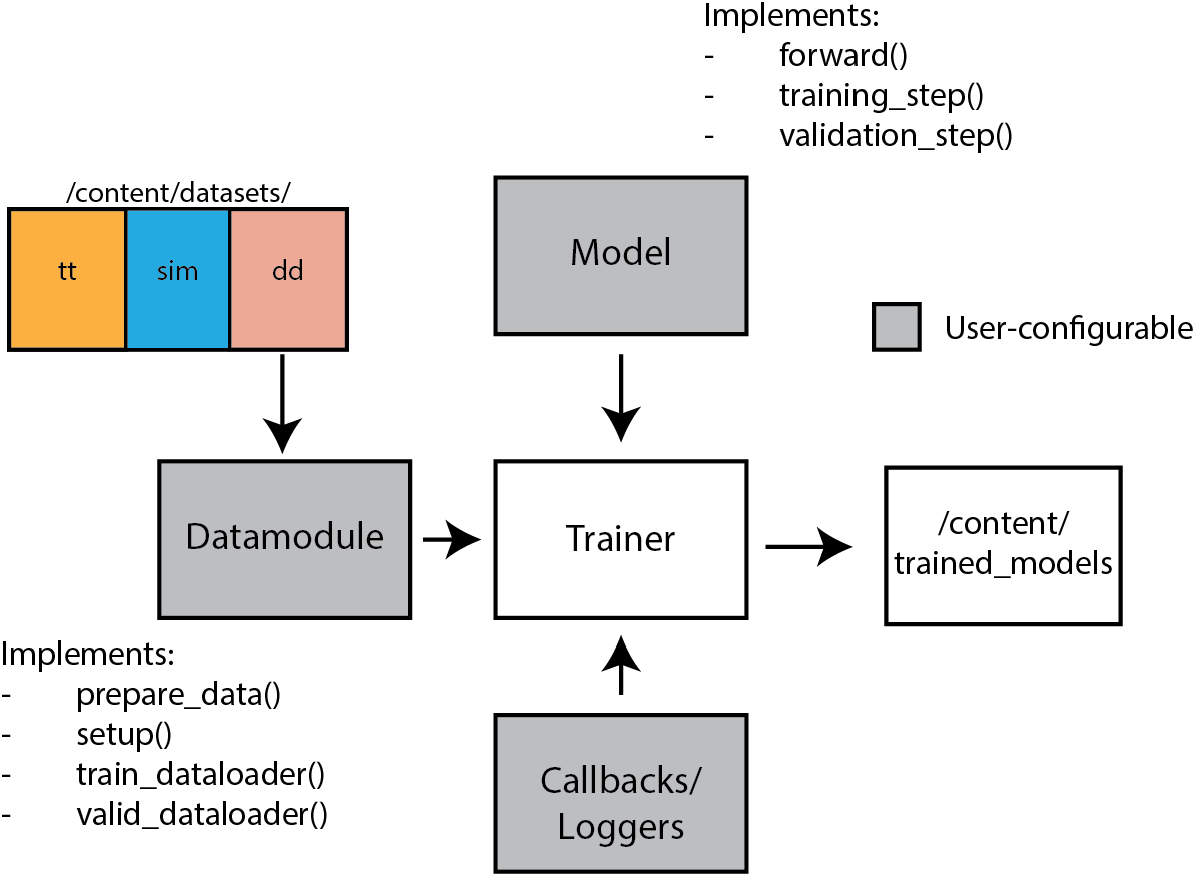
High-level schematic of data-training pipeline. The simulated neural data file is loaded from `dd` by the `prepare_data()` and `setup()` methods of the `DataModule`, and packaged into training and validation dataloaders. These dataloaders are used to train the data-driven model, an implementation of a LightningModule. The Trainer also accepts user-defined callbacks and loggers. The Trainer object automates handling of the training and evaluation loops, as well as model optimization. After training is complete, the data-driven model and the corresponding datamodule are saved to `/content/trained_models`.

A basic DD user can begin training data-driven models on the canonical datasets using the script `/examples/run_data_training.py`. At the top of this script, users must select from the following choices:

1. Choose the data-driven model class (SAE or LFADS).
2. Choose a model architecture (from `/ctd/data_modeling/configs/{MODEL_CLASS}/{MODEL}`)
3. Choose a canonical dataset (options are “NBFF”, “PhaseCodedMemory”, “MultiTask”, or “RandomTarget”).
4. Select whether to infer inputs or supply them to the model (`INFER_INPUTS` flag).

For hyperparameter sweeping, users can modify variables in `SEARCH_SPACE` dictionary to perform hyperparameter sweeps (e.g., `tune.grid_search`) on model parameters. See [24] for additional choices for hyperparameter sweeping.

### D.2 Custom DD Model Interface

To make the addition of user-defined models more straightforward and streamlined, we provide a template class called `TemplateSAE` which can be found at `ctd/data_modeling/models/SAE/template.py`. This template inherits from the Lightning Module object in PyTorch Lightning, and has the following methods and required named outputs:

#### Required Methods

- `forward(data,inputs)` : Defines the series of model operations that act on the data in an end-to-end pass through of the model. Outputs `log-rates` and `latents`.
- `configure_optimizers()` : Handles instantiation of optimizer. This includes determining how different subsets of parameters are trained, as defined by dictionary arguments passed to the user’s optimizer of choice. Must return a PyTorch `Optimizer` object.
- `training_step(batch, batch_idx)` : Definition of the entire end-to-end pass-through of the model, including the computation of the loss and logging of any intermediate metrics at each training step. Receives batches of data from the training dataloader defined in the `datamodule`. Outputs loss on training set for the current step.
- `validation_step(batch, batch_idx)` : Definition of how the model is evaluated on a validation set. This method often parallels `training_step()`, except no gradients are computed. Receives batches of data from the training dataloader defined in the `datamodule`. Outputs loss on validation set for the current step.

This class parallels the SAE baseline, defining all required methods and named outputs in a manner consistent with the rest of the toolkit. Users interested in defining their own models can fill out this template, including any additional custom methods as needed. Provided the user adds the corresponding config files to their repository (mentioned above in Section D.1), they should be able to directly train and analyze their models using the CtDToolkit infrastructure!

### D.3 Baseline Models

As part of CtDToolkit, we provide implementations and/or comparisons for several baselines from across the literature. In the following subsections, we give brief descriptions of each approach, how hyperparameters were selected, and further motivation as to why each method was considered for CtDToolkit.

#### D.3.1 Model Class: Generic Sequential Autoencoder (SAE)

##### Gated Recurrent Unit (GRU)

Our implementation of a GRU SAE can be found at `ctd/data_modeling/models/SAE/dyn_models_GRU.py`. It consists of the following components:

- **Encoder:** A bidirectional GRU processes input sequences to produce hidden states, capturing information from both past and future sequences.
- **Dropout Layers:** Applied to hidden states and initial conditions to prevent overfitting.
- **Initial Condition Linear Layer:** Maps concatenated hidden states from encoder GRU to initial condition for the decoder GRU.
- **Decoder:** An RNN with a GRU cell processes initial conditions and input sequences to produce latent states for each trial.
- **Readout Layer:** A linear map from the latent state **z**_*t*_ to per-neuron log-firing-rates, log 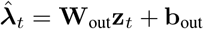. Firing rates are then obtained by exponentiating, 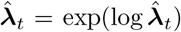, which guarantees non-negativity (the canonical linear–exponential observation model).

The forward pass involves:

1. Passing the data through the encoder to get hidden states.
2. Applying dropout and linear mapping hidden states to the latent space.
3. Using the decoder to produce latent states from initial conditions and supplied inputs.
4. Mapping latent states to log-rates via the linear readout layer, then exponentiating to obtain non-negative firing rates.

The model uses a Poisson negative log-likelihood loss function to compute the difference between these predicted rates and the target spikes, with the potential for alternative loss contributions such as weight decay, etc. The Adam optimizer is used for training, with learning rate adjustments and weight decay to regularize the model.

##### Linear Dynamical System (LDS)

We used a simplified implementation of an LDS to illustrate the utility of the metrics included in CtDToolkit. For this, we replaced the GRU cell from Section D.3.1 with a cell that could only model simple linear dynamics, according the equations:

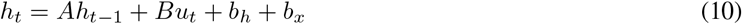

Where:

- *h*_*t*_ is the hidden state at time step *t*.
- *h*_*t*−1_ is the hidden state at the previous time step.
- *u*_*t*_ is the input at time step *t*.
- *A* is a trainable weight matrix for the hidden state update.
- *B* is a trainable weight matrix for the input affecting the hidden state.
- *b*_*h*_ is a trainable bias vector for the hidden state update.
- *b*_*x*_ is a trainable bias vector for the input affecting the hidden state.

##### Neural ODE (NODE)

Our implementation of a SAE class based on Neural Ordinary Differential Equations (Neural ODEs [22]) can be found at `ctd/data_modeling/models/SAE/dyn_models.py`. It consists of the following components:

- **Encoder:** A bidirectional GRU processes input sequences to produce hidden states, capturing information from both past and future sequences.
- **Dropout Layers:** Applied to hidden states and initial conditions to prevent overfitting.
- **Initial Condition Linear Layer:** Maps concatenated hidden states from encoder to initial condition for the decoder.
- **Decoder:** An RNN with a Neural ODE cell processes initial conditions and input sequences to produce latent states. We describe the NODEcell in more detail below.
- **Readout Layer:** Linear mapping from latent states to output data dimensions.

The equations that define the state update for the NODE cell are:

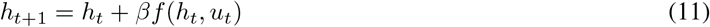

Where:

- *h*_*t*_ is the hidden state at time step *t*.
- *u*_*t*_ is the input at time step *t*.
- *f* is a function representing the change in hidden state, modeled by a Multilayer Perceptron (MLP) with a total of `vf_num_layers` layers that have `vf_hidden_size` hidden units each, and uses ReLU activations between layers.
- *β* is a scalar that improves training stability (set to 0.1).

Unlike typical Neural ODE models, we did not backpropagate the gradients through an ODE solver; instead, we did single-step Euler updates and used standard backpropagation-through-time to train the model. This approach has been shown to produce similar results on synthetic neural datasets as the ODE-solver-based training [66, 50]. Both forward pass and training procedure are the same as in the GRU model (Section D.3.1).

Hyperparameters for the NODE-based SAEs shown in Figure 5C can be found in Table S11.

**Table S11.** Hyperparameters for data-driven models (NODE-D) from Figure 5.

| Hyperparameter | NODE |
| --- | --- |
| encoder_size | 100 |
| latent_size | D |
| weight_decay | 0 |
| learning_rate | 2e-3 |
| vf_num_layers | 6 |
| vf_hidden_size | 128 |
| dropout | 0.05 |
| epochs | 1000 |

#### D.3.2 Model Class: Latent Factor Analysis via Dynamical Systems (LFADS)

Latent Factor Analysis via Dynamical Systems (LFADS) [25] is a popular model that uses a latent dynamics model to reconstruct neural activity. LFADS is a sequential autoencoder that uses variational training to learn the dynamics that generate a set of spiking observations. LFADS uses the same reconstruction objective as the SAE models in Section D.3.1, except LFADS also penalizes the KL divergence of the initial condition (IC) encoder against a Gaussian prior. For the CtD Toolkit we used an open source implementation of LFADS [58] for its modularity and extensibility, which we consider desirable to have compatible with CtDToolkit and available for users as part of the toolkit. Full implementation details are given in [58].

For variants of the SAE that are used to infer inputs, there is an additional bidirectional GRU encoder (referred to as the controller-input encoder, or CI Encoder) that outputs a sequence that is passed to another GRU referred to as the “controller”. The controller is responsible for producing time-varying inputs to the generator based on feedback from the generator at the preceding timestep, and the output of the CI encoder. The weights of the controller and CI encoder are updated to optimize the same loss as the rest of the components of LFADS, while also being directly penalized by the KL divergence against a user-chosen prior distribution. In the experiments shown in Figure 6, we use a Student’s-*t* distribution (*df* = 5) to encourage the controller to infer sparse inputs. Alternative choices supported in CtDToolkit include an Autoregressive Gaussian as well as standard Gaussian priors. Custom priors can be easily added by users. In total, the loss for training an LFADS model in CtDToolkit is given by ℒ in Equation 12 as follows:

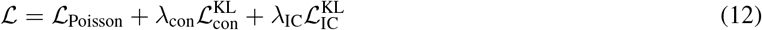

Here, ℒ_Poisson_ is the Poisson negative log-likelihood reconstruction loss, 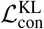 and 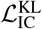 are the KL-divergence penalties on the controller and initial-condition (IC) encoders against their priors, and *λ*_con_ and *λ*_IC_ are their respective weights, with *λ*_IC_ = 0 when inputs were supplied. We considered the LFADS model class as a baseline due to its customizability, enabling researchers to have an accessible but flexible model available for their initial experiments and to use for gaining intuition about Computation through Dynamics.

We provide hyperparameters for the LFADS models included in Figure 6 in Table S12. The “KL Penalty” in Section 3.3 and Figure 6 refers to the parameter `kl_co_scale`, where we tested how varying this parameter affected the quality of inferred dynamics.

We also include an LFADS variant that uses a NODE generator in place of the standard GRU RNN. NODE-based SAEs have been found to improve the ability of dynamics models to infer accurate latent activity [40, 50]. This model provides users with an experimental model with which they can begin probing how the generator might affect performance on the included CtDToolkit metrics. Switching from GRU to NODE-based LFADS is controlled by the gen_type parameter, which takes in a string (default: “RNN”). Changing this parameter to “NODE” changes the generator class to a Neural ODE. We provide a default configuration file (ResLFADS – for residual LFADS) so that users can easily test the effects of a NODE-based generator on the canonical datasets.

##### Low firing-rate cold start (PhaseCodedMemory)

By default, the LFADS rate readout uses a fan-in linear layer initialized with zero bias and Gaussian weights of standard deviation 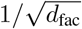. Under the Poisson exp-link, this produces an initial predicted rate of ~ 1 spike/bin per neuron with a log-rate standard deviation of ~ 1 nat. For high firing-rate datasets (e.g. 3BFF, MultiTask, RandomTarget, ChaoticDelayedMatching) this initialization is well-conditioned. For low firing-rate datasets (e.g. PhaseCodedMemory, which simulates hippocampal CA1 neurons at ~ 2 Hz; ~ 4 ×10^−3^ spikes/bin) it places the network many natural-log units above the data, and the resulting large exp(·) pre-activations make Poisson NLL gradients numerically unstable during the first epochs of training. We therefore provide two LFADS configuration options for cold-starting the rate readout, used in the PhaseCodedMemory configurations and disabled (default) on all other canonical datasets: (i) `rate_bias_init` (bool), which at the start of fresh training sets the readout’s per-neuron bias to 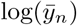 where 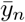 is the empirical mean spike count of neuron *n* on the training set; and (ii) `rate_weight_init_scale` (float, default 1.0), which multiplies the readout weight matrix at initialization. With the bias initialized at 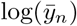 and the weight matrix shrunk by a factor *s*, each neuron’s predicted rate begins at its marginal mean with 𝒪 (*s*) log-rate variability around it, which we found to substantially stabilize training on PhaseCodedMemory. The PCM canonical configurations (`PhaseCodedMemory_LFADS.yaml`, `PhaseCodedMemory_InjLFADS.yaml`) use `rate_bias_init: True` and `rate_weight_init_scale: 0.1`. Both flags are guarded to the Poisson exp link, take effect only at the start of fresh training, and have no effect when loading a saved model for inference or continued training.

**Table S12.** Hyperparameters for data-driven LFADS models in Figure 6.

| Hyperparameter | LFADS |
| --- | --- |
| gen_type | RNN |
| ic_encoder_size | 128 |
| gen_dim | 128 |
| ci_lag | 1 |
| con_dim | 128 |
| inf_inp_dim | 3 |
| fac_dim | 20 |
| cd_rate | 0.3 |
| co_prior.df | 5 |
| co_prior.loc | 0 |
| co_prior.scale | 1.0 |
| ic_prior.mean | 0 |
| ic_prior.variance | 0.1 |
| dropout | 0.02 |
| cell_clip | 5.0 |
| lr_init | 1.0e-3 |
| lr_stop | 1.0e-5 |
| lr_decay | 0.95 |
| lr_adam_beta1 | 0.9 |
| lr_adam_beta2 | 0.999 |
| lr_adam_episolon | 1.0e-7 |
| weight_decay | 1.0e-5 |
| kl_ic_scale | 1.0e-6 |

## E Analysis and Comparison Methods

### E.1 Analysis Objects

#### Task-Trained Analysis

We provide an object that abstracts the data handling away from the task-trained models or datamodules. This class `Analysis_TT` can be found at `ctd/comparison/analysis/tt/tt.py`, and contains helpful functions to obtain latent activity, compute fixed-points, and plot trials. Example usage of the `Analysis_TT` object can be found in `examples/compare_dd_tt_models.ipynb`. In addition, task-specific analysis objects can be found in the subfolder `ctd/comparison/analysis/tt/tasks`. These include functions to compute fixed-points during specific tasks and phases in MultiTask, among other useful features.

#### Data-Driven Analysis

We also provide an analogous object for data-driven models called `Analysis_DD`, located at `ctd/comparison/analysis/dd/dd.py`. By passing the model class (“SAE” or “LFADS”), this model standardizes the functions used to obtain hidden activity, predicted rates, and other metrics from data-driven models of any class included in CtDToolkit. This object also allows for easy fixed-point computation and visualization.

#### Comparison

The final object included in CtDToolkit is the `Comparison` object, located at `ctd/comparison/comparison.py`. Using `load_analysis()`, this object takes in `Analysis` objects (either task-trained or data-driven) and allows automated comparisons of metrics across models. Example usage of the `Comparison` object can be seen in `examples/compare_tt_dd_models.ipynb`.

### E.2 Fixed Point Finding

As discussed in Section 2.4.3, fixed point structure and character can provide an informative qualitative picture of the neural dynamics, and possibly even their goal orientation. We seek to find the points in state space that minimize the magnitude of the kinetic energy of the dynamics, given by Δ*h*^2^ ~ *q* in Equation 13. Because the estimated flow field is essentially never exactly zero to mathematical precision, we instead retain points whose speed falls below a chosen threshold; these slow regions act as *functional fixed points* over the timescales relevant to the behavior, rather than true fixed points where the flow vanishes exactly. The dynamics near such points are sufficiently slow to enable characterization of the dynamics through linearization. The location and stability of each of these fixed points gives us insight into the organization and nature of the features and motifs of the dynamics.

To perform fixed point finding, we use an extension of an existing fixed point finding package [23]. Using ADAM to perform stochastic gradient descent, we optimize the objective given in Equation 13.

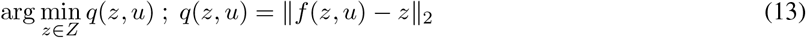

Where *q*(*z, u*) represents the magnitude of the state update at state *z* given inputs *u*. Only fixed points with *q* less than a threshold (depending on specific model) are retained for further analysis.

We characterize fixed point stability by the magnitude and sign of the eigenvalues of the Jacobian of the state transition function, evaluated at each fixed point. In a discrete dynamical system, if all eigenvalues have magnitudes within the unit circle, the fixed point is stable; otherwise, it is unstable.

### E.3 Dynamical Similarity Analysis (DSA)

DSA is a method that combines data-driven techniques inspired from Koopman Operator Theory with Statistical Shape Analysis to identify, from data, whether or not two dynamical systems are equivalent. Two dynamical systems *f* and *g* are equivalent, or topologically conjugate, if there exists a homeomorphism *ϕ* that maps one system to another as follows:

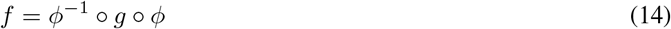

A natural similarity metric thus could be min_*ϕ*_ *f* (*x*) − *ϕ*^−1^ ∘ *g*(*y*) ∘ *ϕ*, averaged over samples from *x* and *y* recorded in a one-to-one fashion. Unfortunately, two problems exist with this direct approach. First, *ϕ* can be arbitrarily nonlinear and hence hard to optimize for, especially with insufficient data. Second, the two systems could be recorded independently, which may make it difficult to align trajectories across systems. DSA circumvents these two problems by mapping each system to an arbitrarily high-dimensional linear system. Conjugacies between linear systems are themselves linear maps, hence the similarity metric becomes Procrustes Analysis over Vector Fields (PAVF):

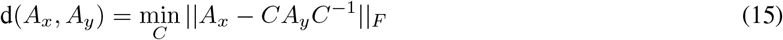

Here, *A*_*x*_ and *A*_*y*_ are the dynamics matrices of the linear systems fit to the high-dimensional embeddings of the two systems being compared, and *C* is a change-of-basis matrix. Typically in DSA, *C* is optimized over the orthogonal group, *O*(*n*), rather than the more general group of invertible matrices *GL*(*n*). This is because the metric over *O*(*n*) is a proper metric, which has yet to be proven for *GL*(*n*), and *O*(*n*) is a compact group. Importantly, [56] showed that this method can be used to identify conjugacy even when *ϕ* was highly nonlinear. In this paper, we used the angular form of the distance, which scales from 0 (most similar) to *π/*2 (most dissimilar). In practice, however the full range is not always used, and so relative distances are typically emphasized over absolute.

According to Koopman Operator Theory [6], dynamical systems can be represented linearly by mapping them into a high-(oftentimes infinite-)dimensional space. Methods that approximate the Koopman operator include Dynamic Mode Decomposition, which is used in DSA [17]. Koopman Operator Theory suggests that operator eigenspectra can characterize conjugacy [12]; accordingly, other work has used the 2-Wasserstein distance between operator eigenvalues to compare dynamical systems [64], but we found that the PAVF better differentiated the models studied here.

#### E.3.1 Picking Hyperparameters

In practice, at least two main hyperparameters are required in delay-embedded DSA, which we chose to use here for its speed. First, the number of delays must be chosen to create the high-dimensional embedding–we swept a large range and did not find a significant change in the results. Second, low-rank regression is used to fit the dynamics matrices, hence the rank must be chosen. Here, we chose to reduce dimensionality of each of our systems first with PCA to the top 5 components, for which previous work suggests choosing a full rank dynamics matrix. However, if intuition is lacking, predictivity metrics can be used to help select hyperparameters.

#### E.3.2 Comparing DSA and fixed-point structure across model capacity

To illustrate how dynamical similarity and fixed-point structure depend on data-driven model capacity, we fit NODE- and GRU-based SAEs to the canonical 3BFF dataset across latent sizes 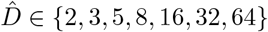 (a single representative seed per size for the matrices and fixed-point panels) and compared each to the ground-truth 3BFF TT model using DSA and fixed-point finding (Supp. Fig. S8). Comparing the boxed DSA-to-TT row of the cross-comparison matrices (Supp. Fig. S8A), replotted against latent size in Supp. Fig. S8B, reveals qualitatively different trends for the two model families: NODE dynamics are most TT-like at intermediate capacity 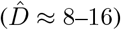 and become *more* dissimilar at high capacity, whereas GRU dissimilarity decreases monotonically and plateaus. This mirrors the fixed-point structure (Supp. Fig. S8C): the 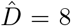 NODE recovers a clean cube of stable and unstable fixed points, while the 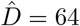 NODE develops a tangled, off-manifold structure, and both GRU models retain the expected cube. The DSA and fixed-point-finding hyperparameters used for this figure are given in Table S13, and the figure is produced by `examples/figures/FigDSA/make_figure_dsa.py`.

**Figure S8.**
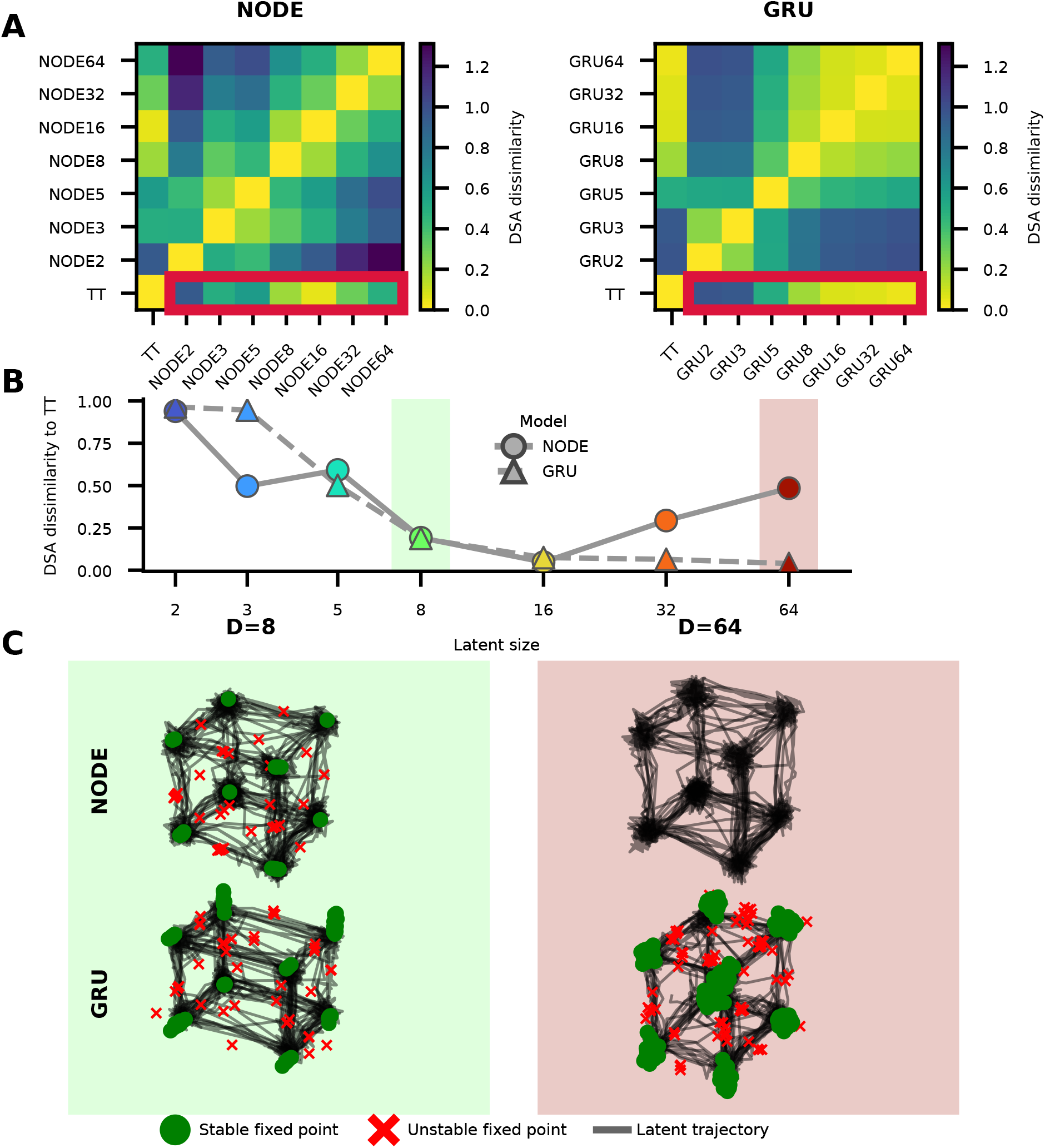
DSA and fixed-point structure for NODE and GRU models across latent size (3BFF). **A)** DSA dissimilarity matrices comparing the ground-truth 3BFF TT model and data-driven NODE (left) and GRU (right) models across latent sizes (rows and columns; the TT model is at the bottom-left, with latent size increasing upward and to the right). Both matrices use a common color scale. The boxed bottom row marks the TT-vs-model dissimilarities — excluding the trivial TT-vs-TT self-comparison — which are replotted in B. **B)** DSA dissimilarity to the TT model as a function of latent size for NODE (circles, solid) and GRU (triangles, dashed). Marker color encodes latent size. The green and red shaded bands mark the latent sizes (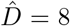 and 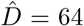) shown in C. **C)** Inferred fixed points (green circles: stable; red × : unstable) over latent trajectories (black) for the 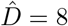 and 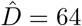 NODE and GRU models. To enable visual comparison across models, each model’s latent trajectories and fixed points are linearly projected onto the ground-truth 3-bit task state, so all panels share a common (bit-1, bit-2, bit-3) orientation; fixed points are computed under zero external input. For visualization, fixed points farther than 0.8 of the trajectory cloud’s RMS radius from any inferred trajectory were discarded, and fixed points within 0.02 of that radius were merged. Panel backgrounds are tinted to match the corresponding bands in B. Hyperparameters in Table S13.

**Table S13.** Hyperparameters for the DSA fits and fixed-point finding in Figure S8.

| Parameter | Value |
| --- | --- |
| *DSA (delay-embedded, angular distance)* |  |
| n_delays | 20 |
| rank | 30 |
| delay_interval | 1 |
| optimization iterations | 1500 |
| learning rate | 5e-3 |
| score method | angular |
| trials used | all validation trials |
| *Fixed-point finding* |  |
| n_inits | 4096 |
| learning rate (ADAM) | 1e-3 |
| max iterations | 10000 |
| noise scale | 0 |
| external input | 0 (zero drive) |
| $q$ threshold | 1e-3 |
| seed | 0 |

### E.4 Nonlinear Cycle Consistency

An alternative nonlinear cycle-consistency metric [66] has been published for datasets with arbitrary nonlinear observation models; below we give its definition and our implementation. Let 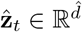 denote the inferred latent variables at time *t*, and let **ŷ**_*t*_ ∈ ℝ^n^ denote the corresponding predicted firing rates produced by the forward inferred observation model 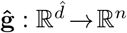 (i.e.,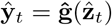). *Notation: noise-perturbed quantities are written with a tilde and a superscript* (*σ*) *(e*.*g*.,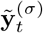*), while baseline inferred quantities retain hats (e*.*g*.,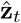).

**1. Learned generalized inverse** We first train a nonlinear mapping 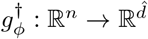 (an MLP parameterized by *ϕ*) to approximate an inverse of the observation model **ĝ** from firing rates to latent variables. The parameters *ϕ* are optimized to minimize MSE on a training set:

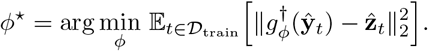

Here 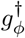 is a *learned* generalized inverse of **ĝ**; it need not coincide with a true inverse and is trained independently from **ĝ**.

**2. Noise injection**. After training, for each validation sample, we inject independent Gaussian noise into the predicted rates, scaled by each neuron’s empirical variability:

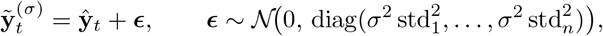

where std_*i*_ is the standard deviation of neuron *i*’s log-rates in the training data. This ensures perturbations are proportional to each neuron’s natural firing-rate scale.

**3. Latent reconstruction**. We apply the trained generalized inverse to the perturbed rates to obtain reconstructed latents:

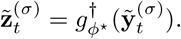

**4. Quantification of degradation**. For each noise level *σ*, we evaluate the coefficient of determination (*R*^2^) between reconstructed and original inferred latents on the validation set:

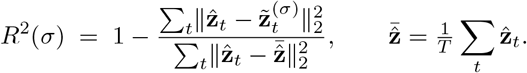

The `nl_cycle_con` curve is *R*^2^(*σ*) as a function of the noise magnitude *σ*. Intuitively, if the forward observation model 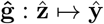 is non-injective, small perturbations in rate space (**ŷ**-space) can yield disproportionately large deviations in reconstructed latents, causing rapid degradation of *R*^2^(*σ*).

We show in Supp. Fig. S9A example `nl_cycle_con` curves for *σ* ∈ [0, 0.01, 0.05, 0.1, 0.25, 0.5], and the relationship between linear and nonlinear cycle consistency in Supp. Fig. S9B (nl_cycle_con evaluated at *σ* = 0.05).

#### MLP hyperparameters and overfitting controls

The learned generalized inverse 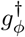 is a multilayer perceptron with two hidden layers of 64 units each and ReLU activations, mapping inferred firing rates to inferred latents. It is trained to minimize the MSE between predicted and inferred latents using the Adam optimizer (learning rate 10^−3^, weight decay 10^−4^) for up to 1000 epochs in full-batch mode. To guard against overfitting of the inverse mapping, the inferred latents and rates are split into training and validation sets and the metric employs early stopping: the validation MSE is monitored each epoch, training halts when the validation loss fails to improve by at least 10^−5^ for 50 consecutive epochs, and the best-validation parameters are then restored. Because a brittle, overfit inverse could itself amplify small rate perturbations and produce rapid degradation of *R*^2^(*σ*) that could be mistaken for a non-injective observation model, we recommend confirming that the training and validation losses track closely (i.e., a small generalization gap) before interpreting the `nl_cycle_con` curve. For the results shown here, the inverse MLP was trained with early stopping on validation loss as described above, and we observed no evidence of overfitting: the training and validation loss trajectories tracked closely across all latent sizes (Supp. Fig. S9C). All of these hyperparameters are exposed in the metric configuration and can be adjusted by the user.

**Figure S9.**
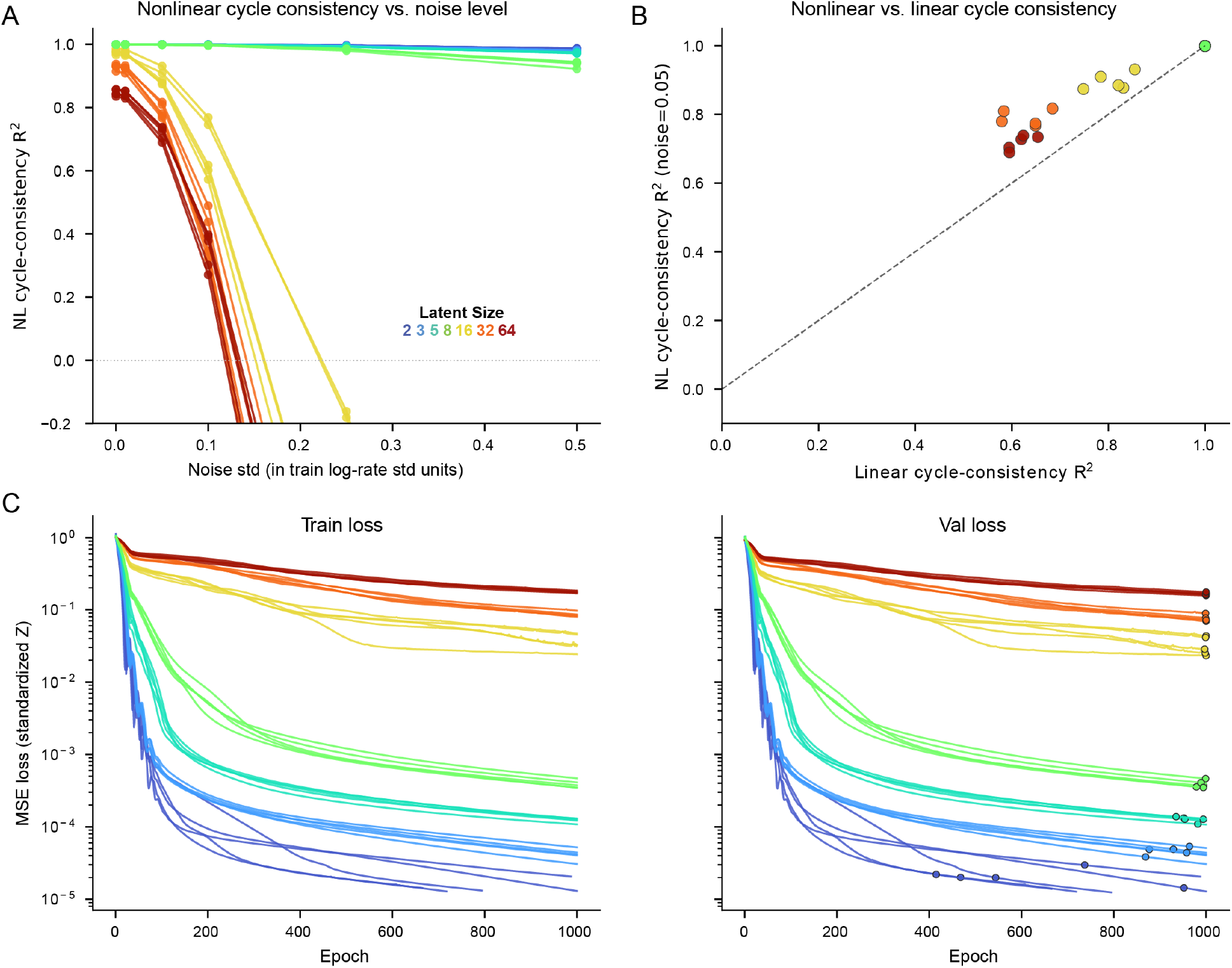
Nonlinear Cycle-Consistency Metric for DD-NODE models trained on NBFF Canonical Dataset: Color indicates DD-NODE latent size across all panels. **A)** `nl_cycle_con` curves (*R*^2^ vs. injected noise standard deviation, in units of the per-dimension training log-rate std) for DD-NODE models across latent sizes and random seeds. Between 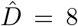 and 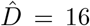, we see a sharp divergence in noise robustness, suggesting the emergence of a non-injective mapping. **B)** Scatter plot comparing nonlinear cycle-consistency (y-axis, evaluated at *σ* = 0.05) against linear cycle-consistency (x-axis). We find a strong linear relationship between these two metrics. **C)** Training diagnostics for the learned inverse MLP 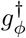 used by the metric: training and validation MSE loss versus epoch, plotted on a shared log axis. Training and validation losses track closely across all latent sizes, indicating that the inverse mapping is not overfit and that the loss of noise robustness in panel A reflects non-injectivity of the observation model rather than a brittle inverse.

### E.5 Distribution Matching (Wasserstein Distance)

The metric takes as input training and validation trajectories from the reference data and the corresponding model-predicted activity. Because raw spikes can produce noisy empirical distributions, trajectories may first be temporally binned and/or smoothed. Optionally, we fit principal components analysis (PCA) using only the training observations and project both the reference activity and model predictions into this same PCA space; when PCA is disabled, the comparison is carried out directly in the original observation channels. When used, this projection ensures that the comparison is performed in a shared observation-derived coordinate system rather than in the potentially arbitrary latent coordinates of the model.

Optionally, we construct a delay embedding of the PCA-projected trajectories. For a trajectory **x**_1:*T*_, with delay lag *τ* and number of delays *K*, the delay-embedded state at time *t* is given by

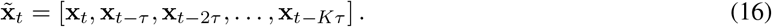

This embedding augments each sample with recent temporal history, allowing the metric to compare distributions over local trajectory segments rather than instantaneous activity alone. After optional PCA projection and delay embedding, the validation trajectories are flattened across trials and time to produce two point clouds: 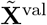 for the reference activity and 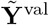 for the model-predicted activity. For computational efficiency, each point cloud may be randomly subsampled to a fixed maximum number of points before the distributional comparison.

Following the marginal CDF-based approach, we compute a one-dimensional Wasserstein-1 distance independently for each dimension of the flattened point clouds. For dimension *d*, this distance is

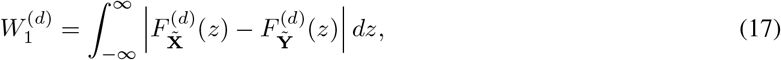

where 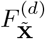 and 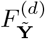 are the empirical cumulative distribution functions of the reference and model-predicted samples along dimension *d*. Equivalently, in one dimension this distance can be computed by comparing the empirical quantile functions of the two distributions.

To make the metric comparable across dimensions with different scales, we normalize each dimension by the range of the reference validation samples:

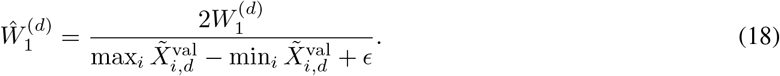

Here *ϵ* = 10^−12^ is a small constant that prevents division by zero. The final metric is the average normalized Wasserstein distance across the *D* dimensions of the comparison space (i.e., the flattened, PCA-projected, and delay-embedded point clouds):

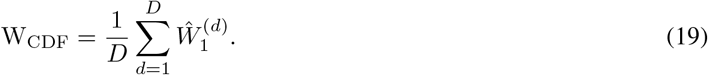

Lower values indicate that the model-predicted activity more closely matches the empirical distribution of the reference validation data in the observation-derived comparison space.

Importantly, this is not a joint multivariate optimal transport distance. Instead, it is an average of one-dimensional Wasserstein distances computed independently across dimensions. This makes the metric computationally efficient and robust in moderate-dimensional spaces, but it also means that correlations between dimensions are not directly assessed except insofar as they are reflected in the PCA and delay-embedding coordinates. Additionally, because samples are flattened across trials and time before the distributional comparison, this metric is order-agnostic: it compares the distribution of visited states or delay-embedded trajectory segments, but does not directly compare trial-by-trial temporal alignment or the underlying flow field.

### E.6 Lyapunov Spectrum

#### Jacobian Computation

For each time step *t* and sampled trajectory, we compute the Jacobian matrices:

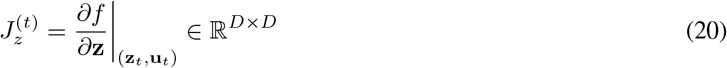

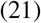

where *D* is the dimensionality of the state space. These Jacobians are computed using automatic differentiation over multiple sampled trajectories.

#### Lyapunov Exponent Estimation

The Lyapunov exponents are estimated using the QR decomposition algorithm (adapted from [67]). We initialize an orthonormal matrix *Q*_0_ = *I*_*D*_ and iteratively apply the following steps:

1. Compute the matrix product: 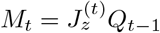
2. QR decomposition: *M*_*t*_ = *Q*_*t*_*R*_*t*_ where *Q*_*t*_ is orthonormal and *R*_*t*_ is upper triangular
3. Ensure positive diagonal elements: *R*_*t*_ ← sign(diag(*R*_*t*_)) · *R*_*t*_, *Q*_*t*_ ← *Q*_*t*_ · sign(diag(*R*_*t*_))
4. Accumulate logarithmic growth rates: 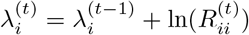

After *T* time steps, the Lyapunov exponents are estimated as:

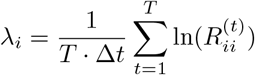

where Δ*t* is the discrete time step size.

We take the maximal Lyapunov exponent *λ*_max_ = max_*i*_ *λ*_*i*_ as our primary stability metric. This quantity governs the overall stability of the dynamical system:

- *λ*_max_ *<* 0: System is asymptotically stable (perturbations decay exponentially)
- *λ*_max_ = 0: System is marginally stable
- *λ*_max_ *>* 0: System exhibits chaotic behavior (perturbations grow exponentially)

We report the maximal Lyapunov exponent as our primary stability metric because it is the component most directly estimable from data, and therefore the one most readily compared across TT and DD models. The maximal exponent alone, however, does not fully characterize stability: the complete Lyapunov spectrum is required to distinguish, for example, a chaotic *attracting* set (positive maximal exponent but negative sum of exponents) from a divergent one. CtDToolkit therefore provides a `compute_lyapunov_spectrum` routine (and a corresponding `Analysis_TT.compute_lyapunov_spectrum` method) that returns the full ordered set of exponents {*λ*_1_ ≥ … ≥ *λ*_*D*_} together with summary statistics, including their sum ∑_*i*_ *λ*_*i*_ *(t*he mean rate of phase-space volume contraction) and the Kaplan–Yorke (Lyapunov) dimension. We validated this estimator on the canonical Lorenz Lorenz attractor, the estimated spectrum [+0.91, −0.01, −14.56] reproduces the reference [+0.906, 0, −14.572], and Rössler systems, recovering spectra that closely match their literature references (Supp. Fig. S10). For the with *λ*_max_ *>* 0 (chaos) and ∑_*i*_ *λ*_*i*_ ≈ −13.7 (an attracting set; Kaplan–Yorke dimension ≈ 2.06). For the Rössler attractor, the estimated spectrum [+0.07, +0.01, − 5.48] likewise matches the reference [+0.071, − 0, 5.394]. The worked example is provided at `examples/figures/supplementary/lyapunov_spectrum_lorenz.py`.

**Figure S10.**
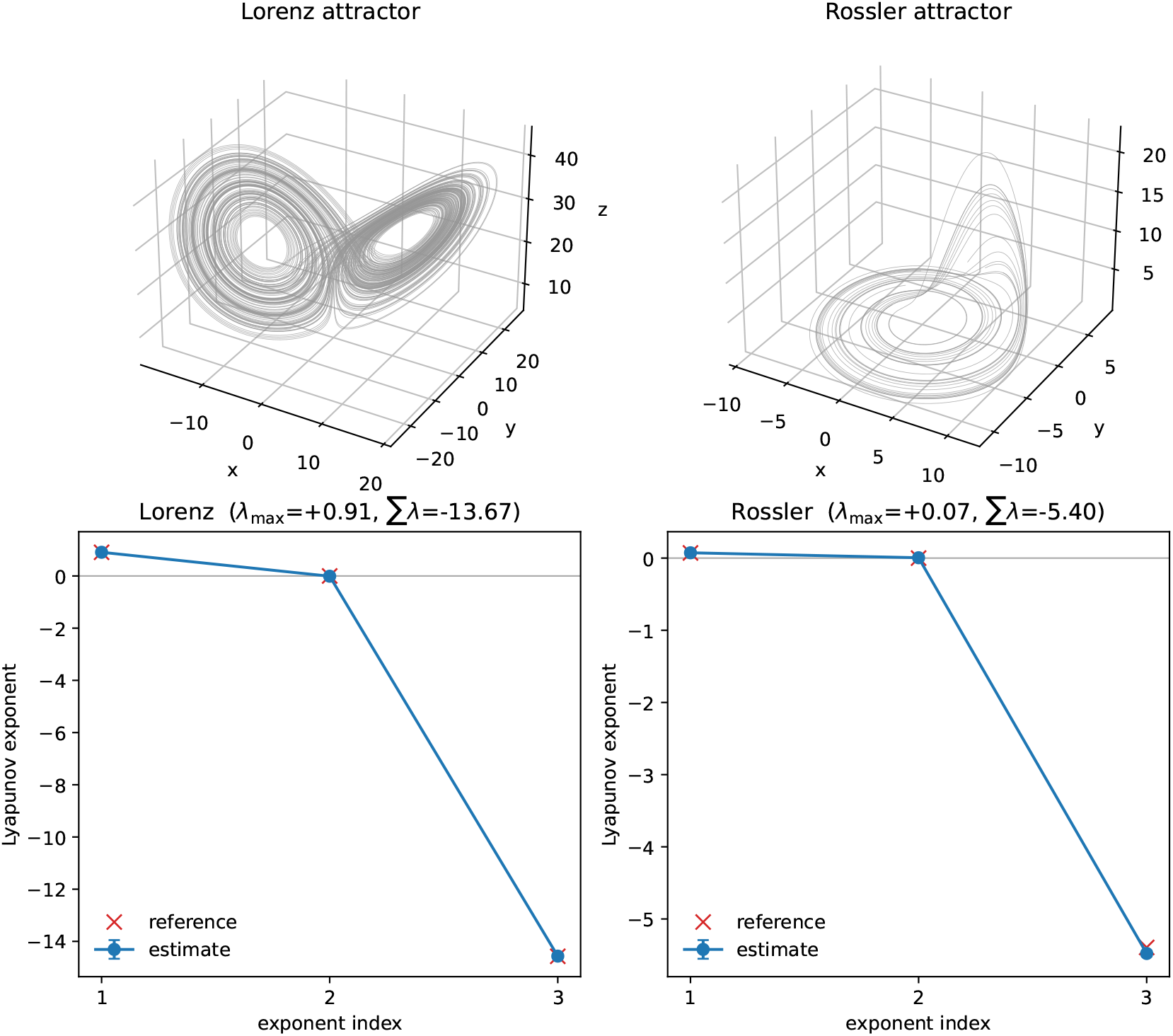
Validation of the CtDToolkit Lyapunov-spectrum estimator on canonical chaotic systems. The full Lyapunov spectrum was estimated for the standard Lorenz (*σ* = 10, *ρ* = 28, *β* = 8*/*3; left) and Rössler (*a* = 0.2, *b* = 0.2, *c* = 5.7; right) systems by integrating the vector field with a fourth-order Runge–Kutta scheme, building discrete-time Jacobians by finite differences, and applying `compute_lyapunov_spectrum` (QR-decomposition method). For each system, points and error bars show the mean ± s.d. of the estimated exponents across *B* = 4 trajectories (*T* = 120,000 RK4 steps at d*t* = 10^−3^, with a 20-time-unit burn-in); red crosses show the literature reference values. Panel titles report the maximal exponent *λ*_max_ (positive indicates chaos) and the sum ∑_*i*_ *λ*_*i*_ *(n*egative indicates an attracting set). Reproducible from `examples/figures/supplementary/lyapunov_spectrum_lorenz.py`.

### E.7 Example Performance of DD Models on CtDToolkit Datasets

To provide a demonstration of the performance metrics applied to each dataset, we replicated the experiment shown in Figure 5 for 3BFF/NBFF, MultiTask, RandomTarget, PhaseCodedMemory, and ChaoticDelayedMatching. For each dataset, we trained DD models across a sweep of latent dimensionalities, using five random seeds per latent size (NODE-based SAE models for 3BFF/NBFF, MultiTask, RandomTarget, and ChaoticDelayedMatching; LFADS models for PhaseCodedMemory). The sweeps contain seven latent sizes for 3BFF/NBFF, six latent sizes for MultiTask, and eight latent sizes each for RandomTarget, PhaseCodedMemory, and ChaoticDelayedMatching. The figures below show the completed sweeps for all five datasets. After training was complete, we used Analysis and Comparison objects to evaluate the performance of all trained models on 6 metrics: 1) Rate *R*^2^, 2) State *R*^2^, 3) co-BPS, 4) Cycle-Consistency, 5) Wasserstein distance, and 6) estimated maximal Lyapunov exponent. The model and training hyperparameters for these sweeps are compiled in Table S14 (NODE-based SAE sweeps for 3BFF/NBFF, MultiTask, RandomTarget, and ChaoticDelayedMatching) and Table S15 (LFADS sweep for PhaseCodedMemory).

**Figure S11.**
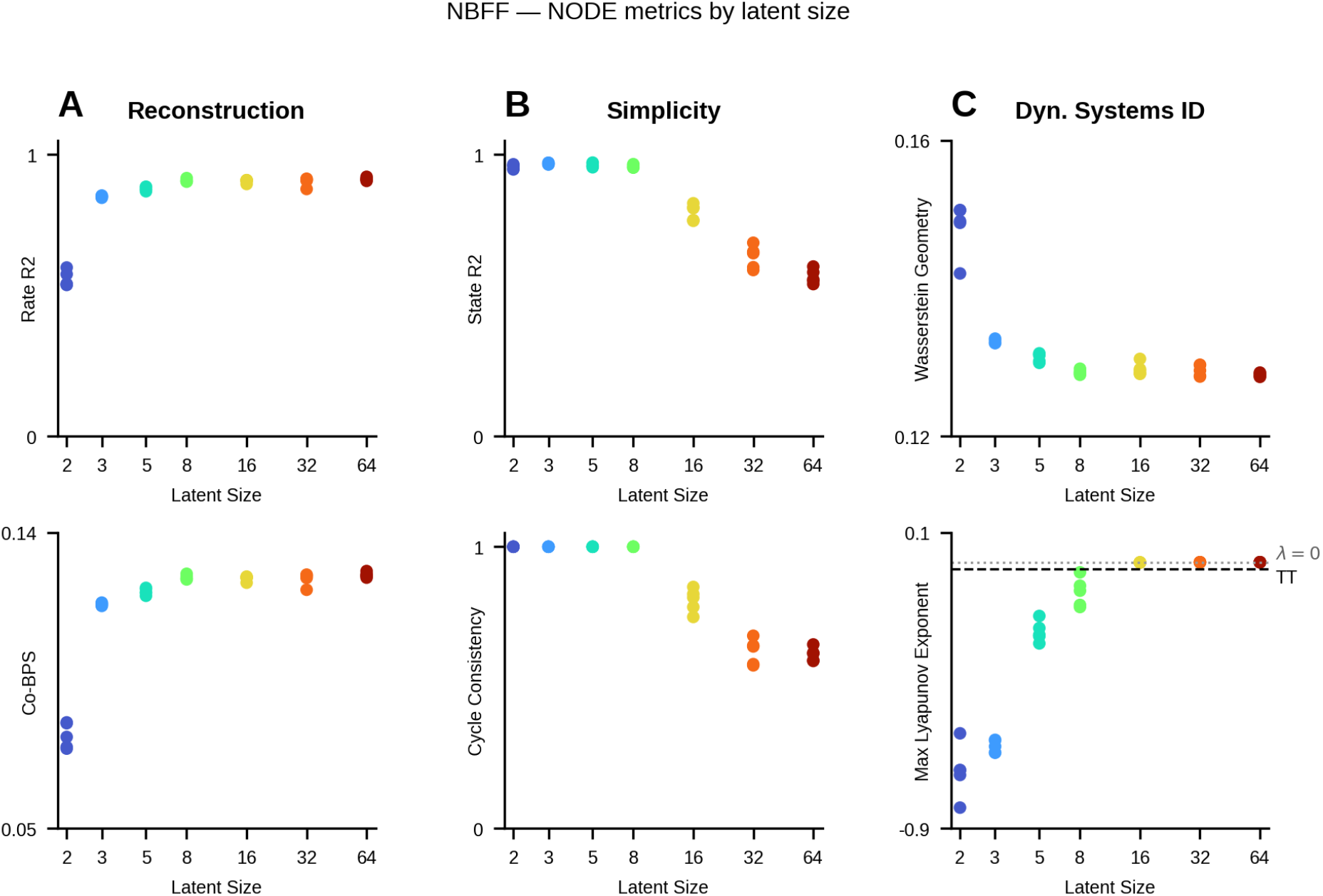
Compiled Metrics for DD-NODE models trained on NBFF Canonical Dataset: In all plots, color indicates latent size of DD model. **A)** Reconstruction metrics for DD-NODEs fit to NBFF Canonical dataset. We see similar trends for Rate *R*^2^ (top row) and co-BPS (bottom row). **B)** Simplicity metrics for models shown in panel A. We see similar trends for State *R*^2^ (top row) and Cycle-Consistency (bottom row) **C)** Metrics inspired by standard dynamical systems ID techniques applied to models presented in panel A. Top row: Wasserstein Distance. Bottom row: Estimated maximal Lyapunov exponent for each DD model. Estimated maximal Lyapunov exponent for TT network shown as dashed horizontal line.

**Figure S12.**
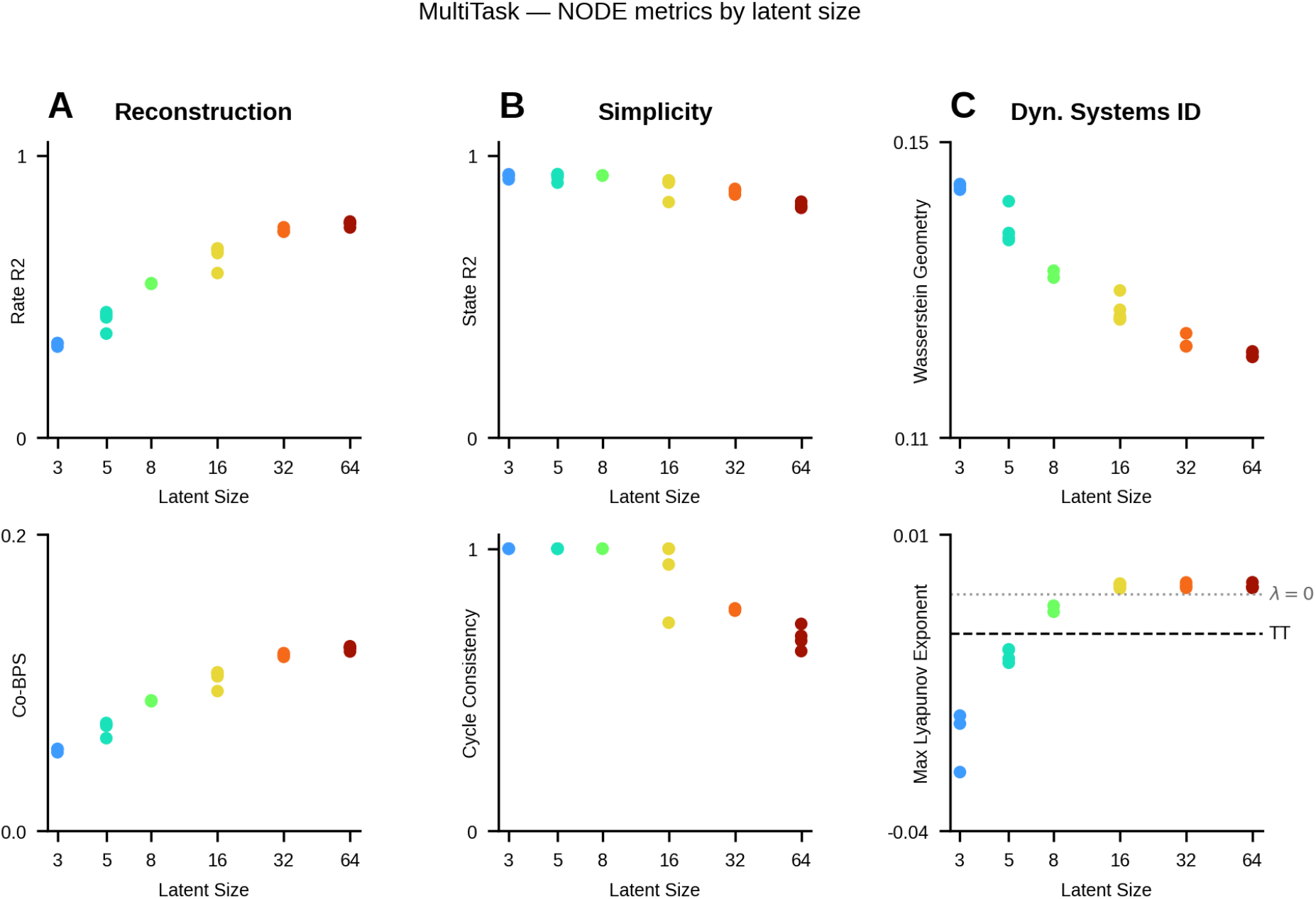
Compiled Metrics for DD-NODE models trained on MultiTask Canonical Dataset: **A)** Reconstruction metrics for DD-NODEs fit to MultiTask Canonical dataset. We see similar trends for Rate *R*^2^ (top row) and co-BPS (bottom row). **B)** Simplicity metrics for models shown in panel A. We see similar trends for State *R*^2^ (top row) and Cycle-Consistency (bottom row) **C)** Metrics inspired by standard dynamical systems ID techniques applied to models presented in panel A. Top row: Wasserstein Distance. Bottom row: Estimated maximal Lyapunov exponent for each DD model. Estimated maximal Lyapunov exponent for TT network shown as dashed horizontal line.

**Figure S13.**
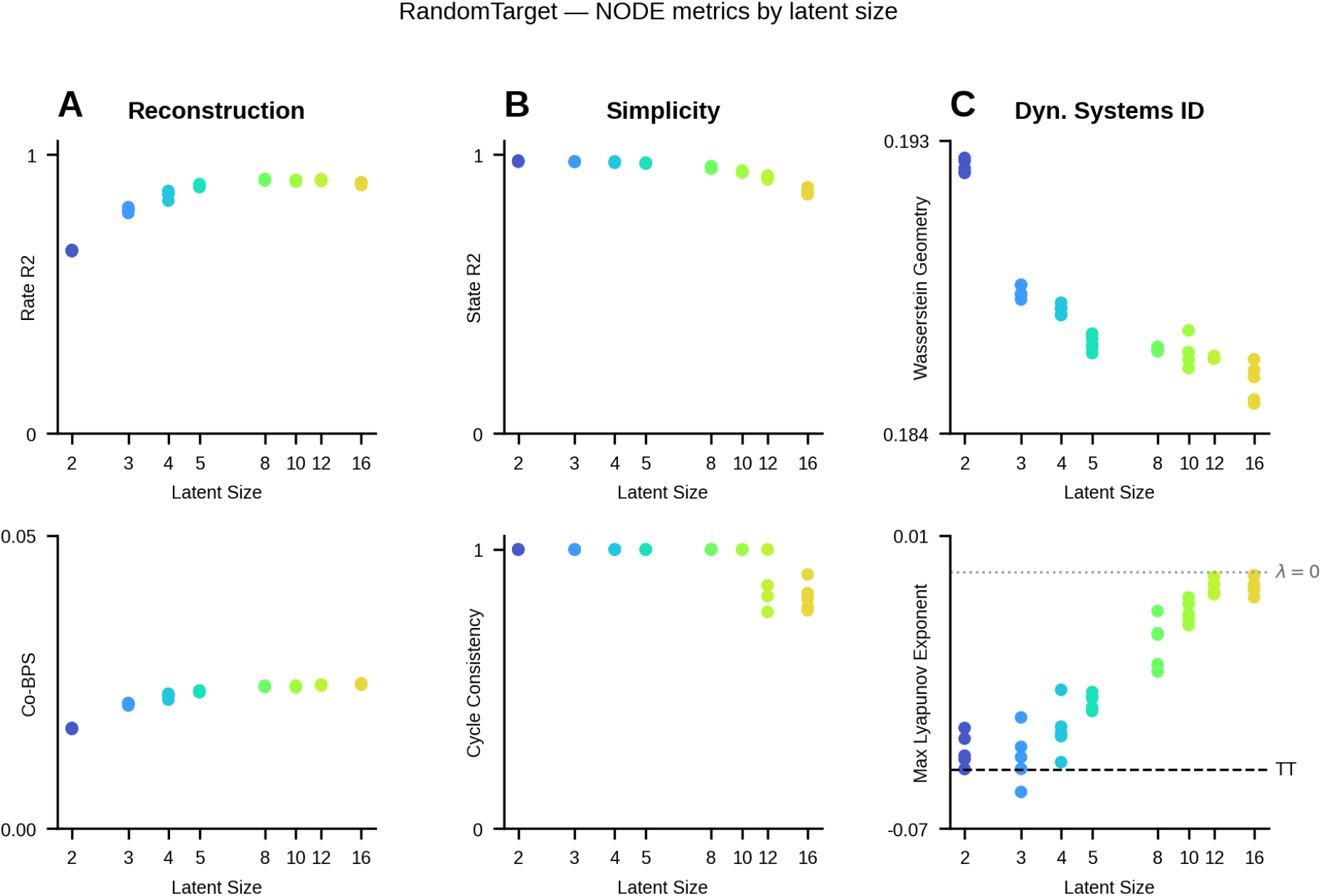
Compiled Metrics for DD-NODE models trained on RandomTarget Canonical Dataset: **A)** Reconstruction metrics for DD-NODEs fit to RandomTarget Canonical dataset. We see similar trends for Rate *R*^2^ (top row) and co-BPS (bottom row). Color indicates the latent size (x-axis) **B)** Simplicity metrics for models shown in panel A. We see similar trends for State *R*^2^ (top row) and Cycle-Consistency (bottom row) **C)** Metrics inspired by standard dynamical systems ID techniques applied to models presented in panel A. Top row: Wasserstein distance. Bottom row: Estimated maximal Lyapunov exponent for each DD model. Estimated maximal Lyapunov exponent for TT network shown as dashed horizontal line.

**Figure S14.**
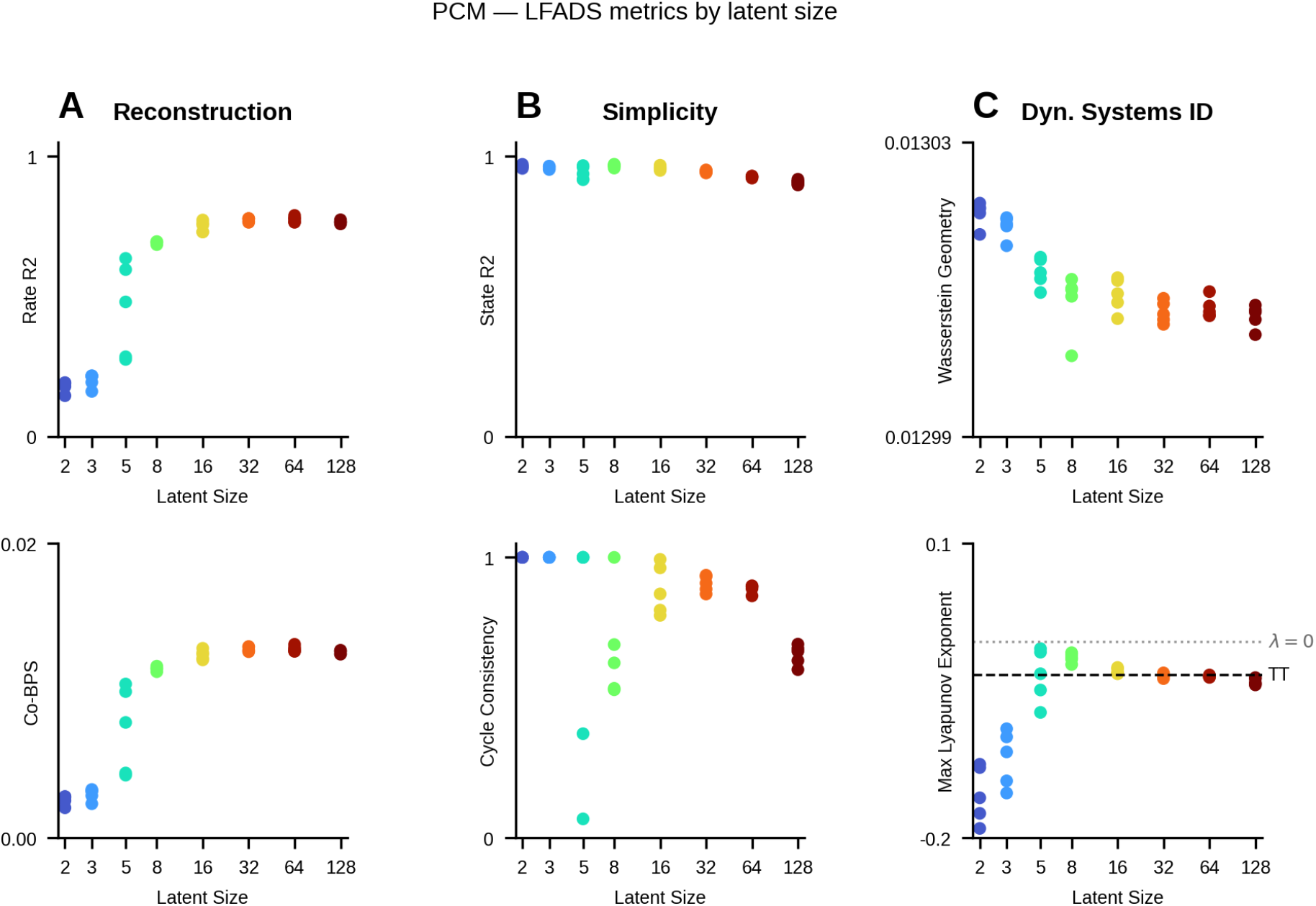
Compiled Metrics for LFADS models trained on the PhaseCodedMemory Canonical Dataset: In all plots, color indicates the latent size of the DD model. **A)** Reconstruction metrics for LFADS models fit to the PhaseCodedMemory Canonical dataset: Rate *R*^2^ (top row) and co-BPS (bottom row). **B)** Simplicity metrics for the models shown in panel A: State *R*^2^ (top row) and Cycle-Consistency (bottom row). **C)** Metrics inspired by standard dynamical systems ID techniques applied to the models in panel A. Top row: Wasserstein distance for each DD model. Bottom row: Estimated maximal Lyapunov exponent for each DD model, with the estimate for the TT network shown as a dashed horizontal line and the *λ* = 0 reference (boundary between contracting and expanding dynamics) shown as a dotted line. As latent size increases, the LFADS estimates converge toward the TT value.

**Figure S15.**
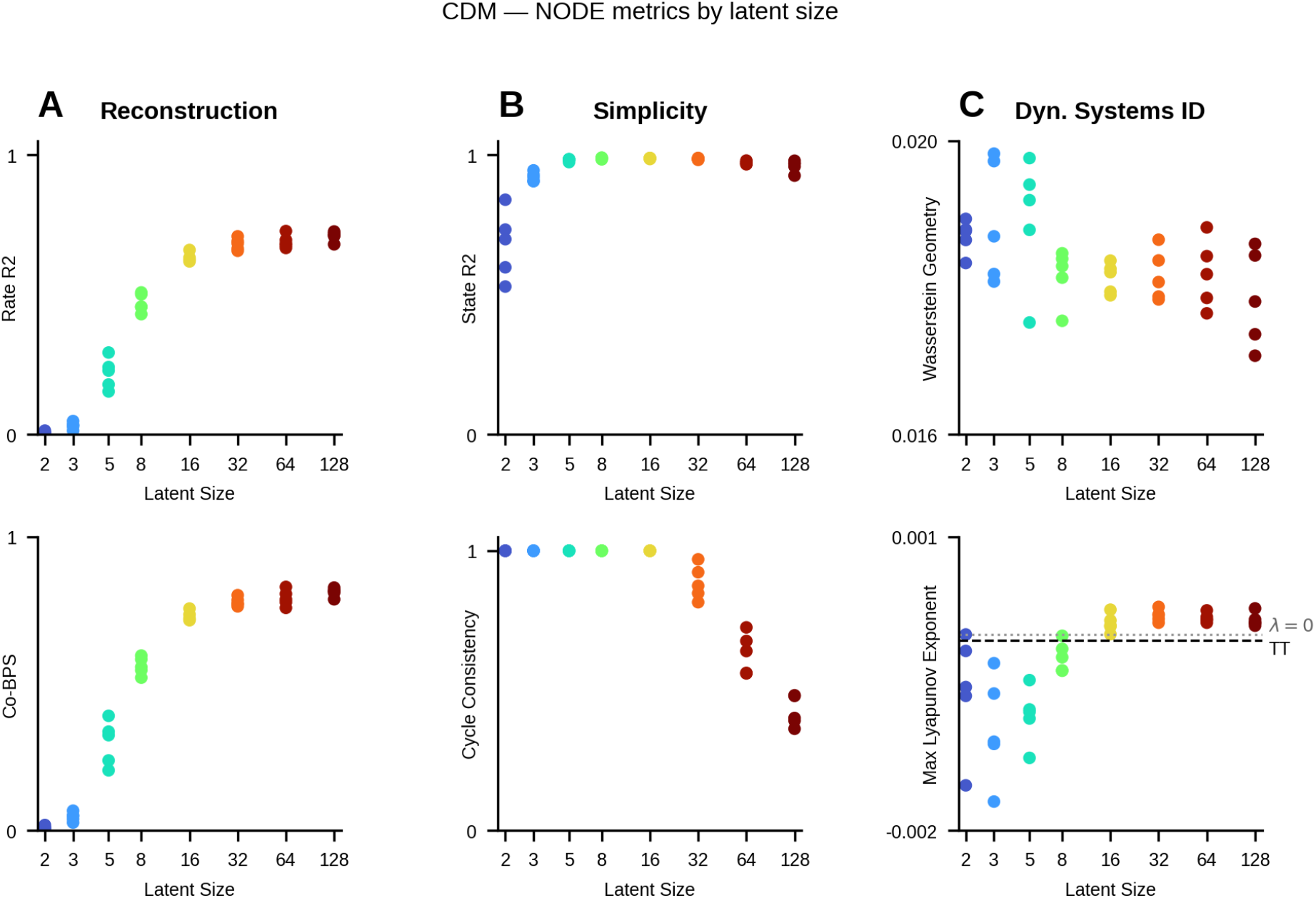
Compiled Metrics for DD-NODE models trained on ChaoticDelayedMatching Canonical Dataset: In all plots, color indicates latent size of DD model. **A)** Reconstruction metrics for DD-NODEs fit to ChaoticDelayedMatching Canonical dataset. We see similar trends for Rate *R*^2^ (top row) and co-BPS (bottom row). **B)** Simplicity metrics for models shown in panel A. We see similar trends for State *R*^2^ (top row) and Cycle-Consistency (bottom row). **C)** Metrics inspired by standard dynamical systems ID techniques applied to models presented in panel A. Top row: Wasserstein distance. Bottom row: Estimated maximal Lyapunov exponent for each DD model. Estimated maximal Lyapunov exponent for TT network shown as dashed horizontal line.

**Table S14.** Model and training hyperparameters for the NODE-based SAE latent-size sweeps used in the compiled-metrics figures (Supp. Figs. S11, S12, S13, S15). Each model is trained at every listed latent size 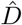 with 5 random seeds (0–4). The vector field of the Neural ODE is an MLP with `vf_num_layers` layers of `vf_hidden_size` units; `encoder_size` is the hidden size of the bidirectional GRU initial-condition encoder. All models use the Adam optimizer with a gradient-clip value of 0.01 and are supplied ground-truth external inputs. ^†^MultiTask uses a task-masked Poisson loss (`MultiTaskPoissonLossFunc`).

| Hyperparameter | NBFF | MultiTask | RandomTarget | CDM |
| --- | --- | --- | --- | --- |
| latent sizes ($\hat{D}$) | 2,3,5,8,16,32,64 | 3,5,8,16,32,64 | 2,3,4,5,8,10,12,16 | 2,3,5,8,16,32,64,128 |
| seeds per size | 5 | 5 | 5 | 5 |
| input_size | 3 | 20 | 17 | 2 |
| encoder_size | 100 | 100 | 100 | 100 |
| vf_hidden_size | 128 | 128 | 128 | 128 |
| vf_num_layers | 6 | 6 | 2 | 6 |
| dropout | 0.05 | 0.05 | 0.05 | 0.05 |
| learning_rate | 2e-3 | 2e-3 | 2e-3 | 1e-3 |
| weight_decay | 0 | 1e-7 | 1e-6 | 0 |
| batch_size | 250 | 250 | 250 | 512 |
| grad_clip | 0.01 | 0.01 | 0.01 | 0.01 |
| epochs | 1000 | 500 | 350 | 200 |
| optimizer | Adam | Adam | Adam | Adam |
| recon. loss | Poisson | Poisson<sup>†</sup> | Poisson | Poisson |

**Table S15.** Model and training hyperparameters for the LFADS latent-size sweep on the PhaseCodedMemory dataset used in the compiled-metrics figure (Supp. Fig. S14). The generator (latent) dimension `gen_dim` is swept across the listed sizes with 5 random seeds (0–4) each, while the initial-condition encoder and factor dimensions are held fixed at 128. Because ground-truth external inputs are supplied, the controller path is disabled (`ci_enc_dim`=`con_dim`=`co_dim`=0), so the controller-input prior and the `kl_co` term are inactive. KL and L2 penalties are linearly ramped from their `*_start_epoch` to `*_increase_epoch`. The low firing-rate cold-start flags (`rate_bias_init`, `rate_weight_init_scale`) are described in Appendix D.3.2.

| Hyperparameter | LFADS (PhaseCodedMemory) |
| --- | --- |
| gen_type | RNN |
| gen_dim ($\hat{D}$, swept) | 2,3,5,8,16,32,64,128 |
| seeds per size | 5 |
| ic_encoder_size (ic_enc_dim) | 128 |
| ic_dim | 128 |
| fac_dim | 128 |
| ext_input_dim | 3 |
| ci_enc_dim / con_dim / co_dim | 0 / 0 / 0 |
| cd_rate | 0.3 |
| ic_prior.mean | 0 |
| ic_prior.variance | 0.1 |
| ic_post_var_min | 1.0e-4 |
| dropout | 0.2 |
| cell_clip | 5.0 |
| loss_scale | 1.0e+4 |
| rate_bias_init | True |
| rate_weight_init_scale | 0.1 |
| lr_init | 5.0e-3 |
| lr_stop | 1.0e-5 |
| lr_decay | 0.95 |
| lr_patience | 6 |
| lr_adam_beta1 / beta2 / epsilon | 0.9 / 0.999 / 1.0e-7 |
| weight_decay | 1.0e-8 |
| l2_gen_scale | 1.0e-5 |
| l2_start / increase_epoch | 0 / 80 |
| kl_ic_scale | 1.0e-3 |
| kl_start / increase_epoch | 0 / 80 |
| batch_size | 1000 |
| epochs | 200 |
| reconstruction | Poisson (variational) |

### E.8 DD Model Validation on PhaseCodedMemory

To complement the canonical metric sweeps above, we examine an example data-driven (DD) model fit on the cyclic PhaseCodedMemory dataset. This example illustrates how CtDToolkit metrics and visualizations can be used to assess whether a DD model has recovered the correct dynamical strategy of the underlying TT system.

#### PhaseCodedMemory: LFADS recovers a rate-coding rather than phase-coding solution

We trained LFADS models across a range of latent dimensionalities (8D, 16D, and 128D) on the canonical PhaseCodedMemory dataset (Supp. Fig. S4) and compared the inferred firing rates and latent activity against the ground-truth TT model (Supp. Fig. S16). Larger models achieved better reconstruction of the simulated firing rates (Rate *R*^2^ increasing from 0.68 at 8D to 0.77 at 128D), but at the cost of a degrading State *R*^2^ (decreasing from 0.97 to 0.92), indicating that their inferred latents drift further from the ground-truth TT latent structure even as they fit the observed rates more closely. Critically, at every latent size the inferred dynamics failed to capture the qualitative phase-coding strategy of the TT model: whereas the ground-truth Stim A and Stim B trajectories form overlapping, rotated rings in both the rate-space and latent-space PCs, the DD models inferred latents in which the two stimulus conditions are largely *separated*, consistent with a rate-coding rather than phase-coding strategy. This demonstrates how DD-inferred dynamics, when interpreted without reference to ground truth, can lead to qualitatively incorrect conclusions about the underlying neural computation, even for models that reconstruct the observed activity well. We additionally swept LFADS models across latent dimensionalities on this dataset and compiled the full suite of CtDToolkit metrics as a function of latent size (Supp. Fig. S14).

**Figure S16.**
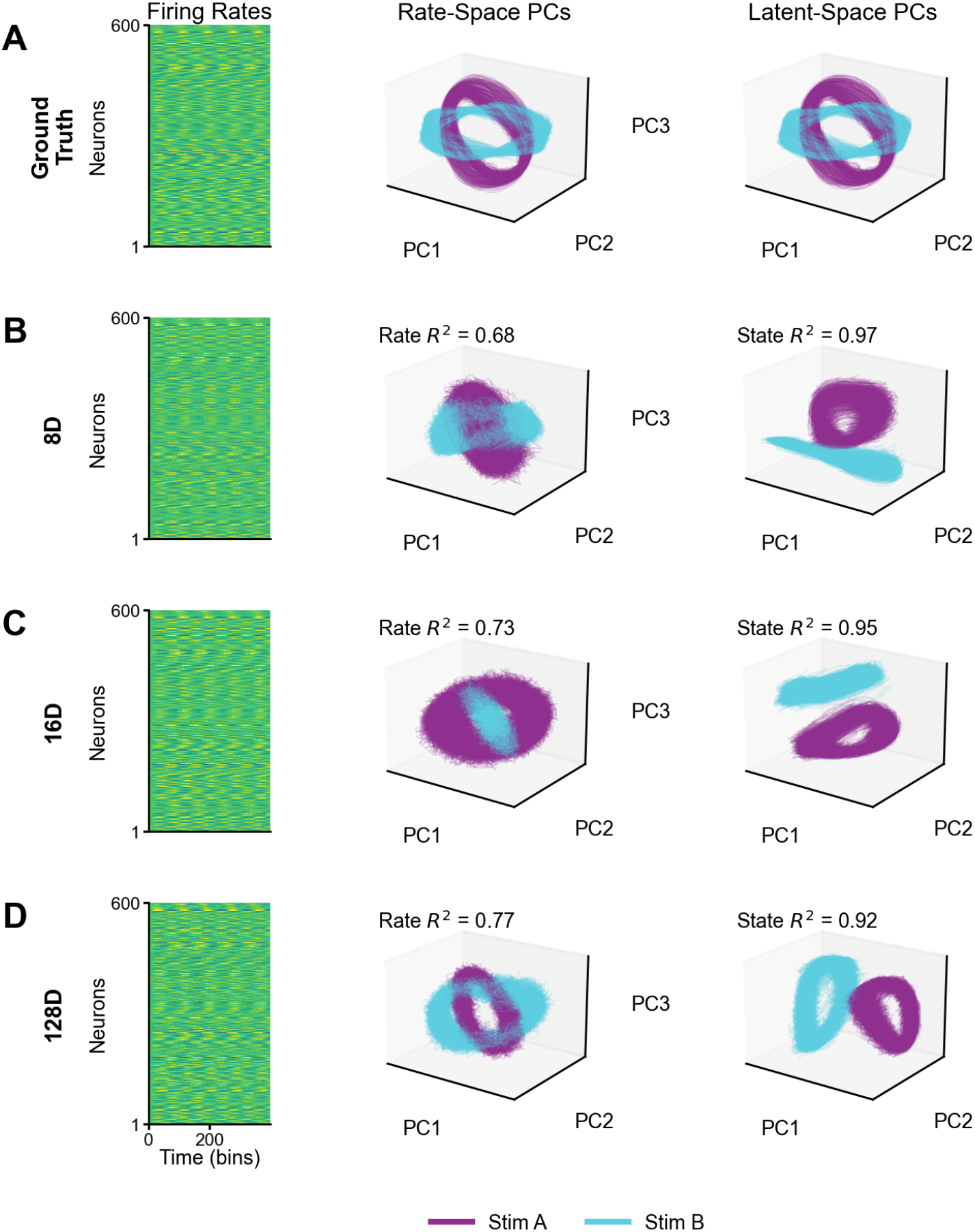
Data-Driven Model Validation on PhaseCodedMemory Each row corresponds to one model: **A**) the ground-truth TT system and (**B**-**D**) LFADS DD models of increasing latent dimensionality (**B**: 8D, **C**: 16D, **D**: 128D). Within each row, column 1 shows an imagesc raster of the firing rates of the 500 held-in simulated neurons for an example trial (ground-truth rates in row A; DD-inferred rates in the model rows); column 2 shows the top 3 PCs of the response-period activity in *firing-rate* space, with trajectories colored by stimulus condition (Stim A, purple; Stim B, cyan). The annotated Rate *R*^2^ is the variance-weighted reconstruction between the ground-truth and DD-inferred firing rates. Column 3 shows the top 3 PCs of the response-period activity in *latent* space (TT latents in row A; DD-inferred latents in the model rows). The annotated State *R*^2^ is computed from a linear map between the TT and DD latents. In the ground-truth TT model, the Stim A and Stim B rings overlap and are primarily rotated relative to one another in both the rate- and latent-space PCs, reflecting a phase-coding strategy (cf. Supp. Fig. S4C). Reconstruction (Rate *R*^2^) improves with latent size while State *R*^2^ degrades, and at no latent size do the DD models recover the qualitative phase-coding geometry: the inferred latents instead separate the stimulus conditions, consistent with an incorrect rate-coding solution. This highlights the danger of interpreting DD-inferred dynamics without guidance from the CtDToolkit metrics.

## F Utilizing the Task-Training Pipeline

We used PyTorch Lightning to modularize the code-base to allow for easy additions and extensions from users. The CtDToolkit pipeline defines five objects designed to simplify the experience for the user. These objects are:

1. TaskEnvironment: The task environment (described in Section F.1, schematic in Fig. S17), contains the task logic for a given task. TaskEnvironment inherits from Gym.Environment [60], meaning that new TaskEnvironments must implement the `step()` and `reset()` functions. Additionally, it must implement the `generate_dataset()` function, which generates a dictionary of inputs and desired outputs for the task-trained model.
2. TT model: Inherits from nn.Module [30]. This object (schematic in Fig. S17) defines the model that will be trained to perform the task. This object must implement the `forward()` method, which takes in the input and the hidden state at time *t* to output the hidden state at time *t* + 1. Additionally, this object must implement the `init_model()` method and (optionally), the `init_hidden()` method.
3. TaskWrapper: Inherits from LightningModule [27]. This object takes in the TaskEnvironment and the TT model and handles the training and validation loops. This object should not need to be modified by the typical user.
4. TTDatamodule: Inherits from LightningDataModule [27]. This object handles the data generation from the TaskEnvironment and batching during training and validation. This object should not need to be modified by the typical user.
5. Simulator: This object handles the generation of synthetic neural activity from the trained TT model. The `_ _init_ _ ()` method of this object takes in two dictionaries, `noise_dict` and `embed_dict` (the simulator configuration name for observation-model settings), which provide settings for the noise model used to sample spiking events from the simulated firing rates and the nonlinear rectification/observation model of the task-trained model’s hidden unit activity, respectively. The Simulator object should not need to be modified by the typical user. We provide more detail on the Simulator in Section C.

**Figure S17.**
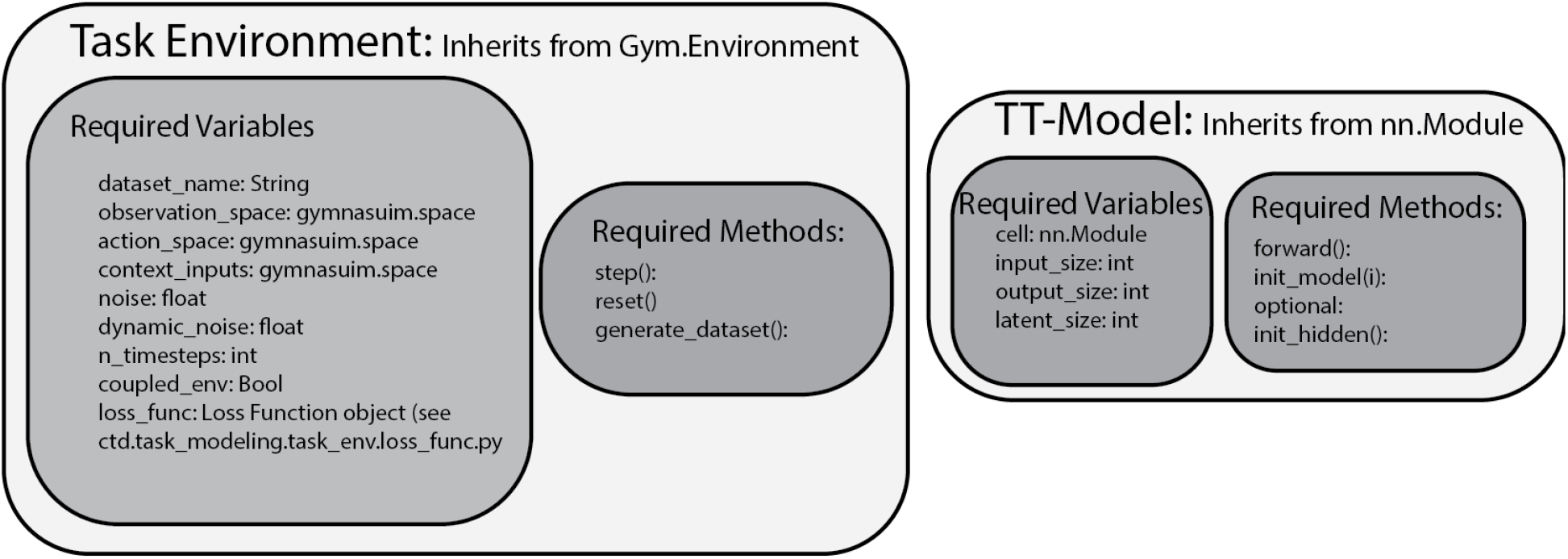
Objects for user-defined tasks and TT models.

### F.1 Task Environment Requirements

Our task environments adhere to a general format that is modular and easily extensible, allowing researchers to add new tasks to CtDToolkit without major code restructuring. TaskEnvironments are subclasses of Gymnasium Gym environments [60]. In total, new task environments must meet the following requirements:

#### Required Variables

- `dataset_name`: *string*. Used as an identifier when saving and loading datasets in the task-training and data-training pipelines.

#### Input/Outputs

These variables define the input/output spaces of the task. All are built using Gymnasium Space objects.

- `action_space`: *Gymnasium Space*. Defines the allowable range of actions to be taken in the environment. `3BFF`, `PhaseCodedMemory`, `MultiTask`, and `RandomTarget` have 3,1, 3, and 6, respectively.
- `observation_space`: *Gymnasium Space*. Defines the allowable range of “sensory” inputs that the environment provides. `3BFF`, `PhaseCodedMemory`, `MultiTask`, and `RandomTarget` have 3,3, 20, and 14, respectively.
- `context_inputs`: *Gymnasium Space*. Defines the allowable range of “context” inputs to the model. Only `RandomTarget` currently has non-zero context inputs (3).

#### Additional Variables

- `noise`: *float*. Describes the standard deviation of Gaussian noise to be added to the inputs in the pre-generated dataset.
- `n_timesteps`: *int*. Provides the trial length for the TaskEnvironment (in bins).
- `coupled_env`: *bool*. Indicates whether the actions need to be provided to the environment to generate the next step of sensory inputs (i.e., if the controller and environment are coupled). Only `True` for `RandomTarget` for canonical datasets.
- `loss_func`: A CtDToolkit class that takes in a `loss_dict` and computes the loss for the environment. See `/ctd/task_modeling/task_env/loss_func.py` for implementations of each `TaskEnvironment` loss.

#### Required Methods

- `step(self, action)`: Inherited from Gym. Should implement the state update for a given `TaskEnvironment`.
- `reset(self)`: Inherited from Gym. Should reset required variables to restart trial (e.g., set initial states, etc.).
- `generate_dataset(self, n_samples)`: Defines how many trials should be generated from the dataset; for `MultiTask`, `n_samples` defines the number of trials to be generated for each sub-task. `generate_dataset` should return a dictionary the following fields given in Section B.1.

We provide a template TaskEnvironment in `/ctd/task_modeling/task_env/task_env.py`

### F.2 How to Submit a CtDToolkit Dataset

1. **Fork the repository**. Create a personal fork of the `CtDToolkit` GitHub repository.
2. **Implement a new task environment**. In your fork, add a new task environment that conforms to the Toolkit’s task-environment API (e.g., initialization, reset, step, observation, reward functions), described above.
3. **Train a task-trained model**. Train a model to perform the task within your new environment. Save both (i) the trained model and (ii) a simulator object capable of generating simulated spiking activity. To use the Analysis and Comparison object methods, the TT model must adhere to the template provided in `/ctd/task_modeling/model/tt_template.py`
4. **Validate model performance**. Demonstrate that the trained model can *adequately perform the task* (e.g., achieving appropriate behavioral performance or reward thresholds) and produces realistic neural activity (e.g., firing-rate statistics or spiking patterns). Summarize these results for review.
5. **Register the dataset**. Modify the `gen_datasets.py` script to add your dataset, linking it to the saved model and simulator object. This ensures your dataset is integrated into the Toolkit’s dataset generation pipeline.
6. **(Recommended) Add documentation**. Provide a short “dataset card” including a description of the task, training details, validation results, and an example usage snippet.
7. **Submit a pull request**. Open a pull request (PR) to the main branch of `CtDToolkit`, briefly describing the task, what makes the dataset novel or useful, and including your validation summary.
8. **Review and merge**. The maintainers will review the PR to ensure correctness, novelty, and scientific utility. Once approved, the PR is merged, and the dataset becomes available as part of the official `CtDToolkit` test-set collection.

## G Compute Resources

We used an internal computing cluster with a total of 30 Nvidia GeForce RTX 2080 Ti GPUs for model training. The task-training pipeline took 0.5, 8, and 4 hours to train for the 3BFF, MultiTask, and RandomTarget models, respectively. The RandomTarget task does not parallelize, due to its serial nature, so we did not use GPUs to train that model. For the data-driven models, both the SAE and LFADS models each took less than 1 hour to train; we trained a total of 95 data-driven models across all figures. With 2 models training on each GPU, all experiments took <150 GPU-hours. FP finding was fast, requiring 1 minute for each model.

## H Open-source Packages Used

- `torch` [30] (BSD license)
- `pytorch-lightning` [27] (Apache 2.0 license)
- `ray.tune` [24] (Apache 2.0 license)
- `lfads-torch` [58] (Apache 2.0 license)
- `hydra` [32] (MIT license)

